# Chromosome-scale genome assembly provides insights into rye biology, evolution, and agronomic potential

**DOI:** 10.1101/2019.12.11.869693

**Authors:** M. Timothy Rabanus-Wallace, Bernd Hackauf, Martin Mascher, Thomas Lux, Thomas Wicker, Heidrun Gundlach, Mariana Báez, Andreas Houben, Klaus F.X. Mayer, Liangliang Guo, Jesse Poland, Curtis J. Pozniak, Sean Walkowiak, Joanna Melonek, Coraline Praz, Mona Schreiber, Hikmet Budak, Matthias Heuberger, Burkhard Steuernagel, Brande Wulff, Andreas Börner, Brook Byrns, Jana Čížková, D. Brian Fowler, Allan Fritz, Axel Himmelbach, Gemy Kaithakottil, Jens Keilwagen, Beat Keller, David Konkin, Jamie Larsen, Qiang Li, Beata Myśków, Sudharsan Padmarasu, Nidhi Rawat, Uğur Sesiz, Biyiklioglu Sezgi, Andy Sharpe, Hana Šimková, Ian Small, David Swarbreck, Helena Toegelová, Natalia Tsvetkova, Anatoly V. Voylokov, Jan Vrána, Eva Bauer, Hanna Bolibok-Bragoszewska, Jaroslav Doležel, Anthony Hall, Jizeng Jia, Viktor Korzun, André Laroche, Xue-Feng Ma, Frank Ordon, Hakan Özkan, Monika Rakoczy-Trojanowska, Uwe Scholz, Alan H. Schulman, Dörthe Siekmann, Stefan Stojałowski, Vijay Tiwari, Manuel Spannagl, Nils Stein

**Affiliations:** Harrow Research and Development Centre, Agriculture and Agri-Food Canada, 2585 County Road 20, Harrow, ON, N0R 1G0, Canada; Lethbridge Research and Development Centre, Agriculture and Agri-Food Canada, 5403 1st Avenue South, Lethbridge AB T1J 4B1, Canada; Chinese Academy of Crop Sciences (CAAS), No.12 Zhongguancun South Street, Haidian District, Beijing 100081, China; Plant Genomics, Earlham Institute, Norwich Research Park, Norwich, Norfolk, NR4 7UG, UK; Department of Botany, Federal University of Pernambuco, Av. Prof. Moraes Rego, 1235, Cidade Universitária, Recife PE, 50670-901, Brazil; Plant Genome and Systems Biology (PGSB), Helmholtz Zentrum München, Ingolstädter Landstr. 1, 85764 Neuherberg, Germany; National Key Laboratory of Crop Genetic Improvement, Huazhong Agricultural University, No. 1 Shizishan Street, Hongshan District, Wuhan, Hubei Province, China; HYBRO Saatzucht GmbH & Co. KG, Langlinger Str. 3, 29565 Wriedel, Germany; Institute of Experimental Botany, Czech Academy of Sciences, Centre of the Region Hana for Biotechnological and Agricultural Research, Šlechtitelů 31, 779 00 Olomouc, Czech Republic; Computational Systems Biology, John Innes Centre, Norwich Research Park, Norwich, NR4 7UH, UK; Crop Genetics, John Innes Centre, Norwich Research Park, Norwich, NR4 7UH, UK; Institute for Biosafety in Plant Biotechnology, Julius Kühn-Institute, Erwin-Baur-Str. 27, 06484 Quedlinburg, Germany; Institute for Breeding Research on Agricultural Crops, Julius Kühn-Institute, Rudolf-Schick-Platz 3a, 18190 Groß Lüsewitz, Germany; Institute for Resistance Research and Stress Tolerance, Julius Kühn-Institute, Erwin-Baur-Str. 27, 06484 Quedlinburg, Germany; KWS SAAT SE & Co. KGaA, Grimsehlstr. 31, 37574 Einbeck, Germany; Federal State Budgetary Institution of Science Federal Research Center, Kazan Scientific Center of Russian Academy of Sciences, ul. Lobachevskogo, 2/31, Kazan 420111, Tatarstan, Russian Federation; Leibniz Institute of Plant Genetics and Crop Plant Research (IPK), Corrensstr. 3, 06466 Stadt Seeland, Germany; Department of Crop Sciences CiBreed - Center for Integrated Breeding Research, Georg-August University Göttingen, Von Siebold Straße 8, D-37075 Göttingen, Germany; German Centre for Integrative Biodiversity Research (iDiv) Halle-Jena-Leipzig, Leipzig, Germany; Montana BioAgriculture Inc., Montana, USA; Kansas State University, 4024 Throckmorton Hall, Kansas State University, Manhattan, KS 66506, USA; Production Systems, Natural Resources Institute Finland (Luke), Latokartanonkaari 9, 00790 Helsinki, Finland; Noble Research Institute, LLC, 2510 Sam Noble Parkway, Ardmore, OK 73401, USA; Vavilov Institute of General Genetics, Russian Academy of Sciences, Gubkina 3, 119991 Moscow, Russia; Molecular Biology, Genetics and Bioengineering, Sabanci University, University Cad No 27, Istanbul, Turkey; Department of Genetics and Biotechnology, Saint Petersburg State University, Universitetskaya emb. 7/9, 199034, St. Petersburg, Russia; Plant Breeding, Technical University of Munich, Liesel-Beckmann-Str. 2, 80333 München, Germany; ARC Centre of Excellence in Plant Energy Biology, School of Molecular Sciences, The University of Western Australia, 35 Stirling Highway, Crawley 6009 WA, Australia; Department of Field Crops, University of Cukurova Faculty of Agriculture, Balcalı, Çukurova Üniversitesi Rektörlüğü, 01330 Sarıçam/Adana, Turkey; Plant Sciences and Landscape Architecture, University of Maryland, College Park, 4291 Fieldhouse Drive 2102, Plant Sciences Building, College Park, MD 20742, USA; Crop Development Centre, University of Saskatchewan, 51 Campus Drive, Saskatoon, Saskatchewan S7N 5A8, Canada; Department of Plant Science, University of Saskatchewan, 51 Campus Drive, Saskatoon, Saskatchewan S7N 5A8, Canada; University of Saskatchewan, Global Institute for Food Security, 110 Gymnasium Place, Saskatoon, SK, S7N 0W9, Canada; Aquatic and Crop Resource Development, National Research Council Canada, 110 Gymnasium Pl, Saskatoon, SK S7N 0W9, Canada; Department of Plant and Microbial Biology, University of Zürich, Zollikerstrasse 107, 8008 Zürich, Switzerland; Department of Biology, ETH Zürich, Wolfgang-Pauli-Strasse 27, 8093 Zürich, Switzerland; Department of Plant Genetics Breeding and Biotechnology, Warsaw University of Life Sciences - SGGW, Nowoursynowska Str 159, 02-776 Warsaw, Poland; Department of Genetics, Plant Breeding and Biotechnology, West Pomeranian University of Technology Szczecin, Słowackiego 17, 71-434 Szczecin, Poland

## Abstract

We present a chromosome-scale annotated assembly of the rye (*Secale cereale* L. inbred line ‘Lo7’) genome, which we use to explore Triticeae genomic evolution, and rye’s superior disease and stress tolerance. The rye genome shares chromosome-level organization with other Triticeae cereals, but exhibits unique retrotransposon dynamics and structural features. Crop improvement in rye, as well as in wheat and triticale, will profit from investigations of rye gene families implicated in pathogen resistance, low temperature tolerance, and fertility control systems for hybrid breeding. We show that rye introgressions in wheat breeding panels can be characterised in high-throughput to predict the yield effects and trade-offs of rye chromatin.

## Main Text

Rye (*Secale cereale L.*) is a member of the grass tribe Triticeae and close relative of wheat (*Triticum aestivum L.*) and barley (*Hordeum vulgare L.*), grown primarily for human consumption and animal feed. Rye is uniquely tolerant of biotic and abiotic stresses and thus exhibits high yield potential under marginal conditions. This makes rye an important crop along the northern boreal-hemiboreal belt, a climatic zone predicted to expand considerably in Eurasia and North America with anthropogenic global warming^1^. Rye chromatin introgressions into bread wheat can significantly increase yield by conferring disease resistance and enhanced root biomass^2-5^. Rye also possesses a unique bi-factorial self- incompatibility system^6^, and rye genes controlling self-compatibility and male fertility have enabled the establishment of efficient cytoplasmic male sterility (CMS)-based hybrid breeding systems that exploit heterosis at large scales^7^. Implementation of such systems in cereals will be invaluable to meeting future human calorific requirements.

Rye is diploid with a large genome (∼7—8 Gbp)^8^ compared even to the diploid barley genome and the subgenomes of the hexaploid bread wheat^9^. Like barley and wheat, rye entered the genomics era very recently. A virtual gene-order was released in 2013^10^, and a shotgun *de novo* genome survey of the same line became available in 2017^11^. Both resources have been rapidly adopted by researchers and breeders^12-14^, but cannot offer the same opportunities as the higher quality genome assemblies available for other Triticeae species^9, 15-19^.

We report the assembly of a chromosome-scale genome sequence for rye line ‘Lo7’, providing insights into rye genome organisation and evolution, and representing a comprehensive resource for genomics-assisted crop improvement.

## Results

### An annotated chromosome-scale genome assembly

We estimated the genome size of 15 rye genotypes by flow-cytometry (Methods, Note S-FLOWCYT) and found ‘Lo7’ among the smaller of these at 7.9 Gbp. We *de novo* assembled scaffolds representing 6.74 Gbp of the ‘Lo7’ genome (Table 1) from >1.8 Tbp of short read sequence (Methods; Notes S-PSASS, S- ASSDATA). The scaffolds were ordered, oriented and curated using a variety of independent data sources including: (i) chromosome-specific shotgun (CSS) reads^10^, (ii) 10X Chromium linked reads, (iii) genetic map markers^11^, (iv) 3D chromosome conformation capture sequencing (Hi-C)^20^, and (v) a Bionano optical genome map (tbls. S-ASSSTATS—S-OPTSTAT). After intensive manual curation, 83% of this assembled sequence (i.e. ∼75.5% of the total genome size) was arranged first into super-scaffolds (N50 >29 Mbp) and then into pseudomolecules. Annotation of various features (Methods) yielded 34,441 high confidence genes, which we estimate comprises 97.9% of the entire gene complement (tbl. S-ANNOTSTAT), 19,456 full-length DNA LTR retrotransposons (LTR-RTs) from six transposon families (tbl. S-TEANNOT)^21^, 13,238 putative miRNAs in 90 miRNA families (tbls. S-miRNA_sequences—S-miRNA target_table), and 1,382,323 tandem repeat arrays (tbls. S-TANDREPCOMPN-S-SAT_ANNOT). Fluorescence *in situ* hybridisation (FISH) to mitotic ‘Lo7’ chromosomes using probes targeting tandem repeats showed that scaffolds for which assignment to a chromosome pseudomolecule was difficult are highly enriched in short repeats (Methods; Note S-REP).

**Table M-STATS.** Genome assembly and annotation statistics. CSS=Chromosome Specific Shotgun. BUSCO=Benchmarking universal single-copy orthologs (v3; https://busco.ezlab.org/).

| Assembly | Raw scaffolds (after chimera breaking) | In chromosome-scale pseudomolecules |  |
| --- | --- | --- | --- |
| Scaffolds | 109,776 | 476 |  |
| Total length (Mbp) | 6,670.03 | 6,206.74 |  |
| N50 length (Mbp) | 15.16 | 29.44 |  |
| % length with chromosome assignment | 95.3% | 100% |  |
| Optical genome map |  |  |  |
| Maps | 5,601 |  |  |
| Total length (Mbp) | 6,660.18 |  |  |
| N50 length (Mbp) | 1.671 |  |  |
| Total aligned length (Mbp) | 6,248.60 |  |  |
| Uniquely aligned length (Mbp) | 6,029.11 |  |  |
| Gene feature annotation | High confidence set | Low confidence set |  |
| Number of genes | 34,441 | 22,781 |  |
| Mean gene length | 2,892 | 946 |  |
| Mean exons per gene | 4.42 | 1.79 |  |
| Proportion of complete BUSCO set | 96.4% | 5.8% |  |
| LTR-RT annotation | Superfamily | Full-length copies | Mean age (Mya) |
| RLC_Angela | Copia | 11,128 | 0.53 |
| RLG_Cereba | Gypsy | 934 | 1.24 |
| RLG_Sabrina | Gypsy | 3,996 | 2.10 |
| RLG_WHAM | Gypsy | 1,457 | 2.06 |
| DTC_Clifford | CACTA | 1,480 | N.A. |
| DTC_Conan | CACTA | 516 | N.A. |

### Gene collinearity among the Triticeae

We used the assembly to closely assess gene-level collinearity between rye, barley and bread wheat (Methods; figs. M-TRACKSa, Note S-COLLIN)^9-11,15,22-24^. As previously reported, Triticeae chromosome groups 1–3 appear essentially collinear across all three species^9,10,15^. Rearrangements such as those between 4R and 7R are observable at high resolution, along with several inversions (e.g. on 1RL and 3RL; fig. M-TRACKSa). Rearrangements affecting subtelomeres were reflected in the absence of hybridisation signals from two subtelomere-specific FISH probes developed in this study (Note S-FISH; tbl. S-FISH). Regions of rye-barley collinearity contrast with distinct low-collinearity ‘modules’ (henceforth denoted LCMs) that surround the centromeres of chromosomes. Such regions, in which enough gene synteny is conserved to demonstrate identity by descent but the order of orthologs significantly differs among relatives, can now be observed in the sequenced genomes of many species^25,26^ (figs. M-TRACKSa; Note S-COLLIN). While centromeres can suffer from assembly difficulties, the LCM boundaries extend well into the pericentromeres, and on several chromosomes occur within large scaffolds validated by multiple sources of data including optical maps. The LCMs of rye, wheat, and barley differ in length, but curiously (i) the sets of genes that fall inside and outside the LCMs are almost the same in all three species, (ii) The LCMs distinctly correlate with regions of low gene density (fig. M-TRACKSb), and (iii) possess a distinct and characteristic repetitive element population (figs M-TRACKSd-g, Note S-REP). We explore these observations in more detail below.

**Figure M-FISH.**
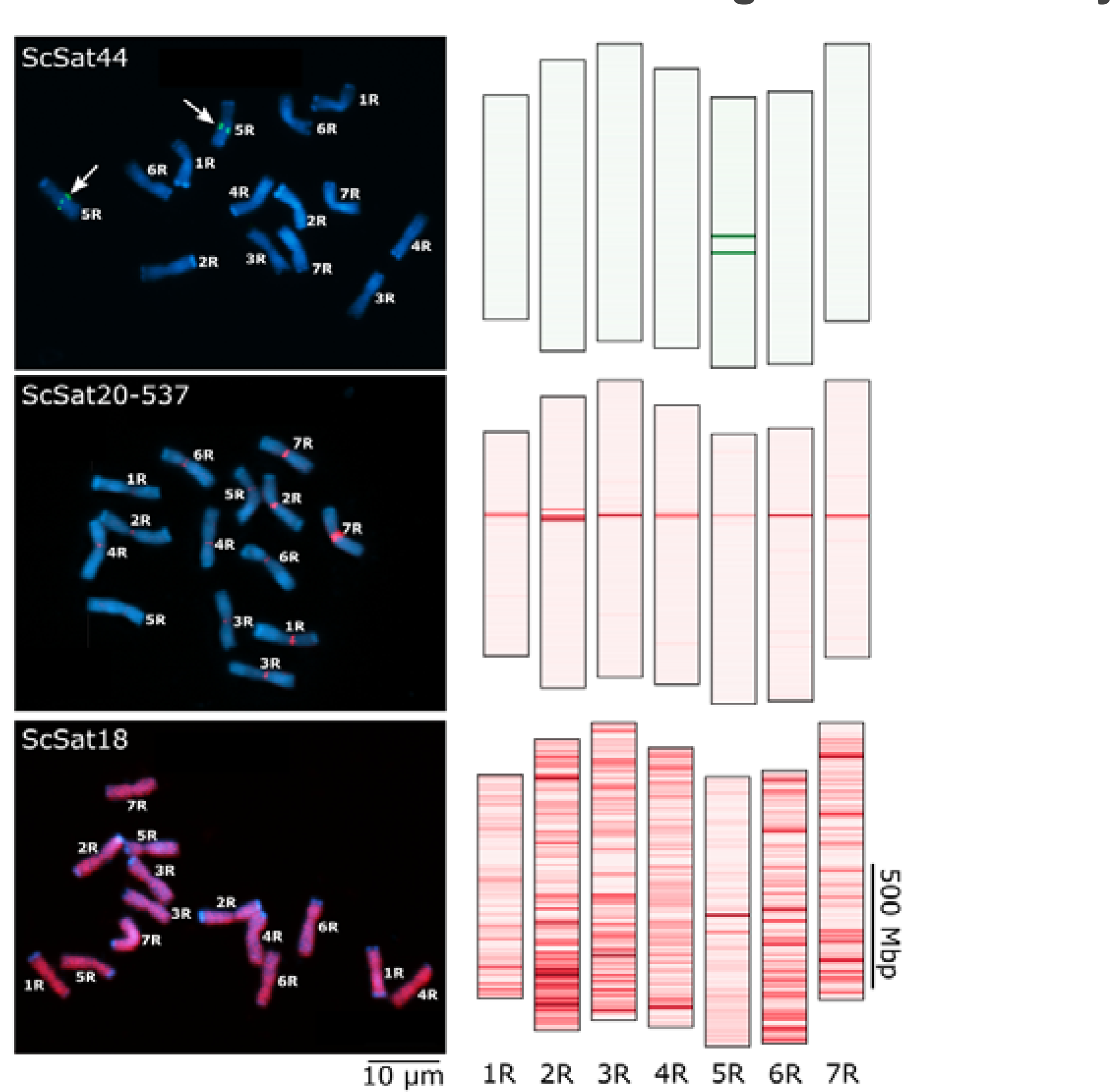
FISH of mitotic rye chromosomes with ScSat44, ScSat18 and ScSat20-537-specific probes (left) and *in silico* predicted repeat distribution (right), showing agreement between real and predicted hybridization sites. Chromosomes are counterstained with DAPI (blue), ScSat44 in green (chromosome 5R is arrowed), ScSat18 and ScSat20-537 in red. Arrows mark chromosome-specific binding of ScSat44 to chromosome 5R. Darkness is scaled evenly between the maximum and minimum densities of each repeat across all assembled chromosomes (Methods).

**Figure M-TRACKS.**
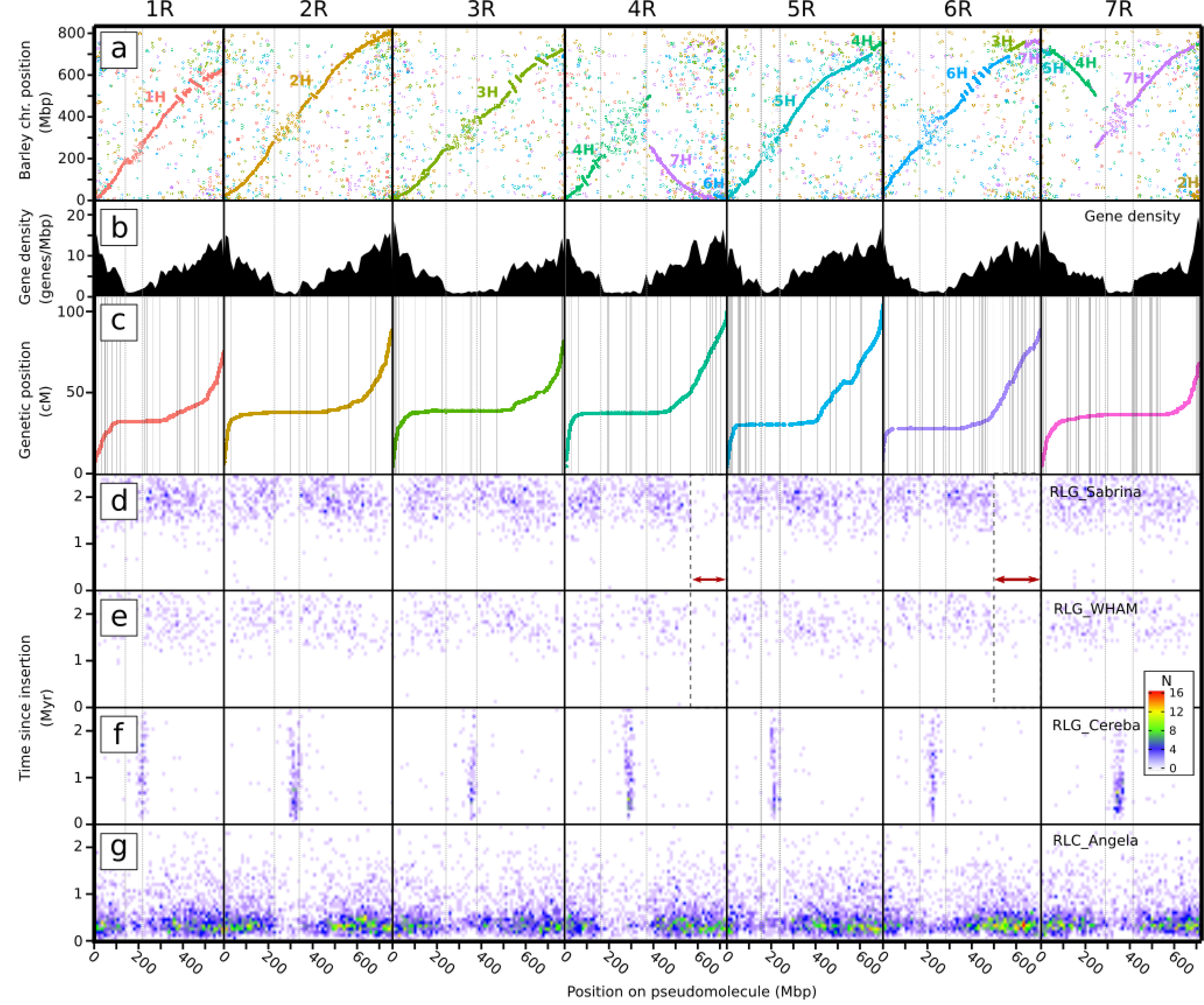
Selected information tracks for ‘Lo7’ chromosomes 1R to 7R (left-to-right). Twin vertical grey lines in each chromosome denote the boundaries of the LCMs for each chromosome. **A)** Gene collinearity with barley (cv. Morex), with the position on the Morex pseudomolecules on the vertical axis. Text and point colours represent barley chromosomes as labelled. **B)** Density of annotated gene models. **C)** Genetic map positions of markers used in assembly. Scaffold boundaries marked by grey vertical lines. **D-G)** Positions and ages of four LTR retrotransposon families in the genome, represented as a heatmap. Binned ages are on the vertical axis (from 0 Mya at the bottom), and bin positions are across the horizontal. Heat represents the number of TEs in each age/position bin (see legend inset). Red arrows mark notable changes in LTR-RT profiles.

### Evolutionary dynamics of the intergenic space

Transposable elements, especially long terminal repeat retrotransposons (LTR-RTs), exert a primary influence on Triticeae genome structure and composition^27-29^. Full-length LTR-RTs represent the same proportion of the total assembly size as exhibited by other major Triticeae reference assemblies (fig. S- RPT_ASSCMP, tbl. S-TE_ASSCMP_ANNOTSTATS), indicating similar assembly completeness^30^. Past LTR-RT activity can be inferred by estimating the insertion ages of individual LTR-RT elements, and the evolutionary relationships among LTR-RT families (Methods; Note S-REP).

As in barley and wheat, rye LTR-RT show clear niche specialisation across genomic compartments ^27,28^(fig. M-TRACKSc—f; Note S-REP): *RLC_Sabrina*, *RLG_WHAM*, and *RLC_Angela* are depleted in centromeres and pericentromeres, with the depleted region normally corresponding closely to the LCMs (fig. M-TRACKSb-f). *RLC_Cereba* strictly occupies centromeres^31^. The long arm termini of chromosomes 4RL and 6RL bear distinct tandem repeat (Note S-REP) and LTR-RT profiles (fig. M-TRACKSc,d; figs. S- TETERMPROF, S-KMERREP): *DTC_Clifford* elements are two to four times more abundant than on the long arm termini of the other chromosomes, while *RLG_Sabrina* and *RLG_WHAM* elements are almost absent. We suspect such changes are most likely the result of ancestral chromosome arm translocations from a close relative. In the case of 4RL the profile changes are particularly clear and we can ascertain that: (i) since the altered TE profile boundaries do not coincide with a collinearity break with wheat or barley (figs. M-TRACKSa, S-KMERREP; Note S-COLLIN), the donor is likely of rye lineage; (ii) since in the donated segments, *DTC_Clifford* is more abundant than *RLG_Sabrina* and *RLG_WHAM*, the donors must have diverged from the ‘Lo7’ ancestor prior to the expansion of the latter elements in earnest, around 3.5 Mya; and (iii) since the recent *RLC_Angela* expansion is recorded across 4R, the introgressions occurred before its beginning around 1.8 Mya.

The timing of expansions differs markedly between LTR-RT families of the rye genome, demonstrating that older families degrade as younger families expand. Repetitive insertion into the centromere suggests a centromere-outwards chromosome expansion mechanism, as is most apparent for chromosomes 2R, 4R, and 6R, by the distribution of older *Cereba* elements being more distant from the centromere than the younger. Comparing rye with wheat and barley, the variously curved and straight slopes of collinear runs of genes (Note S-COLLIN) suggest physical genome expansion acted quite uniformly across the rye genome since its split from wheat. Conversely, the size changes that separate rye from wheat and barley are pronounced near telomeres, indicating that genome expansion mechanisms alter over million year timescales and likely contribute to both speciation and ancient hybridisation events^32^. In rye, barley and in each individual wheat subgenome, the TE superfamilies *Gypsy* (RLG) and *Copia* (RLC) expanded in the same order^27,28^, but not at the same time: The *Gypsy-to-Copia* progression was probably set in motion by the LTR-RT composition of a shared ancestral genome, but the *rates* of expansion and suppression of each superfamily would have depended upon functional and selective peculiarities of each genome or sub-genome (arguments expanded in note S-TEEXP).

### Structural variation and *Secale* genome evolution

The many Triticeae gene-collinearity disruptions observable as inversions and pericentromeric LCMs suggest rapid accumulation of structural variations (SVs) that might segregate in rye populations causing undesired linkage in breeding and mapping efforts. To investigate further, we used Hi-C data from single individuals of four rye species to identify candidate SVs among *S. cereale* and three other *Secale species*. We included a second *S. cereale* genotype, ‘Lo225’, an inbred line from which the mapping population used for assembly was derived. To provide phylogenetic context, we extended the *Secale* phylogeny of Schreiber et al. (2019)^33^, adding 347 genotypes, and calling variants against the new genome assembly (Methods; fig. M-PHYLO). Many inversions (>10) were observed to segregate among non-‘Lo7’ *Secale* genotypes, making assembly artefacts a highly unlikely source of error (Note S-SV). One such ‘Lo7’— ‘Lo225’ inversion on 5RL corresponds to a distinct local plateau in the genetic map (fig. M-SV), representing complete linkage between the 382 annotated high confidence genes in this region. Rye pericentromeres are especially prone to large-scale SVs (p<0.001; Note S-SV), in agreement with previous findings^29,34^. This confirms SV as one possible mechanism for the formation of LCMs, and helps to explain the lack of genes in these regions, since recombination-suppressed genes are evolutionarily disadvantaged by Muller’s ratchet^35^. Such SVs likely contribute to phenotypic diversity (and potentially heterosis, as suspected for maize^36,37^), and influence *Secale* evolution by creating postpollination reproductive barriers that enable allopatric speciation^38^.

**Figure M-PHYLO.**
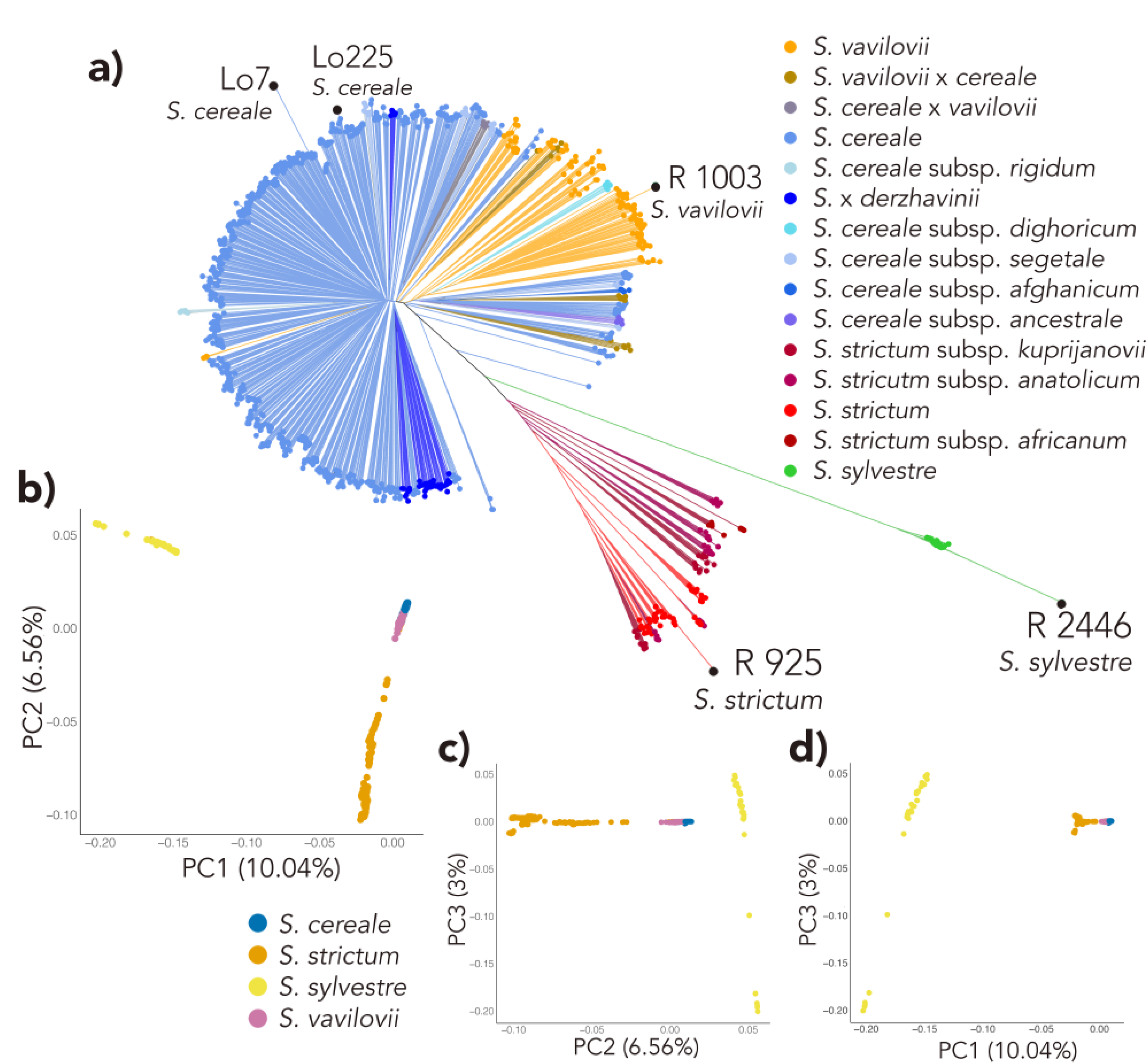
Diversity and relationships among *Secale* taxa. The population structure corresponds to the structure of three taxa as presented in Schreiber et al. 2019 but gives a clearer grouping due to the additional wild accessions, especially with regard to S. *vavilovii*, the wild progenitor, which was previously indistinguishable from domesticated rye but is now forming a subgroup within *S. cereale*. **a)** Neighbour- joining tree, with taxonomic assignments to subspecies level, according to genebank passport data. **b–d)** The first three prinicipal components of genetic variance within the dataset, with samples coloured according to species.

**Figure M-SV.**
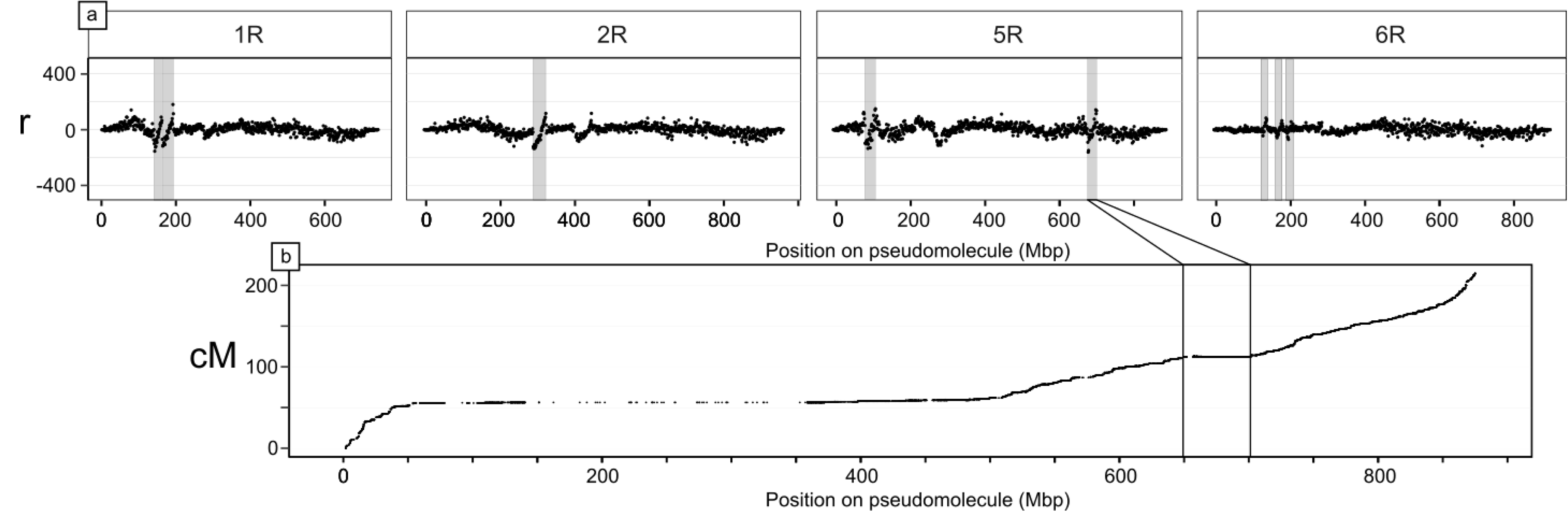
Hi-C asymmetry detects SVs between the reference genotype ‘Lo7’ and *S. cereale* ‘Lo225’ on four chromosomes. **a)** SVs result in discontinuities in *r*, the ratio of Hi-C links mapping left:right relative to ‘Lo7’. Large inversions (marked) typically produce clean, diagonal lines. Visually-identified candidate SVs are shaded, but shading is omitted from some *r* anomalies around centromeres where missing sequence causes artefacts. **b)** The rightmost inversion marked on 5R corresponds to a region of reduced recombination on chromosome 5R.

### Revised hypotheses on ancient translocations and the origin of the rye genome

It has been proposed that the cereal rye genome is a mosaic of Triticeae genomes resulting from reticulate evolution because variations in the degree of gene sequence divergence between various regions of the genome and their Triticeae orthologs indicate a number of distinct translocation donors^10^. We have presented evidence that the LTR-RT profile (figs. M-TRACKSd—e, Note S-REP) is a result of such reticulation within the rye lineage. It remains to be established whether significant chromatin introgressions occurred involving genera besides *Secale*. We exploited the new assembly to more closely investigate the cause of differential sequence divergence rates by estimating the divergence rate of synonymous coding sequence sites between rye and the wheat D genome (Methods, Note S-REP). The D genome was selected because it (i) contains no large chromosomal translocations relative to ancestral Triticeae karyotype (Note S-COLIN), and (ii) diverged from the ancestral rye genome only after the split from barley, meaning R-D divergence places a coarse lower bound on how much divergence it is possible to accumulate since the R-H split. The rates we recorded (∼0.06—0.14 subs/synonymous site/year) can account for the ∼5—15% identity spread of divergences that Martis et al. (2013) measured between rye and barley, without recourse to introgressions from beyond the R-D split. No cleanly-delimited divergence-level blocks are immediately evident to support extra-*Secale* introgressions. While some of the variation in divergence levels might yet be caused by such ancient translocations, inferring to what degree is confounded by other sources of random variation, probably including segregating recombination-suppressing SVs as observed in this study. We conclude that the mosaic hypothesis is indeed necessary to explain rye evolution, and currently most parsimonious when limited to introgressive hybridisations primarily between divergent *Secale* populations.

### Enhancing the benefits of rye germplasm in wheat breeding lines

The transfer of rye chromatin into bread wheat can provide substantive yield benefits and tolerance to biotic and abiotic stressors^39^, though at the expense of bread making quality^40^. These transfers are thought to have involved a single 1BL.1RS Robertsonian translocation originating from cv. Kavkaz and a single 1AL.1RS translocation from cv. Amigo (fig. M-INTROGa)^3,4^. Breeding efforts face a trade-off between yield and quality. Breeders must screen breeding panels for rye introgressions, an effort hitherto dependent upon arduous cytogenetics or marker genotyping, which has limited resolution and is sensitive to genetic variation among lines. With a full reference genome, inexpensive low-density high throughput sequencing (HTS) of a wheat panel proved sufficient to identify the positions of rye introgressions^41^. We implemented an HTS approach on four expansive wheat germplasm panels (KSU, USDA-RPN, CIMMYT, WHEALBI; Methods) segregating for both 1RS.1AL and 1RS.1BL. Translocations into wheat can be observed as obvious changes in normalised read depth across both the translocated and replaced chromosomal regions (fig. M-INTROGb; Note S-INTROG). A range of translocation junctions and karyotypes can be distinguished.

**Figure M-INTROG.**
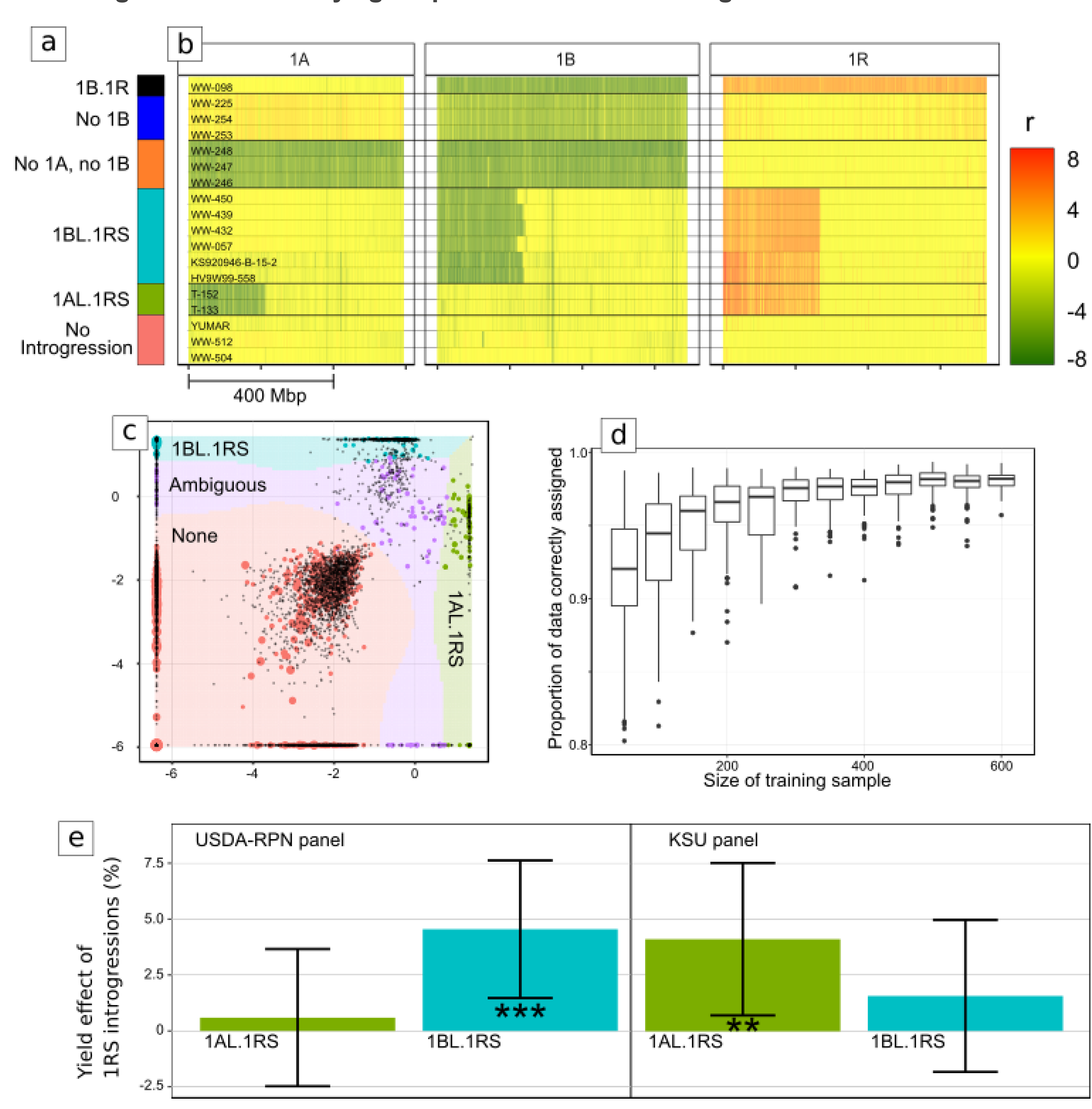
Combined reference mapping as a means to classify wheat and wheat-rye introgression karyotypes. **a)** Colour key for subfigures b, c, e. **b)** Normalised read mapping depths for 1 Mbp bins of chromosomes 1A, 1B, and 1R, for a selection of wheat lines (including also some *Aegillops tauschii* accessions which contain no A or B subgenome) with various chromosome complements and introgressions (rows). The value *r* denotes the difference between the log_2_ reads per million mapping to a bin, compared to *T. aestivum* cv. Chinese Spring. **c)** Visual representation of an SVM classifier, with the two selection features shown on the x and y axes. Points represent training samples, with colour corresponding to human-designated classification, and size proportional to the total number of mapped reads for the sample. Background colours represent the hypothetical classification that would be given to a sample at that position. **d)** Results of cross-validation testing the accuracy of the classifier and its relationship to the size of the training set. **e)** Comparison of yields between non-ambiguous predicted karyotypes, modelled using an MLM with testing year and location as random effects and rye introgressions as fixed effects. Results are shown for panels maintained by two institutions, USDA-RPN (left) and KSU (right). Bar height = Predicted yield effect of introgressions, +/- 1SD. Significant differences (Student’s t) given as: ‘**’ p<.01; ‘***’ p<.001.

The power of this sequence-based approach over previous markers was validated by confirming the karyotype of the novel 1AL.1RS—1BL.1RS recombination line KS090616K-1 (KSU panel; Note S-INTROG) that produces high yields, without sacrificing bread making quality. We confirmed that the KS090616K-1 breeding line carried a 1R translocation on group 1A, and after re-sequencing the wheat parents that carry donors of 1A.1R and 1B.1R, used high-density polymorphisms in the translocated 1R arm to precisely identify the recombination breakpoints, which fall at around 6 Mb from the tip of 1RS (Kavkaz- derived) onto the 1AL.1RS (Amigo-derived) line (Note S-INTROG). Moreover, this analysis conclusively confirmed the universal common origins of the Kavkaz- and Amigo-derived translocations respectively (Methods; Note S-INTROG).

Visual classification of a whole panel of karyotypes is still time-costly, so we developed an automated support vector machine classifier to alleviate this bottleneck (Methods; figs. M-INTROGc). Automatic classification consistently replicated human assignment with over 97% accuracy (fig. M-INTROGd). We then proved that the automated classifications predict yield. A mixed-effects linear model applied to yield data available for the USDA-RPN and KSU panels showed that 1R introgressions could produce ∼3— 5% better yields on average (Methods; fig. M-INTROGe; tbls. S-INTROGPHENO—S-GLMRES). The 1A.1R karyotype outyielded 1B.1R in the KSU panel, but the reverse was true of the USDA panel. This likely owes to the diversity of wheat genotypes and environmental conditions used in the trials; the effects of foreign chromatin are highly non-uniform and influenced by diverse factors, in particular the wheat genetic background^40,42^. Only one multi-site study has, to our knowledge, studied yield in 1RS- introgressed wheats on a large scale (Note S-1RS_PUBLIC), in which the best overall yield was achieved by a 1RS.1AL introgression line, both with and without the application of fungicidal treatments and during a drought year, while a 1RS.1BL line in the same panel performed less well, similarly suggesting significant variability in the pathogen resistance and root morphology traits that 1RS can confer to improve yield. Improved knowledge of the individual rye genes that confer these benefits is required to help untangle these factors.

### Rye genes for enhanced breeding and productivity

#### Enhanced fertilisation control: Rye as a model for hybrid breeding systems in Triticeae

Efficient hybrid plant breeding requires lines exhibiting either self-incompatibility (SI), switchable fertility control mechanisms, or gynoecy. Unlike wheat and barley, rye naturally enables both pollen guidance via SI, and switchable fertility via CMS and restorer-of-fertility (*Rf*) genes.

Rye’s SI is controlled by a two-locus system typical in Poaceae species. Pollen tube germination is suppressed when both stigma and pollen possess identical alleles at two SI loci, termed the *S*- and the *Z*- locus^6^, previously mapped to chromosomes 1R and 2R^43-45^ respectively. The breakdown of SI is poorly understood, yet essential for the development of inbred lines, which is in turn indispensable for producing heterotic seed and pollen parent lines in hybrid breeding. A DOMAIN OF UNKNOWN FUNCTION gene, designated *DUF247*, is a prime candidate for the *S*-locus in the related ryegrass (*Lolium perenne*, Poaceae, Tribe Poeae)^46^. We mapped the rye *S*-locus-controlled SI phenotype to an interval on 1R, which falls about 3 Mbp from the rye ortholog of *L. perenne’s DUF247 (SECCE1Rv1G0014240*; Methods; tbls. S-QTLS—S-1RSTS) Similarly, the *Z*-locus-linked marker *TC116908*^45^ mapped within about 0.2 Mbp of two other *DUF247* homologs (*SECCE2Rv1G0130770; SECCE2Rv1G0130780*) on 2R. This proximity suggests that *DUF247* might have been involved in SI since at least the time of the Triticeae— Poaceae split, making it a candidate for investigation relevant to barley and wheat^47,48^.

Turning to fertility control, mitochondrial genes that selfishly evolve to cause CMS prompt the evolution of nuclear *Rf* genes to suppress their expression or effects. Known *Rf* genes belong to a distinct clade within the family of pentatricopeptide repeat (PPR) RNA-binding factors, whose encoded proteins are referred to as *Rf*-like (RFL)^49,50^. Members of the mitochondrial transcription TERmination Factor (mTERF) family are likely also involved in fertility restoration in cereals^51^. The repertoire of restorer genes is predicted to expand in outcrossing species^35,52^. We investigated this hypothesis by comparing RFL and mTERF gene counts between rye and several closely and distantly allied species including barley and the subgenomes of various wheat species. The numbers of rye RFLs (n=82) and mTERFs (n=131) place it clearly within the range occupied almost exclusively by outcrossers (i.e. n_mTERF_ > 120 and n_RFL_ > 65; tbls. S- PPR_BREEDINGSYS; Note S-OUTIN), an indicator that rye’s younger RFL/mTERF genes evolved under selection to suppress CMS. The ‘Lo7’ sequence assembly reveals strong overlap in the distribution of PPR-RFLs and mTERF gene clusters, and strong correlation of these clusters with known *Rf* loci (Methods; fig. M-GENESa—f; tbls. S-QTL, S-PPR, S-MTERF). A PPR-RFL/mTERF hotspot on 4RL coincides with known *Rf* loci for two rye CMS systems known as CMS-P (the commercially predominant ‘Pampa’- type) and CMS-C^7,14,53,54^ (fig. M_GENESb,e,f; tbls. S-PPR, S-QTL). We determined, as previously hypothesised, that these two loci, *Rfp* and *Rfc*, are indeed closely linked but physically distinct^55^ (tbl. S- QTLS). Two members of the PPR-RFL clade reside within 0.186 Mbp of the *Rfc1* locus (tbls. S-PPR, S-QTL). The *Rfp* locus is, in contrast, neighboured by four mTERF genes (tbls. S-MTERF, S-QTL), in agreement with previous reports that an mTERF protein represents the *Rfp1* candidate gene in rye^56^.

**Figure M-GENES.**
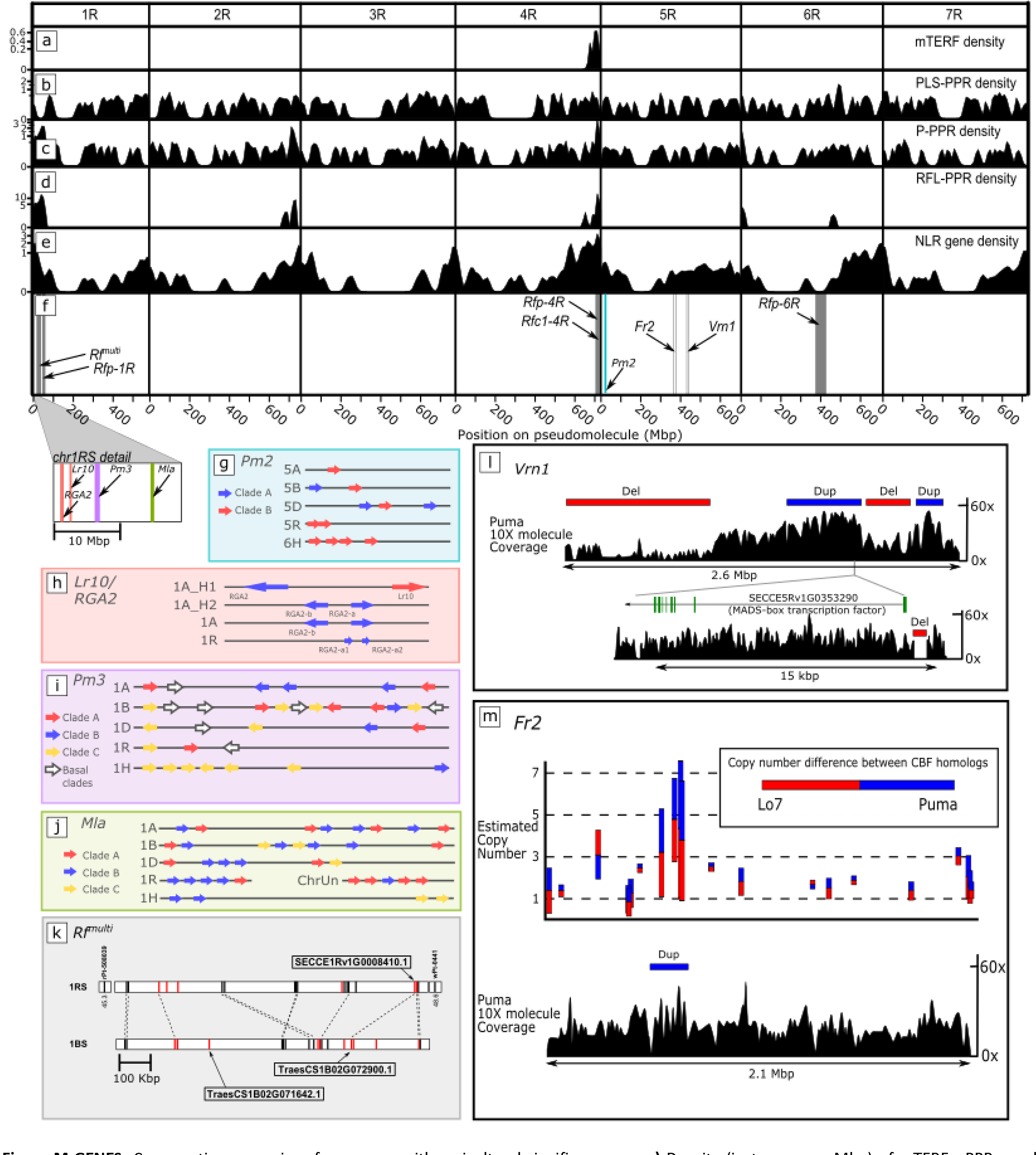
Comparative genomics of rye genes with agricultural significance. **a—e)** Density (instances per Mbp) of mTERFs, PPRs, and NLRs across the pseudomolecules (see also tables S-NLR to S-MTERF). For visualisation, the y-axis is transformed using x→ x⅓. **f)** Genes and loci discussed in the text (see also table S-QTLs). Colours correspond to box outlines in panels g—m; **g—j)** Physical organisation of selected NLR gene clusters compared across cultivated Triticeae genomes. **k)** Organisation of RFL genes at the ‘Lo7’ Rf^*multi*^ locus compared to its wheat (Chinese Spring) counterpart. Flanking markers are shown on either end of the rye sequence. Two full-length wheat RFLs and a putative rye ortholog are labelled. PPR genes are coloured red. **l—m)** CNV between ‘PUMA-SK’ and ‘Lo7’ revealed by 10X Genomics linked read sequencing. (Dup)lications and (Del)etions flagged by the Loupe analysis software are marked. The estimated copy number differences between ‘Lo7’ and ‘Puma’ are shown for *Cbf* genes within the *Fr2* interval.

While the most commonly used restorer cytoplasm in wheat hybrid breeding is derived from *Triticum timopheevii* Zhuk. (CMS-T)^57^, alternative sterility-conferring cytoplasms acquired from *Aegilops kotschyi* Bois., *Ae. uniaristata* Vis. and *Ae. mutica* Bois.^58^ can be efficiently restored by the wheat locus *Rf^multi^* (*Restoration-of-fertility* in multiple *CMS systems*) on chromosome 1BS. Replacement of the *Rf^multi^* locus by its rye ortholog produces the male-sterile phenotype^59,60^. Characterising this pair of sterility-switching genes could expedite flexible future solutions for the development of exchangeable wheat restorer lines. At the syntenic position of *Rf^multi^*, the wheat B subgenome and rye share a PPR-RFL gene cluster— with almost twice the number of genes in wheat^9^ (fig. M-GENESm; tbl. S-QTLs; Notes S-OUTIN—S- RFMULTI). Only two wheat RFL-PPR genes in the cluster, *TraesCS1B02G071642.1* and *TraesCS1B02G072900.1*, encode full length proteins with only the latter corresponding to a putative rye ortholog (*SECCE1Rv1G0008410.1*). Thus, the absence of a *TraesCS1B02G071642.1* ortholog in the non-restorer rye suggests it as an attractive *Rf^multi^* candidate. The only current implementations of a wheat- rye *Rf^multi^* CMS system involve 1RS.1BL translocations^5,58,61^, which are typically linked to reduced baking quality^40^. Breaking this linkage may now benefit from marker development and/or genome editing approaches targeting *TraesCS1B02G071642.1*.

#### New allelic variety in NLR genes and opportunities for pathogen resistance

Nucleotide-binding-site and leucine-rich repeat (NLR)-motif containing genes commonly associate with pest and pathogen resistance^62^. We annotated 792 full-length rye NLR genes (tbls. S-NLR—S-RLOC), finding them enriched in distal chromosomal regions, similar to what has been seen recently in the bread wheat genome^9,63^ (fig. M-GENESa; Note S-NLR). Distal parts of chromosomes 4RL and 6RL, which bear a distinct TE composition, are also particularly rich in NLR genes, further corroborating a unique, evolutionary-distinct origin for these segments.

We compared the genomic regions in rye that are orthologous to resistance gene loci *Pm2, Pm3, Mla, Lr10* from wheat and barley (tbl. S-RLOC; fig M_GENESg-j; Note S-NLR). Besides the *Lr10* locus, all loci contained complex gene families with several subfamilies that were present or absent in some genomes, indicating either functional redundancy, or the evolution of distinct resistance pathways or targets. For example, the wheat *Pm3* and rye *Pm8/Pm17* genes are orthologs and belong to a subfamily (clade A, fig. M_GENESi) which is absent in barley, whereas a different distinct subfamily (clade B, fig. M-GENESi) of the *Pm3* genes is present in wheat and barley but absent in rye (fig. M-GENESi, Note S-NLR). A similar case occurs in the *Mla* family: One of two main identified clades (clade B, fig. M_GENESj) contains known wheat resistance genes *TmMla1*^64^, *Sr33* and *Sr50*^65^ and yet is absent in barley, while a second *Mla* subfamily (clade C, fig. M_GENESj) contains all known barley *Mla* resistance alleles^66^, yet the clade is absent from rye (Note S-NLR). Rye inbred line ‘Lo7’, therefore appears to have lost whole subclades of pathogen resistance genes since its split from wheat.

#### The genetic basis of cold tolerance in rye, and its applications to wheat

As the most frost tolerant crop among the Triticeae^67^, rye is an ideal model to investigate the genetic architecture of low temperature tolerance (LTT) in cereals. Genetic mapping has revealed a locus *Fr2* on the group 5 chromosomes controlling LTT^68^ in rye^69^, *T. monococcum*^70^, bread wheat^71,72^, and barley^73^. In cold-tolerant varieties, the *Fr2* locus up-regulates LTT-implicated *Cbf* genes during seedling development under cold conditions^74^. *Cbf* genes are highly conserved in the Triticeae^75^. We identified the *Fr2* locus as a cluster of 21 *Cbf*-related genes at 614.3—616.5 Mbp on 5R (tbl. S-FR2). The region also contained 12 other genes that have been implicated in plant development, such as MYB transcription factors and a FAR1-related gene (tbl. S-FR2). A comparison of annotated Triticeae protein sequences within *Fr2* suggest the *Cbf* gene family expanded in rye, a mechanism for rye’s LTT, consistent with findings from other Triticeae^76^ (Note S-COLD).

To identify variation that may be important for cold acclimation we used recurrent selection to develop an *Fr2* homozygous line of the self-incompatible rye variety ‘Puma’, which exhibits exceptional LTT. We sequenced 10X Genomics Chromium libraries of this line (designated ‘Puma-SK’) and performed a comparison to the ‘Lo7’ reference sequence as a control since ‘Lo7’ has comparatively poor LTT. Mapping depth analysis detected copy number variation (CNV) patterns in four *Fr2 Cbf* genes (*SECCE5Rv1G030450, SECCE5Rv1G030460, SECCE5Rv1G030480*, and *SECCE5Rv1G030490*; fig. M-COLDm; tbl. S-CNV; Note S-COLD). Encouragingly, all four are members of the *Cbf* subfamily (‘group IV’, see fig. S- CBFPHYLO) for which CNV has been previously implicated in LTT in wheat^76^. Interestingly, we also detected a 597 bp deletion in the promoter of ‘Puma’’s *Vrn1* (SECCE5Rv1G0353290) allele. Although the effect of this deletion on LTT is not yet established, *Vrn1* is known to progressively down-regulate the expression of LTT genes during the vegetative/reproductive transition, impairing the plant’s ability to acclimate to cold stress^77,78^.

**Figure M-COLD.**
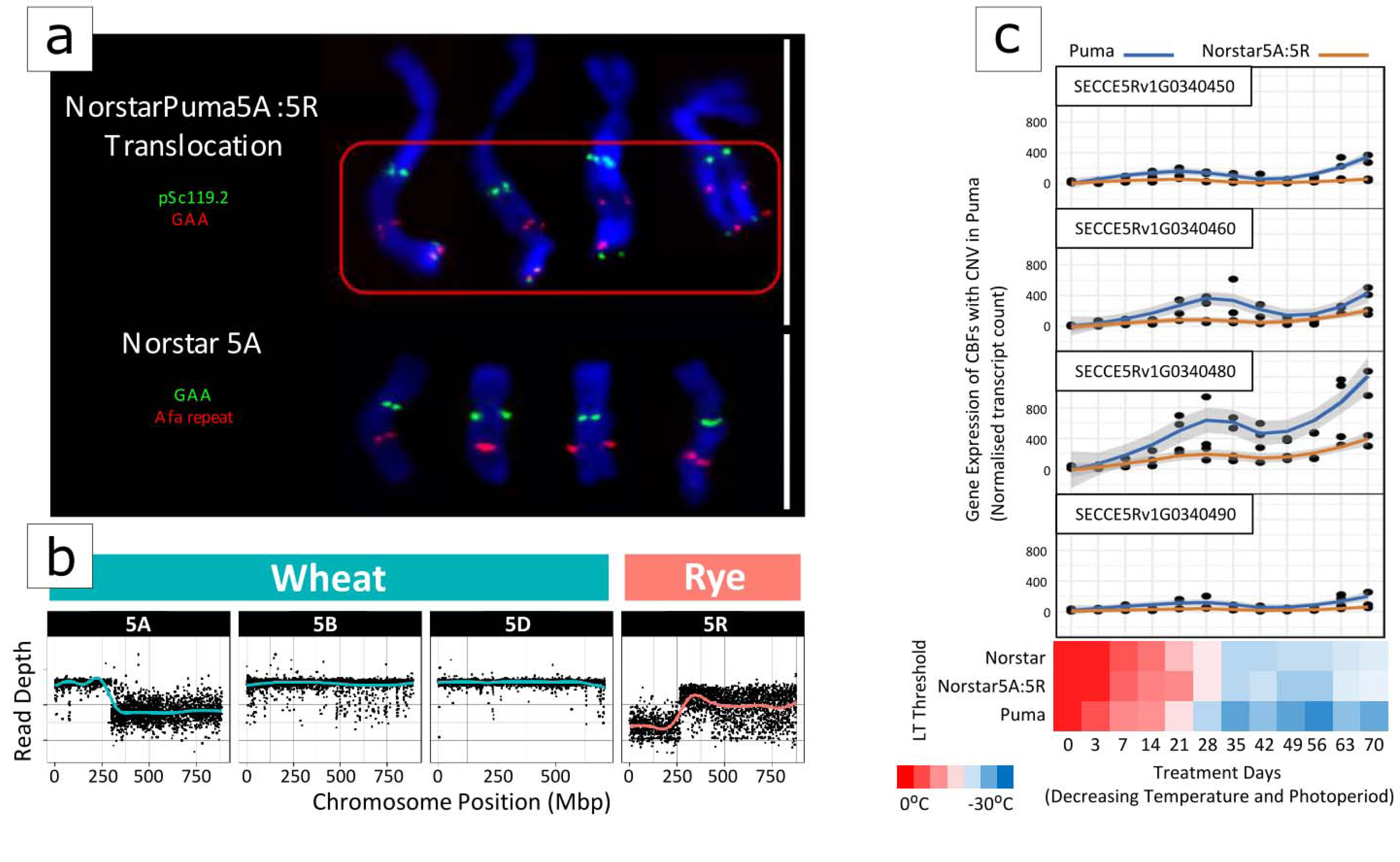
Cold tolerance region *Fr2* in ‘Puma’ and ‘NorstarPuma5A:5R’ translocation line. **a)** Chromosome labelling (top) using wheat and rye specific probes for chromosome 5A in ‘Norstar’ and 5A:5R in the ‘Puma’/‘Norstar’ translocation line confirms the presence of a rye translocation (red box). Read depth (bottom) of group 5 chromosomes confirms the balanced translocation event, gain of a large region of chromosome 5R from ‘Puma’ (rye - light read line) and loss of a large region on chromosome 5A of ‘Norstar’ (wheat - light blue line) in ‘NorstarPuma5A:5R’. White bars = 10 μm. **b)** Confirmation of the 5A.5R translocation into ‘Norstar’ using the combined reference mapping method. Read depth is given in log2 reads per million vs Chinese Spring. **c)** Gene expression analysis of rye *Cbf* genes with copy number variation in ‘Puma’ (blue line) and ‘NorstarPuma5A:5R’ (orange line). Plants were grown in a time series with decreasing day length and temperature over a 70 day period and the temperatures at which fifty percent lethality was observed (LT_50_) were recorded (heatmap).

We also assessed LTT-implicated genes’ potential for transfer to other members of the Triticeae, mainly wheat. ‘Norstar’ winter wheat is an important Canadian line with LTT sufficient to allow experiments in the Canadian winter—but weaker than ‘Puma’’s LTT, making it suitable for a comparison of LTT between wheat and rye^77^. A locus influencing ‘Norstar’’s superior LTT occurs on chromosome 5A^71^ and, like ‘Lo7’, contains tandemly repeated *Cbfs*^79^. We thus developed a 5A.5RL translocation line in the ‘Norstar’ winter wheat genetic background using ‘Puma’ as the 5R donor, which we confirmed using cytogenetics and combined reference mapping (Methods; fig. M-COLDa,b). As a result of the translocation, the wheat *Cbf* and *Vrn1* cluster is replaced completely by the orthologous rye locus (fig. M-COLDb; tbl. S-CNV). However, the LTT of ‘Norstar’ was not significantly altered by the translocation (fig. M-COLDc), suggesting that the rye *Cbf* gene cluster is activated in wheat, but it is differentially regulated in the wheat background, as previously suggested by Campoli et al. (2009)^74^. We used RNAseq to confirm that expression of ‘Puma’ *Vrn1* and those *Cbfs* with CNV were indeed attenuated during treatments of cold stress in ‘Norstar5A:5R’ (fig. M-COLDc; Note S-COLD). Characterisation of these important regulatory factors is an ongoing effort, necessary to facilitate improvement of wheat temperature tolerance using rye cytoplasm introgressions.

## Discussion

The high-quality chromosome-scale assembly of rye inbred line ‘Lo7’ constitutes an important step forward in genome analysis of the Triticeae crop species, and complements the resources recently made available for different wheat species^16,26,80-82^ and barley^15,83^. This resource will help reveal the genomic basis of differences in major life-history traits between the self-incompatible, cross-pollinating rye and its selfing and inbreeding relatives barley and wheat. Our comparative genomic exploration demonstrates how LTR-RT movement histories influence genome expansion and record ancient translocations. The precise nature and origin of the LCMs remains an opportunity for future research, requiring harmonisation of knowledge about the mechanics of pericentromeric structural variation, and the evolutionary effects of gene order disruption. The joint utilisation of the rye and wheat genomes to characterise the effects of rye chromatin introgressions may provide a short-term opportunity to breeders as they continue to better separate confounding variables from the genetic combinations that best improve yield in various environments; but these benefits will ultimately be limited by negative linkage so long as whole chromosome arm translocations are involved. Discoveries at the single-gene level—such as the contributions offered here to pathogen resistance, LTT, the root system (tbl. S-QTLs), SI, and male fertility restoration control—will be best tested and exploited by finer-scale manipulation in dedicated experiments^14^. This is an indispensable pre-requisite for the development of gene-based strategies that exploit untapped genetic diversity in breeding materials and *ex situ* gene banks to improve small grain cereals and meet the changing demands of global environments, farmers and society.

## Methods

### ‘Lo7’ genome assembly

Descriptions of the assembly methods are given in notes S-PSASS—S-ASSDATA, and figures S-ASSOVER— S-HICSV.

### Gene annotation

We performed de novo gene annotation of the rye genome relying on a previously established automated gene prediction pipeline^15,82^. The annotation pipeline involved merging three independent annotation approaches, the first based on expression data, the second an *ab initio* prediction for structural gene annotation in plants and the third on protein homology. To aid the structural annotation, RNAseq data was derived from five different tissues/developmental stages, and IsoSeq data from three (Supplementary Note 3).

IsoSeq nucleotide sequences were aligned to the rye pseudomolecules using GMAP^84^ (default parameters), whereas RNASeq datasets were first mapped using Hisat2^85^ (arguments --dta) and subsequently assembled into transcript sequences by Stringtie^86^ (arguments -m 150 -t -f 0.3). All transcripts from IsoSeq and RNASeq were combined using Cuffcompare^87^ and subsequently merged with Stringtie (arguments --merge -m 150) to remove fragments and redundant structures. Transdecoder github.com/TransDecoder) was then used to find potential open reading frames (ORFs) and to predict protein sequences. BLASTp^88^ (ncbi-blast-2.3.0+, arguments -max_target_seqs 1 -evalue 1e-05) was used to compare potential protein sequences with a trusted set of reference proteins (Uniprot Magnoliophyta, reviewed/Swiss-Prot) and hmmscan^89^ was employed to identify conserved protein family domains for all potential proteins. BLAST and hmmscan results were fed back into Transdecoder-predict to select the best translations per transcript sequence.

Homology-based annotation is based on available Triticeae protein sequences, obtained from UniProt (uniprot.org). Protein sequences were mapped to the nucleotide sequence of the pseudomolecules using the splice-aware alignment software GenomeThreader (http://genomethreader.org/; arguments - startcodon -finalstopcodon -species rice -gcmincoverage 70 -prseedlength 7 -prhdist 4). Evidence-based and protein homology based predictions were merged and collapsed into a non-redundant consensus gene set. *Ab initio* annotation using Augustus^90^ was carried out to further improve structural gene annotation. To minimise over-prediction, hint files using IsoSeq, RNASeq, protein evidence, and TE predictions were generated. The wheat model was used for prediction.

Additionally, an independent, homology-based gene annotation was performed using GeMoMa ^91^ using eleven plant species: *Arabidopsis thaliana* (n=167), *Brachypodium distachyon* (314), *Glycine max* (275), *Mimulus guttatus* (256_v2.0), *Oryza sativa* (323), *Prunus persica* (298), *Populus trichocarpa* (444), *Sorghum bicolor* (454), *Setaria italica* (312), *Solanum lycopersicum* (390), and *Theobroma cacao* (233). All versions were downloaded from Phytozome (phytozome.jgi.doe.gov/pz). Initial homology search for coding exons was done with mmseqs2^92^. These results were then combined into gene models with GeMoMa using mapped RNAseq data for splice site identification. The resulting eleven gene annotation sets were further combined and filtered using the GeMoMa module GAF. The following filters were applied: a) complete predictions (i.e. predictions starting with Methionine and ending with a stop codon); b) relative GeMoMa score >=0.75; c) evidence>1, (i.e. predictions were perfectly supported by at least two reference organisms), or tpc=1 (i.e., predictions were completely covered by RNA-seq reads), or pAA>=0.7 (i.e., predictions with at least 70% positive scoring amino acid in the alignment with the reference protein).

All structural gene annotations were joined with EvidenceModeller^93^, and weights were assigned as follows: Expression-based Consensus gene set (RNAseq, and IsoSeq and protein homology-based): 5; homology-based (GeMoMa), 5; ab initio (augustus), 2.

In order to differentiate candidates into complete and valid genes, non-coding transcripts, pseudogenes and transposable elements, we applied a confidence classification protocol. Candidate protein sequences were compared against the following three manually curated databases using BLAST: firstly PTREP (botserv2.uzh.ch/kelldata/trep-db), a database of hypothetical proteins that contains deduced amino acid sequences in which, in many cases, frameshifts have been removed, which is useful for the identification of divergent TEs having no significant similarity at the DNA level; secondly UniPoa, a database comprised of annotated Poaceae proteins; thirdly UniMag, a database of validated magnoliophyta proteins. UniPoa and UniMag protein sequences were downloaded from Uniprot (www.uniprot.org/) and further filtered for complete sequences with start and stop codons. Best hits were selected for each predicted protein to each of the three databases. Only hits with an E-value below 10e-10 were considered.

Furthermore, only hits with subject coverage (for protein references) or query coverage (transposon database) above 75% were considered significant and protein sequences were further classified using the following confidence: a high confidence (HC) protein sequence is has at least one full open reading frame and has a subject and query coverage above the threshold in the UniMag database (HC1) or no BLAST hit in UniMag but in UniPoa and not TREP (HC2); a low confidence (LC) protein sequence is not complete and has a hit in the UniMag or UniPoa database but not in TREP (LC1), or no hit in UniMag and UniPoa and TREP but the protein sequence is complete.

The tag REP was assigned for protein sequences not in UniMag and complete but with hits in TREP. Functional annotation of predicted protein sequences was done using the AHRD pipeline (github.com/groupschoof/AHRD). Completeness of the predicted gene space was measured with BUSCO (v3; https://busco.ezlab.org/).

### RNA isolation and sequencing

#### RNA-seq for annotation

Seeds of ‘Lo7’ were sown in a Petri dish on moistened filter paper and treated with cold stratification (4 °C) for two days during imbibition. After an additional day at room temperature (∼20 °C) seedlings were transferred to a 40-well tray containing a peat and sand compost and propagated in a Conviron BDW80 cold environment room (CER; Conviron) with set points of 16 h day/8 h night and temperatures of 20/16 °C for a further three days. Tissues were sampled at six stages, described in table S-RNAGROWTH. Plants for sampling timepoints 1—3 were transferred to a CER set at 16-hour photoperiod (300 μmol m−2 s−1), temperatures of 20 and 16 °C, respectively, and 60% relative humidity. Plants for sampling timepoints 4—6 were transferred to a vernalisation CER running at 6 °C with 8 hours photoperiod for 61 days. After this period the plants were transferred to 1 L pots containing Petersfield Cereal Mix (Petersfield, Leicester, UK) and moved to the CER with settings as described above. Total RNA was extracted from each of the six organ/stages using RNeasy plant mini-kits (Qiagen). For the RNAseq data sets used for the annotation. RNA from 3 biological replicates for each organ/stage was pooled and for the 6 pooled samples, library construction and sequencing on the Illumina NovaSeq platform was performed by Novogene using a standard strand specific protocol (en.novogene.com/next-generation-sequencing- services/gene-regulation/mrna-sequencing-service) and generating >60 M 150 PE reads per sample.

For the IsoSeq data used in the annotation RNA from root and shoot samples were used (timepoints 1 and 2 in table S-RNAGROWTH). The IsoSeq libraries were created starting from 1µg of total RNA per sample and full-length cDNA was then generated using the SMARTer PCR cDNA synthesis kit (Clontech) following PacBio recommendations set out in the IsoSeq method (pacb.com/wp-content/uploads/Procedure-Checklist-Iso-Seq-Template-Preparation-for-Sequel-Systems.pdf). PCR optimisation was carried out on the full-length cDNA using the KAPA HiFi PCR kit (Kapa Biosystems) and 10—12 cycles was sufficient to generate the material required for SMRTbell library preparation. The libraries were then completed following PacBio recommendations, without gel-based size-selection (pacb.com/wp-content/uploads/Procedure-Checklist-Iso-Seq-Template-Preparation-for-Sequel-Systems.pdf).

The library was quality checked using a Qubit Fluorometer 3.0 (Invitrogen) and sized using the Bioanalyzer HS DNA chip (Agilent Technologies). The loading calculations for sequencing were completed using the PacBio SMRTlink Binding Calculator v5.1.0.26367. The sequencing primer from the SMRTbell Template Prep Kit 1.0-SPv3 was annealed to the adapter sequence of the libraries. Each library was bound to the sequencing polymerase with the Sequel Binding Kit v2.0. Calculations for primer and polymerase binding ratios were kept at default values. Sequencing Control v2.0 was spiked into each library at ∼1% prior to sequencing. The libraries were prepared for sequencing using Magbead loading onto the Sequel Sequencing Plate v2.1. The libraries were sequenced on the PacBio Sequel Instrument v1, using 1 SMRTcell v2 per library. All libraries had 600-minute movies, 120 minutes of immobilisation time, and 120 minutes pre-extension time (tbl. S-DATACCESS).

#### RNA-seq for expression profiling of ‘NorstarPuma5A:5R’ and ‘Puma’

Total RNA was extracted from 48 samples, representing both ‘NorstarPuma5A:5R’ and ‘Puma’ lines at each sampling date of the 12 time points during cold acclimation (Note S-COLD), using the Plant RNA Isolation Mini Kit (Agilent Technologies). The yield and RNA purity were determined spectrophotometrically with Nanodrop 1100 (Thermfisher), and the quality of the RNA was verified by Agilent 2100 Bioanalyzer (Agilent Technologies). Purified total RNA was precipitated and re-suspended in RNase-free water to a final concentration of 100 ng/µl. Libraries were constructed using the TruSeq RNA Sample Preparation Kit v2 (Illumina) with two replicates at each time point. Paired-end sequencing was conducted on the Illumina HiSeq2500, generating 101 bp reads (tbl. S-DATACCESS).

### Annotation of repetitive elements

For use in the evolutionary analyses presented in the main text (e.g. fig. 4d—g) annotated a high-stringency set of full-length transposon copies belonging to single TE families (tbl. S-TEANNOT) using BLASTn^88^ searches (using default parameters) against the ‘Lo7’ pseudomolecules for long terminal repeats (LTRs) documented in the TREP database (botinst.uzh.ch/en/research/genetics/thomasWicker/trep-db.html) that occur at a user-defined distance range in the same orientation: For RLC_Angela elements, the two LTRs had to be found within a range of 7,800—9,300 bp (a consensus RLC_Angela sequence has a length of approximately 8,700 bp), while a range from 6,000—12,000 bp was allowed for RLG_Sabrina and RLG_WHAM elements. For the centromere-specific RLG_Cereba elements, a narrower range of 7,600-7,900 bp was used. Multiple different LTR consensus sequences were used for the searches in order to cover the intra-family diversity. A total of 18 LTR consensus sequences each were used for RLC_Angela, seven for RLG_Sabrina elements, 6 were used for RLG_WHAM elements, and 5 for RLC_Cereba elements.

To validate the extracted TE populations, the size range of all isolated copies and the number of copies that flanked by target site duplications (TSDs) were determined. A TSD was accepted if it contained at least 3 matches between 5’ and 3’ TSD (e.g. ATGCG and ACGAG). This low stringency was applied because TSD generation is error-prone^94^, and thus multiple mismatches can be expected. Across all surveys, 80-90% of all isolated full-length elements were flanked by a TSD.

The pipeline also extracts so-called “solo-LTRs”—products of intra-element recombination that results in loss of the internal domain and generation of a chimeric solo-LTR sequence—as a metric of how short repetitive sequences are assembled.

The two LTRs of each TE copies were aligned with the program Water from the EMBOSS package^95^ and nucleotide differences between LTRs were used to estimate the insertion age of each copy based on the estimated intergenic mutation rate of 1.3E-8 substitutions per site per million years^96^.

Full-length DNA transposons were identified by BLASTn searches of consensus sequences of the terminal inverted repeats (TIRs) of a given family. TIRs were required to be found in opposite orientation in a user-defined distance interval of 7,000—15,000 bp.

To produce a library of full length LTR-retrotransposons suitable for quantitative assembly completeness comparison (fig. S-RPT_ASSCMP), we required an annotation performed identically to those carried out on other assemblies (tbls. S-TE_ASSCMP_ANNOTSTATS, S-DATAACCESS). We therefore implemented the methods described in Monat et al. (2019)^83^ on a selection of genome assemblies given in note S-REP.

Tandem repeats where annotated with TandemRepeatsFinder^97^ under default parameters (tbls. S- SAT_ANNOT, S-TANDREPCOMPN). Overlapping annotations where removed with a priority-based approach assigning higher scoring and longer elements first. Elements which overlapped already assigned elements were either discarded (>90% overlap) or shortened (<=90% overlap) if their remaining length exceeded 49 bp.

To obtain a collection of nonredundant tandem repeat units suited for FISH probe development, the consensus sequences of the tandem repeat units (output of TandemRepeatsFinder) where clustered with vmatch dbcluster (vmatch.de) at high stringency with >=98% identity and a mutual overlap >=98% (98 98 -v -identity 98 -exdrop 3 -seedlength 20 -d -p). The 300 largest clusters with member sizes from 199 to 343 where each subjected to a multiple sequence alignment with MUSCLE^98^ under default parameters. A consensus sequence (>=70% majority) derived per cluster from the MUSCLE score file served as template sequence for the FISH probes (tbl. S-FISH).

The distribution of TRs across the genome (main fig. M-FISH) was visualised using R base plotting functions. Colours were selected from the package colourspace palettes ‘Reds3’ and ‘Greens3’, e.g. using the command sequential_hcl(‘Reds3’,105)[105:5] to achieve 100 grades of a palette, and then selected to represent relative TR densities by scaling the output of the ‘density’ function run over the tandem repeats (with automatic bandwidth selection) on each chromosome to between 1 and 100 (for each TR family).

### Annotation of miRNAs

MicroRNA identification was performed by following a two-step homology-based pipeline. The ‘Lo7’ pseudomolecules were compared with all known mature plant miRNA sequences retrieved from miRBase^99^ (v21; www.mirbase.org). This step was performed using SUmirFind (https://github.com/ hikmetbudak/miRNA-annotation/blob/master/ SUmirFind.pl), an in-house script, and the matches with no mismatch or only one base mismatch between a mature miRNA sequence and the pseudomolecule sequence were accepted. A second in-house script, SUmirFold (https://github.com/hikmetbudak/miRNA-annotation/blob/master/ SUmirFold.pl), was used to obtain precursor sequences of the candidate mature miRNAs from the pseudomolecules and assess their secondary structure-forming abilities with UNAFold^100^ (tbls. S-miRNA1—S-miRNAX) together with the following criteria: 1) No mismatches are allowed at Dicer cut sites; 2) No multi-branched loops are allowed in the hairpin containing the mature miRNA sequence; 3) Mature miRNA sequence cannot be located at the head portion of the hairpin; 4) No more than 4 and 6 mismatches are allowed in the miRNA and its hairpin complement (miRNA*), respectively^101,102^. The final set of identified miRNAs from the pseudomolecules was obtained by SUmirScreen script (https://github.com/hikmetbudak/miRNA-annotation/blob/master/ SUmirScreen.py). The resulting miRNAs were mapped back to the pseudomolecules and the genomic distribution statistics were recorded with SUmirLocate script (https://github.com/hikmetbudak/miRNA-annotation/blob/master/ SUmirLocate.py).

Coding targets of the identified miRNAs were predicted by the web-tool psRNAtarget, using S. cereale coding sequences retrieved from NCBI^103,104^. Potential target sequences were compared with the viridiplantae proteins by using BLASTx^88^ (arguments -evalue 1E-6 –outfmt 5). Functional annotations of the potential targets were performed using Blast2GO software^105^. Finally, repeat contents of the pre-miRNAs were assessed with RepeatMasker (http://www.repeatmasker.org/).

### Fluorescence *in situ* hybridisation (FISH)

Three days old roots of the rye accession WR ‘Lo7’ were pre-treated in 0.002 M 8-hydroxyquinoline at 7°C for 24 h and fixed in ethanol:acetic acid (3:1 v/v). Chromosome preparation and FISH were performed according to the methods described by Aliyeva-Schnorr et al. (2015)^106^. The hybridization mixture contained 50% deionized formamide, 2× SSC, 20% dextran sulfate, and 5 ng/µl of each probe. Slides were denatured at 75°C for 3 min, and the final stringency of hybridization was 76%. Thirty-four to forty-five nt long 5’-labelled oligo probes designed for the *in silico* identified repeats and the published probes sequence pSc119.2.1^107^ were used as probes (tbl. S-FISH). Images were captured using an epifluorescence microscope BX61 (Olympus) equipped with a cooled CCD camera (Orca ER, Hamamatsu). Chromosomes were identified visually based primarily on morphology, heterochromatic DAPI+ bands, and the localisation of pSc119.2.1^107^.

### Rye Gene level synteny with other Triticeae species

High confidence gene sequences from the ‘Lo7’ gene annotation were aligned to the annotated transcriptomes of bread wheat^9^ (*Triticum aestivum* cv. Chinese Spring) and barley^15^ (*Hordeum vulgare* cv. Morex) using BLASTn^88^ with default parameters. The lowest E-value alignment for each gene against the transcriptome associated with each subject genome (or subgenome) was selected, with the longest alignment chosen in the case of a tie. Only reciprocal best matches per (sub/)genome were accepted. BLAST hit filtering and subsequent visualisation were performed in the R statistical environment exploiting the packages ‘data.table’ and ‘ggplot’.

### Wheat (D subgenome)—rye substitution rate variation across the genome

Probable orthologs shared by the wheat D subgenome^9^ and rye line ‘Lo7’ were identified by aligning BLASTp^88^ (default parameters) the predicted proteins of either each genome against the other and applying the reciprocal best match criterion. The identified homologs were first aligned at the protein level and, based on the protein alignment, a codon-by-codon DNA alignment was generated. For comparison of substitution rates, only fourfold degenerate third codon positions were used, namely those of the codons for Ala, Gly, Leu, Pro, Arg, Ser, Thr and Val. From the alignments of fourfold degenerate sites, the ratio of synonymous substitutions per synonymous site was calculated for each gene pair, if at least 100 fourfold degenerate sites could be aligned. Substitution rates along chromosomes were calculated as a 100 genes running average. Because even bi-directional closest homologs may still include “deep paralogs” (i.e. genes that were duplicated in the ancestor of which one copy was deleted in one species while the other copy is deleted in the other species), we performed the same analysis using exclusively single-copy genes. Single-copy genes were identified as follows: all individual rye coding DNA sequences (CDSs) were used in BLASTn searches against all other predicted rye CDSs. A gene was considered single-copy if it had no homologs with E-values below 10e-20. Substitution rates were then calculated as the rates of synonymous substitutions per synonymous site in fourfold degenerate codon sites in coding regions of genes.

### Phylogenetic analysis

The genotyping-by-sequencing (GBS) data set of 603 samples from Schreiber et al. (2019)^33^ was extended by a 347 further GBS samples from the IPK gene bank (mainly wild *Secale taxa*), and the five samples used in the Hi-C SV-detection study (‘Lo7’, ‘Lo225’, ‘R1003’, ‘R925’, ‘R2446’). The resulting sample set (n=955) and passport data are listed in table S-DIVERSITYPSPT. DNA isolated from the five Hi-C samples was sent to Novogene (en.novogene.com/) for Illumina library construction and sequencing in multiplex on the NovaSeq platform (paired end 150 bp reads, approximately 140 Gbp per sample, S2 flow cell). Demultiplexing, adapter trimming, read mapping and variant calling correspond to the approach described in Schreiber et al. (2019)^33^, using the new reference for read mapping. The data set was filtered for a maximum of 30% missing data and a minor allele frequency of 1% resulting in 72,465 SNPs used for the phylogenic analyses. A neighbor joining tree was constructed with the R package ‘ape’ version 5.3^108^, based on genetic distances computed with the R package SNPRelate^109^. PCA was performed with smartPCA from the EIGENSOFT package (github.com/DReichLab/EIG) using least square projection without outlier removal.

### Wheat-rye introgression haplotype identification and classification

We assayed for the presence of 1R germplasm in wheat genotypes *in silico* by mapping various wheat sequence data to a combined reference genome made up of the pseudomolecules of rye line ‘Lo7’ (this study) and wheat cv. Chinese Spring^9^. Publicly available data was obtained from the Wheat and barley Legacy for Breeding Improvement (WHEALBI) project resources^110^ (n=506), the International Maize and Wheat Improvement Centre (CIMMYT; n=903), and Kansas State University (KSU; n=4277). GBS libraries were constructed and sequenced for samples from the United States Department of Agriculture Regional Performance Nursery (USDA-RPN; n=875; tbl. S-DATAACCESS) as described in Rife et al. (2018)^111^. Based upon the approach described by Keilwagen et al. (2019)^91^, reads were demultiplexed with a custom C script (github.com/umngao/splitgbs) and aligned to the combined reference using bwa^112^ mem (arguments -M) after trimming adapters with cutadapt^113^. The aligned reads from all panels were filtered for quality using samtools^114^ (arguments flags -F3332 -q20). The numbers of reads aligned to 1 Mbp non-overlapping bins on each pseudomolecule were tabulated. The counts were expressed as *rpmm log_2_(reads mapped to bin per million reads mapped)*. To control for mappability biases over the genome, the *rpmm* for each bin was normalised by subtracting the *rpmm* attained by the Chinese Spring sample for the same bin to give the normalised *rpmm, r*.

To investigate the possibility of classifying the samples automatically, visual representations of *r* across the combined reference genome were inspected, and obvious cases of 1R.1A and 1R.1B introgression were distinguished from several other karyotypes including non-introgressed samples, and ambiguous samples showing a slight overabundance of 1RS reads, but less discernible signals of depletion in 1A or 1B (see Note S-INTROG). We defined the following feature vectors: *featureA = -log[ ( mean(r1A_I_) - mean(r1A_N_) ) x ( mean(r1R_I_) - mean(r1R_N_) ) ] and featureB = -log[ ( mean(r1B_I_) - mean(r1B_N_) ) x ( mean(r1R_I_) - mean(r1R_N_) ) ]*. Whenever the term inside the log was negative (and would thus give an undefined result), the value of the feature was set to the minimum of the defined values for that feature. The quantity *mean(r1R_I_)* refers to the average value of r for all bins within the terminal 200 Mbp of the normally (*I*)ntrogressed end of 1R (an _*N*_ in the subscript denotes the terminal 300 Mbp of the normally (*N*)on-introgressed arm), and so forth for other chromosomes. This choice of feature definition meant that, wherever little difference in r occurred between 1RS and 1RL, suggesting no presence of rye, the factor *mean(r1R_I_) - mean(r1R_N_)* would pull the feature values close to the origin, and differences between *r* on the long and short arms of 1A or 1B would pull the values of A or B respectively away from the origin, depending upon which introgressions are present. A classifier was developed by training a support vector machine to distinguish non-introgressed, 1A.1R-introgressed, 1B.1R-introgressed, and ambiguously-introgressed samples, using the function ksvm (arguments type=“C-svc”, kernel=’rbfdot’, C=1) from the R package kernlab. Classification results are given in table S-INTROG_PREDICTED. Testing was performed by generating sets of between 50 and 600 random samples from the dataset and using these to train a model, then using the kernlab::predict to test the model’s accuracy of prediction on the remaining data not used in training. This was repeated 100 times for each training data set size.

To investigate the 1R-recombinant genotype KS090616K, raw reads of genotypes Larry, TAM112 and KS090616K (NCBI SRA project id: PRJNA566411) were mapped to the combined wheat/rye reference, and mapping results processed with samtools^114^. The bcftools^114^ mpileup and call functions were used to detect and genotype single-nucleotide polymorphisms (SNPs) between the two samples. SNP positions at which Larry and TAM112 carried different alleles were used to partition chromosome 1RS in KS090616K into parental haplotypes (Note S-INTROG).

To confirm the common origin of the 1AL.RS and 1BL.1RS introgressions, predicted 1RS carriers were selected to form a combined 1RS panel (over twelve hundred lines) to call SNPs. A total of over 3 million SNPs were called with samtools/bcftools (mpileup -q 20, -r chr1R:1-300000000; call -mv). SNPs were filtered based on combined minimum read depth of 25, minor allele frequency of 0.01. A total of over 900 thousand SNPs were obtained. All pair-wise identity by state (IBS) percentages were calculated and the square root values of percent different calls were used to derive a heatmap for all pair-wise comparisons.

### Identification and analysis of gene families

#### Resistance gene homologs

To investigate rye homologs of the wheat and barley genes *Pm2, Pm3, Mla, Lr10* and *RGA2* (GeneBank IDs in tbl. S-NLRSEARCH), homology searches were performed against the rye ‘Lo7’, bread wheat^9^ (cv. Chinese Spring), and barley^15^ (cv. Morex) genome sequences, using BLASTn^88^ (default parameters). Hits with at least 80% sequence identity were visualised using dotter^115^ for manual assessment and annotation. The obtained coding sequences were converted to protein sequences, allowing comparison with the EMBOSS program WATER (emboss.sourceforge.net), ClustalW^116^, or MUSCLE^98^, with reference sequences and other obtained sequences to aid distinction between potentially functional full-length genes, and pseudogenes with truncations or premature stop codons.

Annotated genes were aligned using MUSCLE (default parameters), and the phylogenetic relationships among them were inferred using MisterBayes^117^ (GTR substitution model with gamma distributed rate, variation across sites, and a proportion of invariable sites).

Manually-annotated positions of the genes *Pm2, Pm3, Mla, Lr10* and *RGA2* on the ‘Lo7’ pseudomolecules were compared with the annotated NLR genes identified by the gene feature annotation pipeline (described above) in order to link the genome-wide NLR analysis with the detailed analysis of the four R loci. Pairwise distances between NLRs were calculated based on the resultant tree using the cophenetic.phylo function in the R package ‘ape’^108^, and multidimensional scaling on the pairwise distances was conducted with the core R function ‘cmdscale’.

#### PPR and mTERF genes

The ‘Lo7’ pseudomolecules were scanned for ORFs with the getorf program of the EMBOSS package^95^. ORFs longer than 89 codons were searched for the presence of PPR motifs using hmmsearch from the HMMER^118^ package (http://hmmer.org) and the profile hidden Markov models (HMMs) as defined in Cheng et al. (2016)^119^ for the PPR family PF02536 from the Pfam 32.0 database (http://pfam.xfam.org) and for the mTERF motif^120^. Downstream processing of the hmmsearch results for the PPR proteins followed the pipeline described in Cheng et al. (2016)^119^. A score was attributed to each PPR sequence (the sum of hmmsearch scores for all PPR motifs in the protein). In parallel, the HC and LC protein models from the gene feature annotation (described above) were screened to identify the annotated proteins containing PPR motifs. Five-hundred and twenty-six PPR models were identified in the HC and seventy-six in the LC protein datasets respectively, and scored using the same approach as with the hmmsearch results. Where putative exons identified from the six-frame translations of the genome sequence overlapped with gene models in the ‘Lo7’ annotation, only the highest scoring of the overlapping models were retained. P- and PLS-class genes with scores below 100 and 240, respectively, were removed from the annotation, as they are unlikely to represent functional PPR genes. Only genes encoding mTERF proteins longer than 100 amino acids were included in the final annotation.

#### Mapping genes governing the reproduction biology in rye

Molecular markers previously mapped in relation to Rf and SI genes were integrated in the ‘Lo7’ assembly (S-QTL) based on BLASTn sequence similarity searches as described by Hackauf et al. (2009)^121^. The *S* locus genomic region in rye was identified using orthologous gene models from *Brachypodium dystachion* including *Bradi2g35750*, that is predicted to encode a protein of unknown function *DUF247*^46^. Furthermore, we included the marker *SCM1* from Hackauf and Wehling (2002)^122^ in our analyses, that represents the rye ortholog of a thioredoxin-like protein linked to the *S* locus in the grass *Phalaris coerulescens*^123,124^. Likewise, the isozyme marker *Prx7* linked to the *S* locus in rye was investigated as described by Wricke and Wehling (1985)^43^. The *S* locus was mapped in a F2 population (n= 96), produced by crossing the self-incompatible variety ‘Volhova‘ with the self-fertile line No. 5 (‘l.5’), the latter of which carrying the mutation for self-fertility at the *S* locus on chromosome 1R (Voylokov et al. 1993, Fuong et al. 1993). Progeny from this cross are heterozygous for the self-fertility mutation. The gametic selection caused by self-incompatibility in such crosses was used for the mapping of *S* relative to markers (S-1RSTS) according to previously described protocols^45,125^. The SI mechanism prevents fertilization of all pollen grains except those carrying the *Sf* allele. As a consequence, only those 50% of the pollen grains carrying the mutation will be able to grow and fertilize upon self-pollination of a F1 hybrid from the cross. Therefore, the functional *S* allele results in distorted segregation of marker loci linked to the self-fertility mutation in the F2. The degree of segregation distortion depends on the recombination frequency *r* between the segregation distortion locus (SDL) and analyzed marker loci. For example, after selfing a F1 with the constitution *SM1/SfM2*, where *S* and *Sf* are active (wild type) and inactive (mutant) alleles of the self-incompatibility locus *S*, respectively, and M1 and M2 are alleles of a marker locus linked in coupling phase, the expected segregation ratio for the marker will be as follows^126^:

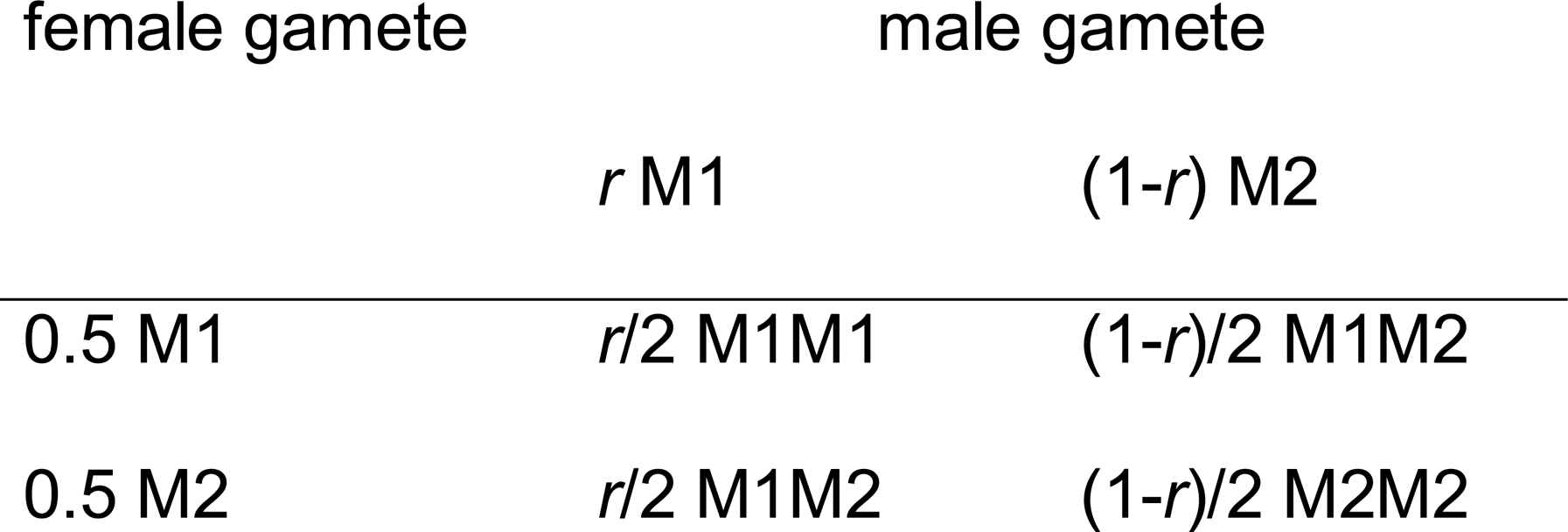

In case of *r*= 0 the frequency of heterozygous genotypes for the marker locus is equal to 0.5, and a significant excess of homozygous genotypes for the allele that originated from the self-fertile line (M22) is observed. Distorted segregation of marker loci were statistically analysed for mapping the *S* locus as outlined by Voylokov et al. (1998)^127^.

#### Genes affecting low temperature tolerance

The line ‘Puma-SK’ was produced by subjecting ‘Puma’ by recurrent selection under extreme cold winter conditions (−30 °C) to purify for the alleles contributing to increased cold tolerance. ‘Puma-SK’ was used in an intergeneric cross with the Canadian winter wheat cultivar ‘Norstar’, which generated a winter wheat introgression line (containing a segment of 5RL from ‘Puma’ (designated herein as ‘Norstar-5A5R’) that contained *Fr2*^128^.

To characterize the *Fr2* region in ‘Puma-SK’ and the introgression in ‘NorstarPuma5A:5R’, whole genome sequencing was performed using the Chromium 10X Genomics platform. Nuclei were isolated from 30 seedlings, and high molecular-weight genomic DNA was extracted from nuclei using phenol chloroform according to the protocol of Zheng et al. (2012)^129^. Genomic DNA was quantified by fluorometry using Qubit 2.0 Broad Range (Thermofisher) and size selection was performed to remove fragments smaller than 40 kbp using pulsed field electrophoresis on a Blue Pippin (Sage Science) according to the manufacturer’s specifications. Integrity and size of the size selected DNA were determined using a Tapestation 2200 (Agilent), and Qubit 2.0 Broad Range (Thermofisher), respectively. Library preparation was performed as per the 10X Genomics Genome Library protocol (https://support.10xgenomics.com/genome-exome/library-prep/doc/user-guide-chromium-genome-reagent-kit-v2-chemistry) and uniquely barcoded libraries were prepared and multiplexed for sequencing by Illumina HiSeq. De-multiplexing and the generation of fastq files was performed using LongRanger mkfastq (https://support.10xgenomics.com/genome-exome/software/pipelines/latest/using/mkfastq; default parameters).

Sequencing reads from ‘Puma-SK’ and ‘NorstarPuma5A:5R’ were aligned to the rye line ‘Lo7’ and bread wheat cv. Chinese Spring^9^ genome assemblies, respectively, using LongRanger WGS (https://support.10xgenomics.com/genome-exome/software/pipelines/latest/using/wgs; arguments - vcmode ‘freebayes’). Large scale structural variants detected by LongRanger were visualized with a combination of Loupe (https://support.10xgenomics.com/genome-exome/software/visualization/latest/what-is-loupe; tbl. S-DATAACCESS). Short variants were called using the Freebayes software (github.com/ekg/freebayes) implemented within the Longranger WGS pipeline. For determining the introgression, ‘NorstarPuma5A5R’ reads which did not map to the Chinese Spring reference were aligned to the ‘Lo7’ assembly using the LongRanger align pipeline (https://support.10xgenomics.com/genome-exome/software/pipelines/latest/advanced/other-pipelines). Samtools^114^ bedcov was used to calculate the genome-wide read coverage across both references. Copy number variation between ‘Puma-SK’ and ‘Lo7’ was detected using a combination of barcode coverage analysis output by the Longranger WGS pipeline, and read depth-of-coverage based analysis using CNVnator^130^ and cn.mops^131^.

To identify differentially expressed genes that may be contributing to the phenotypic differences in cold tolerance, ‘Puma-SK’ and ‘NorstarPuma5A:5R’ were grown and crown tissues harvested at different stages of cold acclimation. Both genotypes were grown for 14 days (d) at 20 °C with a 10 hour (h) day length. Plants were then treated to decreasing temperatures and daylengths over a 70d period, designed to mimic field conditions for winter growth habit. After the initial 14 d growth period, the temperature was reduced to 18 °C, then after 3 d (15 °C), 7 d (12 °C), 14 d (9 °C), 21 (6 °C), 28 d (3 °C), 35 d (2 °C), 42 d (2 °C), 49 d (2 °C), 56 d (2 °C), 63 d (2 °C), and 70 d (2 °C). In addition to adjusting the temperature, the day length was adjusted incrementally from 13.5 h at 0 d to 9.2 h at 70 d. Day length changes were programmed to occur on day 3 and day 4 of each week. For each change in temperature, crowns were sampled from two independent replicate plants for each genotype, which were used for analysis of gene expression by RNA sequencing. Crown tissue was sampled one hour after the lights came on in the morning to minimize circadian rhythm effects. In addition, at each change in temperature, five plants from each genotype were used to analyze the rate of plant phenological development (dissection of the plant crown to reveal shoot apex development) and cold hardiness during cold acclimation. Cold hardiness was determined using LT50 measurements, the temperature at which 50% of the plants are killed by LT stress, using the procedure outlined by Fowler et al. (2016)^72^.

Sequencing adapters were removed and low-quality reads were trimmed using Trimmomatic^132^. RNA reads from ‘NorstarPuma5A:5R’ and ‘Puma’ were aligned to the ‘Lo7’ reference using Hisat2^85^ (default arguments) and transcripts were quantified with htseq^133^. Differential expression analysis was carried out using DESeq2^134^ (default parameters).

### Data Availability

Data access information including raw sequence data, selected assembly visualisations, gene annotation, and optical map data, is tabulated in the table S-DATAACCESS.

## Acknowledgements

We thank the following for their valuable contributions: Manuela Knauft, Ines Walde, and Susanne Koenig, and Stefanie Thumm (IPK), Jennifer Ens (University of Saskatchewan), Cristobal Uauy and James Simmonds (John Innes Centre), Susan Duncan (Earlham Institute), Zdeňka Dubská, and Jitka Weiserová (Institute of Experimental Botany), Alex Hastie (Bionano Genomics), Kobi Baruch (NRGene), and Stefan Taudien (Universitätsmedizin Göttingen) provided technical, laboratory, and greenhouse services. Anne Fiebig, Jens Bauernfeind, Thomas Münch, and Heiko Miehe (IPK) provided IT services. Andreas Graner provided advice. Bionano optical maps were generated through funding The Czech Science Foundation (grant number: 17-17564S) via H. S., and the German Federal Ministry of Education and Research via Eva Bauer (grant number: 0315946A), who also contributed funding for WGS sequencing. KWS LOCHOW GMBH funded the CSS sequencing. A. L. received funding from the Agriculture and Agri-Food Canada International Collaboration Agri-Innovation Program. A. S. received funding from the Natural Resources Institute Finland (Luke) Innofood Stategic Funds program. A. Hall received funding from the Biotechnology and Biological Sciences Research Council Designing Future Wheat program (grant number: BB/P016855/1). K. F. X. M. received funding from the Bundesministerium für Bildung und Forschung (de.NBI, number 031A536) and from the Bundesministerium für Ernährung und Landwirtschaft (WHEATSEQ number 2819103915). D. S. received funding from HYBRO Saatzucht GmbH & Co. KG. J. D. received funding from the European Regional Development Fund’s plants as a tool for sustainable global development project (grant number: CZ.02.1.01/0.0/0.0/16_019/0000827). B. W. received funding from the 2Blades Foundation. B. H. and F. O. received funding from the Julius Kühn-Institute. V. K. received funding from KWS SAAT SE & Co. KGaA. A. Houben received funding from the Deutsche Forschungsgemeinschaft (grant number: HO 1779/30-1). U. S. received funding from the Bundesministerium für Bildung und Forschung (de.NBI, number FKZ 031A536). H. B. received funding from the Montana Wheat and Barley Committee. X-F. M. received funding from the Noble Research Institute, LLC. E. B. received funding from the Bundesministerium für Bildung und Forschung via the project “RYE-SELECT: Genome-based precision breeding strategies for rye” (grant number: 0315946A). I. S. and J. M. received funding from the Australian Research Council (grant number: CE140100008). C. J. P. received funding from Genome Canada and Genome Prairie (grant number: CTAG2). D. K. and A. S. received funding through the National Research Council Canada’s Wheat Flagship Program. D. B. F. received funding from the Province of Saskatchewan Agriculture Development Fund (ADF). B. K. received funding from the Bundesamt für Landwirtschaft, Bern (grant number: PGREL NN-0036). M. R-T., H. B-B., S. S., and B. M. received funding from the Polish National Science Centre (grant numbers: DEC-2015/19/B/NZ9/00921; DEC-2014/14/E/NZ9/00285; 2015/17/B/NZ9/01694).

## Author Contributions

*Project conception and consortium coordination*

N. S. (leader), K. F. X. M., M. M., V. T., N. R.

*Manuscript and main figures*

M. T. R-W. (leader), N. S., B. H., with input from all authors.

*Genome assembly and data integration*

M. T. R-W. (leader), M. M.

*Provision, curation, cultivation, and phenotyping of genetic resources*

A. B. (Secale diversity panel); V. K. (‘Lo7’); D. B. F., B. H., Q. L., C. J. P., B. B. (‘Norstar’, ‘Puma’); V. K., B. H., M. R-T., H. B-B., S. S., B. M. (Secale genome size estimation panel).

*Sequencing data [ADD SECTION FOR FUNDING SEQUENCING?]*

J. L., A. L., J. J., A. Hall, D. S., A. Himmelbach, S. P., A. H. S., J. P., B. B.

*Genome size estimation and chromosome flow sorting*

J. D., J. Č., J. V.

*Bionano optical map*

H. Š., H. T., E. B.

*FISH*

M. B., A. Houben.

*Gene annotation*

D. S., G. K., T. L., M. S., K. F. X. M., J. K.

*Repetitive element annotation and analysis*

H. G., T. W., M. S., K. F. X. M.

*miRNA annotation*

H. B., B. S.

*Secale diversity analysis*

M. S., M. M., H. S., U. S.

*Hi-C-based SV detection*

M. T. R-W. with input from M. M.

*Resistance gene identification and analysis*

B. S., B. W, B. H., B. K., C. P., T. W.

*SI and CMS gene identification and analysis*

B. H., I. S., J. M.

*Mapping of S- locus*

B. H., A. V. V., N.T.

*Wheat-rye introgression analysis*

J. P., L. G., M. T. R-W., M. M., with input from B. H.

*Low temperature tolerance analysis*

C. J. P., B. B., S. W.

