## Supplementary Notes and Figures for "Chromosome-scale genome assembly provides insights into rye biology, evolution, and agronomic potential"

Supplementary Materials

Rabanus-Wallace et al. 2019

Supplementary Note X. S-FLOWCYT: Genome size estimation by flow cytometry	3

Supplementary Note X. S-PSASS: Pseudomolecule assembly procedure	4

X.X Overview	4

Figure X.X. S-ASSOVER	5

X.X. Contigging and Scaffolding (step 1)	6

X.X. Scaffold chromosome assignments using genetic map, CSS, and TCC Hi-C	6

X.X. Guide Hi-C map construction (steps 3, 5, and 7)	7

X.X. Comments on manual AGP editing using the visualisation suite (steps 4, 6, and 8)	8

X.X. Manual chimera detection and breaking (step 4)	9

X.X. Superscaffolding (step 6)	9

X.X. Final order adjustment (step 8)	9

Supplementary Figure X.X. S-ASSEVO	10

Supplementary Figure X.X. S-CSSDEMO	11

Supplementary Figure X.X. S-HICDEMO	12

Supplementary Figure X.X S-MAPDEMO	13

Supplementary Figure X.X S-OPTDEMO	14

Supplementary Figure X.X. S-OPTRESDEMO	15

Supplementary Note X. S-ASSDATA: Assembly data generation and processing	16

X.X. Paired-end and mate paired read sequencing	16

X.X. 10X Chromium linked read sequencing	16

X.X. Hi-C sequencing	17

X.X. Chromosome Sorted Shotgun (CSS) sequence	17

X.X. Bionano optical map generation and alignment to assembly scaffolds	18

X.X. Lifting of genetic marker positions from Bauer et al. contigs	19

Supplementary Figure X.X. S-HICDIST	20

Supplementary Figure X.X. S-10XHIST1	21

Supplementary Figure X.X. S-10XHIST2	22

Supplementary Note X. S-REP. Investigations into the repetitive genome	23

Supplementary Figure X. S-RPT_ASSCMP	24

Supplementary Figure X. S-SATS	25

Supplementary Figure X. S-SATSFISH	26

Supplementary Figure X.X. S-KMERREP	28

Supplementary Figure X.X. S-TEMAINPROF	29

Supplementary Figure X.X. S-TETERMPROF	30

Supplementary Figure X.X. S-SUBRATE	31

Supplementary Figure X.X. S-RECOMB	32

Supplementary Figure X.X. S-TETRANSLOC	33

Supplementary Note X. S-TEEXP: Arguments on the timing of TE superfamily expansions in various Triticeae	34

Supplementary Note X. S-SV: Secale diversity and segregating structural variations	35

Supplementary Figure X.X. S-HICSV	35

Supplementary Figure X.X. S-HICPERM	36

Supplementary Note X. S-COLIN	37

Supplementary Figure X.X. S-COLIN_lg1	37

Supplementary Figure X.X. S-COLIN_lg2	38

Supplementary Figure X.X. S-COLIN_lg3	39

Supplementary Figure X.X. S-COLIN_lg4	40

Supplementary Figure X.X. S-COLIN_lg5	41

Supplementary Figure X.X. S-COLIN_lg6	42

Supplementary Figure X.X. S-COLIN_lg7	43

Supplementary Note X. S-OUTIN	44

Supplementary Figure X.X.	44

Supplementary Note X: S-RFMULTI	45

Supplementary Figure X.X. S-RFMULTI	45

Supplementary Note X. S-NLR: Comparative cluster arrangement and phylogenetics of resistance genes	46

X.X Phylogenetic and homology-based investigations of resistance gene orthologs	46

Supplementary Figure X.X. S-NLRPHYLO	48

Supplementary Figure X.X. S-PM2PHYLO	49

Supplementary Figure X.X. S-PM2COMP	50

Supplementary Figure X.X. S-PM2EVO	51

Supplementary Figure X.X. S-PM3PHYLO	52

Supplementary Figure X.X. S-PM3COMP	53

Supplementary Figure X.X. S-MLAPHYLO	54

Supplementary Figure X.X. S-MLACOMP	55

Supplementary Figure X.X. S-LR10CMP	56

X.X Phylogenetic and homology-based investigations of low temperature tolerance gene homologs	57

Supplementary Figure X.X. S-CBFPHYLO	58

Supplementary Note X. S-COLD: Experimental investigations of low temperature tolerance.	59

Supplementary Figure X.X. S-LT50EXPT	60

Supplementary Figure X.X. S-VRN1EXPN	61

Supplementary Note X. S-INTROG.	62

Supplementary Figure X.X. S-CLASSCASES	63

Supplementary Figure X.X. S-KSRECOMB	64

Supplementary Figure X.X. S-1RSCOMMON	65

Supplementary Note X. S-PUBLIC	66

Supplementary Figure X.X S-PUBLICTRIAL	66

Supplementary References	68

### Supplementary Note X. S-FLOWCYT: Genome size estimation by flow cytometry

We characterised the landscape of S. cereale genome sizes in order to contextualise the size of the ‘Lo7’ genome and gain an impression of genome size variation within the species.

Grains from fifteen diverse rye accessions were provided by nine providers listed in table S-FLOWCYT. Plants of pea served as an internal reference standard in flow cytometric estimation of nuclear DNA content in all accessions, except of the tetraploid accession ACE-1, for which *S. cereale* line ‘Lo7’ was used as a reference. Seeds of pea (*Pisum sativum* cv. Ctirad) were obtained from Semo (Smržice, Czech Republic) breeding station. Plants were raised in garden compost in pots and maintained in a greenhouse until they reached a height of 10–20 cm.

Nuclear genome size was estimated as described by Doležel et al. (2007)^1^. Briefly, 10 mg of fresh leaf tissue of each of the rye accessions and the reference standard were chopped together in a 1 mL volume of LB01 solution^2^ using a razor blade. The resulting homogenate was filtered through a 50-µm nylon mesh. The filtrate was made up to 50 µg/mL propidium iodide and 50 µg/mL RNase, and subjected to flow cytometry using a CyFlow Space flow cytometer (Sysmex Partec GmbH, Görlitz, Germany) equipped with a 532 nm green laser. The gain of the instrument was adjusted so that the peak representing G1 nuclei of the standard was positioned approximately on channel 100 on a histogram of relative fluorescence intensity when using a 512-channel scale. Five individual plants per each test species were sampled, and each sample was analyzed three times, each time on a different day. A minimum of 5000 nuclei per sample was analyzed and 2C DNA contents (in pg) were calculated from the means of the G1 peak positions by applying the following formula:

$$2C nuclear DNA content\frac{= sample G1 peak mean \times standard 2C DNA content}{standard G1 peak mean}$$

Mean nuclear DNA content (2C) was then calculated for each accession. DNA contents in pg were converted to genome size in bp using the conversion factor 1 pg DNA = 0.978 Gbp^3^. Statistical analysis was performed using NCSS 97 statistical software package (Statistical Solutions Ltd.). One-way ANOVA and a Bonferroni's (All-Pairwise) multiple comparison test were used for analysis of variation in monoploid (1Cx) genome size. A significance level α = 0.01 was used.

### Supplementary Note X. S-PSASS: Pseudomolecule assembly procedure

#### X.X Overview

The general assembly workflow is depicted in figure S-ASSOVER. In step 1, raw scaffolds were assembled using NRGene’s DeNovoMAGIC3.0 pipeline^4-7^, using sequence data from paired-end (PE) shotgun, mate-paired (MP), and 10X chromium molecule-linked (ML) reads. Scaffolds were enriched with genetic positions and chromosome assignments by mapping the contigs of a draft rye genome assembly of Bauer et al. (2017)^8^ to the new assembly scaffolds, and lifting this information from one assembly to the other. In step 2, misjoined (‘chimeric’) scaffolds were detected and broken using two different automatic approaches. Scaffold arrangement into pseudomolecules (steps 3 to 8) was primarily performed in the R statistical environment exploiting in particular the packages ‘data.table’ and ‘ggplot2’. The procedure focussed on manual curation, beginning by manually breaking any potential chimeras that were not detected automatically (step 4), then on concatenating adjacent scaffolds into superscaffolds (step 6), and finally on arranging the superscaffolds into pseudomolecules (step 8). Preceding each of these three manual steps, a candidate scaffold order was generated with the aid of Hi-C link frequency information (steps 3, 5, and 7). The candidate scaffold orders functioned each time as the basis of a suite of data (primarily Hi-C, optical map, genetic map, and CSS) visualisations designed to give the user an intuitive impression of a candidate’s accuracy, at the different resolutions offered by the various datasets. Summary statistics describing the assembly at progressive points through the assembly procedure are detailed in supplementary table S-ASSSTATS.

#### Figure X.X. S-ASSOVER

Assembly procedure overview. Coloured circles represent data sources used in each of the steps. Black=Mate paired and paired end read data; Pale blue=10X Chromium linked reads; Orange=Hi-C data (‘Lo7’, TCC method with HindIII digest); Dark blue=BioNano optical map alignment; Green=Genetic map positions lifted from Bauer et al. (2017)^8^ assembly contigs. Yellow=CSS reads.

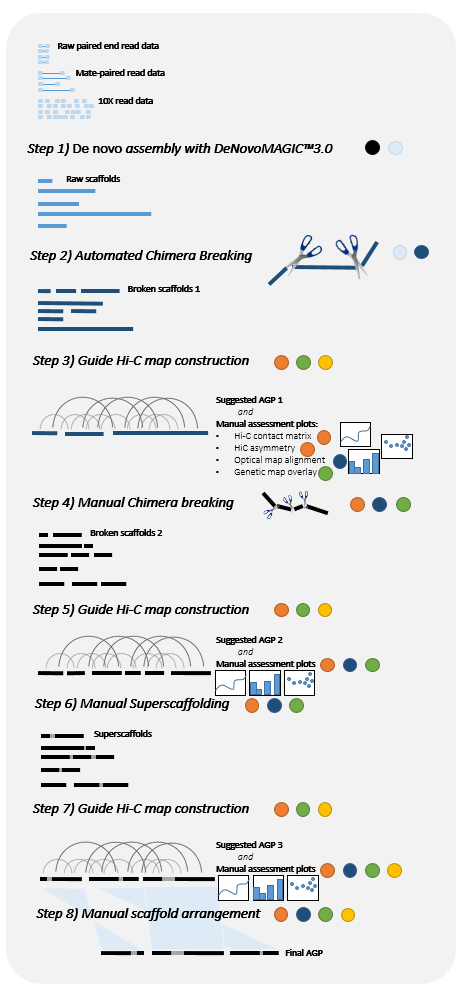

#### X.X. Contigging and Scaffolding (step 1)

*De novo* assembly was performed using NRGene’s proprietary DeNovoMAGIC3.0 pipeline, as previously described^4-7^. Briefly, the procedure can be divided into the following steps: 1) Data pre-processing and quality control. Adapter sequences are trimmed and overlapping PE reads are merged. Reads containing probable sequencing errors are purged by removing any reads that contain unique or near-unique sequences; 2) Contig assembly. A 127mer De Bruijn graph assembly is constructed from the contigs present in all PE and MP reads. The PE and ML reads are then used to identify and validate likely paths through the graph; 3) Scaffolding. PE and ML reads are used to identify/validate scaffold-to-scaffold joins, and to estimate the distances between them; 4) Gap filling. The PE and MP data are employed to identify unique De Bruijn graph paths that fill intra-scaffold gaps; 5) Scaffold validation with ML reads. A window-based approach is used to determine whether the mapped positions of ML reads supports the contiguity of the scaffolds. Scaffolds are broken wherever the number of inferred molecules overlapping edges of a 20 kb window is judged inconsistent with the number spanning the whole window; 6) Scaffold concatenation with ML reads. A graph is constructed linking terminal bins containing multiple shared ML read barcodes. The scaffolds in any group forming a linear path through the graph are joined together to form a single scaffold. The assembly produced 107,580 scaffolds with N50 length >22 Mbp, and a combined length of 6,724,307,309 bp (tbl. S-ASSSTATS).

#### X.X. Scaffold chromosome assignments using genetic map, CSS, and TCC Hi-C

Following the methods described in Beier et al. (2017)^9^, each scaffold was also assigned to a chromosome based on the genetic map, and also given a CSS-based chromosome assignment, wherever the map markers/CSS bins associated with the scaffold showed a clear majority preference for one chromosome. Scaffolds for which the CSS and map assignments agree were then used as the basis for a Hi-C-based chromosome assignment: Hi-C links between chromosome-assigned scaffolds and the focal scaffold were counted, and a Hi-C-based assignment awarded if the links overwhelmingly associated the scaffold with other scaffolds from one particular chromosome (details in Beier et al., 2017)^9^.

The acquisition/generation and processing of the Hi-C, genetic map, and CSS data are described in supplementary note S-ASSDATA.

Automated chimera detection and breaking (step 2)

The initial set of scaffolds (“scaffolds_v1” in figure. S-ASSOVER) were broken wherever chimeric breakpoints were suspected. Two automated methods of chimera detection were used. Firstly, a group of scaffold breakpoints were suggested by the Bionano assembly software, which flags conflicts between the optical map alignments (henceforth *optigs*—*op*tical con*tigs*) and the scaffolds, and a report is returned suggesting where either the scaffolds or the optigs aligned to them should be broken to resolve the conflict. For each conflict, choice between breaking the optig or the scaffold is based on the level of support for the optig’s contiguity provided by the raw optical molecule data (<https://bionanogenomics.com/support-page/bionano-solve/>).

At suspected breakpoints, the scaffold was split into three: two main fragments, and a small fragment of length 20 kbp surrounding the nominated breakpoint. This precaution aims to ensure the real breakpoint—which is expected to differ slightly from the nominated breakpoints owing to estimation inaccuracy—is contained in a short scaffold, and subsequently relegated to the ‘unassigned’ chromosome chrUn.

The second automated breakpoint detection method made use of 10X Chromium molecule-linked read data based on the expectation that the absence of molecules spanning breakpoints should result in a sharp decrease of molecule coverage near breakpoints. The method is part of the TRITEX pipeline described in Monat et al. (2019)^10^, and the TRITEX R function break_10x() was run with the parameters ratio=-3, interval=5e4, minNbin=100, and dist=1e4. Breaking the scaffolds at the points detected by these methods yielded the second scaffold set (“scaffolds_v2” in figure. S-ASSOVER).

#### X.X. Guide Hi-C map construction (steps 3, 5, and 7)

The first of several suggested optimal scaffold arrangements (“A golden path”; AGP) was constructed for the v2 scaffolds (“scaffolds_v2” in figure. S-ASSOVER) using the Hi-C-based method described by Burton et al. (2013)^11^, using the implementation described in Beier et al. (2017)^9^. Briefly, a graph structure is constructed for each chromosome, with vertices representing scaffolds, and weighted edges representing the frequency of Hi-C links between them. After removing any edges that conflict with available genetic map information, Prim’s algorithm is used to yield a minimum spanning tree, the longest path through which functions as a “backbone” arrangement of putatively-ordered scaffolds. Scaffolds not included in the backbone are then inserted wherever their inclusion incurs the least additional weight. The total weight of this candidate order is then further minimised by inducing local perturbations using the K-opt and node relocation heuristics (with k=2) and accepting any improvements, as described by Wu et al. (2008)^12^.

#### X.X. Comments on manual AGP editing using the visualisation suite (steps 4, 6, and 8)

The uses of the different data visualisations in manual scaffold arrangement editing are described in this section. Some methodological details are given in supplementary note S-ASSDATA, and examples are given in figs ASSEVO--S-OPTRESDEMO. The visualisations as produced for the final assembly are available as public data (see table S-DATAACCESS). The evolution of the assembly through two rounds of manual editing are visualised in figure S-ASSEVO.

The first tentative AGP produced (“AGP_v1” in figure. S-ASSOVER) was used as the basis for a first suite of data visualisations for manual curation. It was noted at this stage that short scaffolds (< 500 kb in length) usually clustered into groups, which, while accounting for only a small proportion of the total assembly, had a strongly adverse effect on the contiguity of the assembly, manifesting most starkly as heavy vertical striations through the Hi-C contact matrices. These inaccuracies could be improved by excluding short scaffolds (at the expense of assembly completeness), but this detracted from the ability to rescue some short scaffolds by manually incorporating them into superscaffolds (in step 6). It was decided therefore to include all chromosome-anchored scaffolds in the pseudomolecules until step 7.

CSS chromosome assignment plots (fig. S-CSSDEMO) are simply stacked histograms showing the numbers of reads from CSS libraries deriving from each chromosome that map within bins across each chromosome (see supplementary note S-ASSDATA). These were used to manually identify obvious inter-chromosomal chimeras not identified in the automated chimera breaking steps (step 4).

Hi-C contact plots represent the frequency of Hi-C links between parts of the genome as a heatmap^13^. Briefly, the links between restriction fragments falling within 1 Mbp bins are counted, and a normalisation is performed to account for known biases introduced by several factors including the number of restriction fragments in the bin and the GC content surrounding these sites^14^. The counts in the resulting matrix are represented as a heatmap. These plots show discontinuities that can reflect inversions and badly-ordered scaffolds^11^ though scaffold boundaries and repetitive sequence that affects the mapping rate may also affect the continuity.

Hi-C asymmetry plots^15^ (fig. S-HICDEMO) are derived from the same data as Hi-C contact plots. These show the ratio between the numbers of Hi-C links connecting bins to the left and right respectively of each bin along each chromosome, and are particularly useful for revealing inverted scaffolds.

The genetic positions of markers superimposed upon the AGP allows rapid detection of larger-scale inversions and misplaced scaffolds (fig. S-MAPDEMO), which we often found to provide information at a finer scale than was visible in the Hi-C asymmetry plots. Conversely, the Hi-C asymmetry plots easily reveal ordering errors in the pericentromeric regions where the genetic map shows little change owing to low recombination.

Alignment of the optigs to the assembly allows validation of the internal sequence accuracy of the scaffolds, and—wherever an optig contains matches to several scaffolds—assessment or revision of the suggested order and orientation (fig. S-OPTDEMO). The optical map can be used to join neighbouring scaffolds to create superscaffolds, given all other sources of data support (or at least fail to contradict) the join.

#### X.X. Manual chimera detection and breaking (step 4)

Sixteen chimeric scaffold breakpoints were manually detected by close inspection of the visual suite over the first suggested AGP (“AGP_v1” in figure. S-ASSOVER). Where suspected breakpoints coincided with the comparatively small gaps (usually 5-20 kb) between sequential alignments to different optigs, the break was placed at the midpoint of these gaps. The scaffolds were broken with the same procedure as described for step 2, to yield the scaffolds_v3 scaffold set.

A second Hi-C guide map (AGP_2) was then constructed from the broken scaffolds (step 5).

#### X.X. Superscaffolding (step 6)

The visual suite was manually assessed to identify candidate scaffold joins for the creation of superscaffolds. This process primarily utilised the optical map visualisation (fig. S-OPTDEMO), with each candidate being then checked against the other visualisations to assure consistency. Superscaffolds were created only where optical map contigs and where the scaffold join A) was not contested or made ambiguous by any optig alignments, and B) did not incur any ‘trade-offs’ between data sets (for instance, if the optical map strongly suggested a join between two nearby scaffolds, but to do so would incur a small incongruity in the linear order of marker positions along these scaffolds, then that join was not made).

The scaffold set including superscaffolds (“scaffolds_v4” in figure S-ASSOVER) was then used to produce a third Hi-C guide map, (step 7; “AGP_3” in figure S-ASSOVER). In the construction of AGP_3 the more stringent scaffold length limit was imposed as discussed above, resulting in a number of short scaffolds with highly uncertain placements (and which caused notable incongruity in the Hi-C contact matrix) being relegated to the unknown chromosome.

#### X.X. Final order adjustment (step 8)

A series of small adjustments to AGP_3 were tried by comparing the visual suite (in particular, the Hi-C asymmetry plots and contact matrices, and the genetic map) before and after making changes, and accepting those judged to improve the assembly, and which incurred no contradictions between datasets.

#### Supplementary Figure X.X. S-ASSEVO

Evolution of the scaffold arrangements in pseudomolecules imposed during manual editing steps. Chromosomes are arranged vertically in rows, with Mbp positions given on the horizontal-axis. Common sequences are linked between steps by green and blue blocks. The evolution of the order of sequences in each chromosome’s pseudomolecule progresses from bottom to top beginning with the Hi-C-based suggested order from step 5 (lowest), to the suggested order following superscaffold construction (middle), to the final order (highest). Red triangles represent sequences lost from the pseudomolecules from one step to another (i.e. which were either automatically assigned to chrUn owing to unresolved automated chromosome assignment, or excluded by minimum scaffold length cutoffs), and blue triangles represent sequence manually added (or re-added) to the pseudomolecules in the final step.

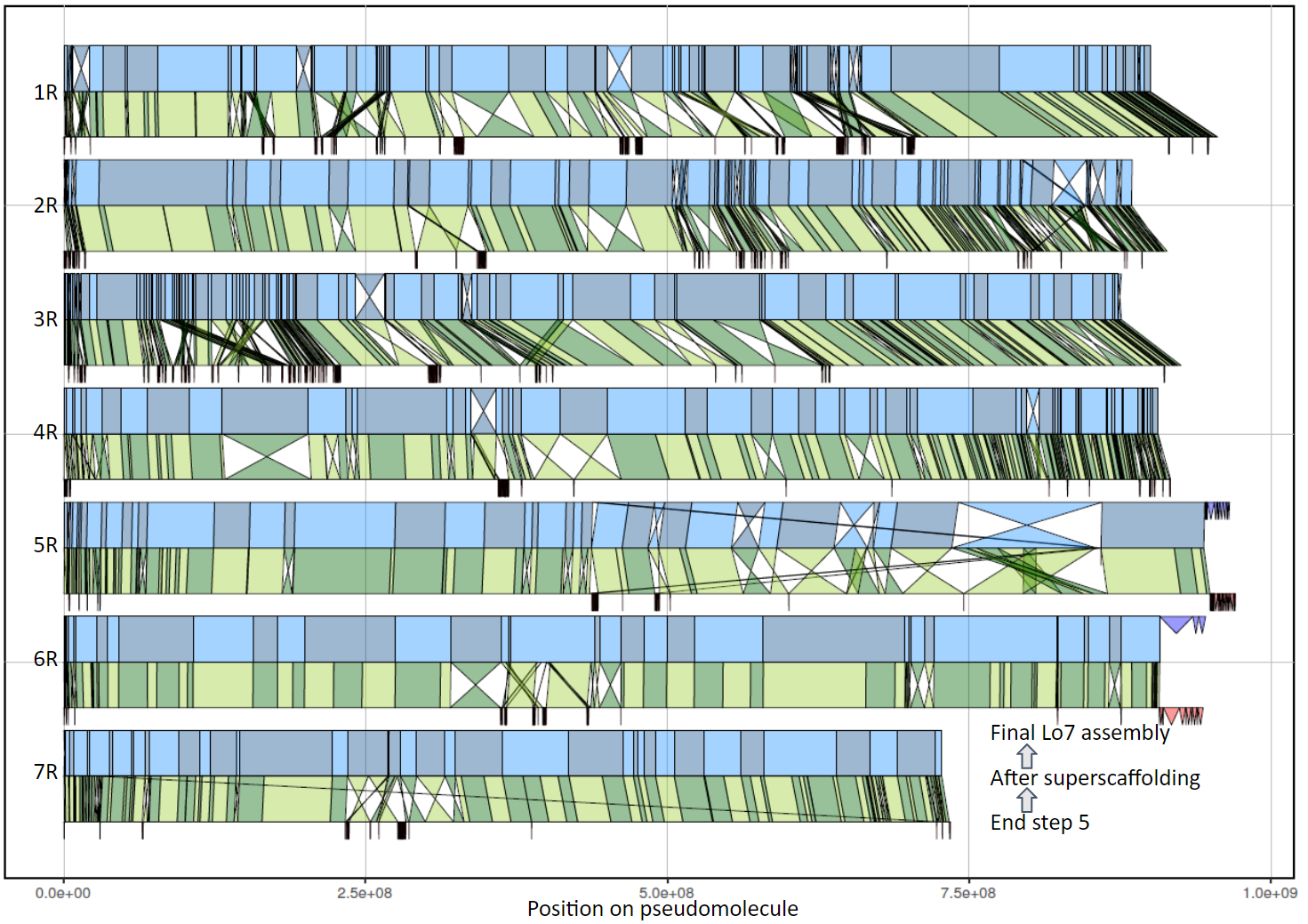

#### Supplementary Figure X.X. S-CSSDEMO

Example of an inter-chromosomal chimera, readily identifiable using CSS read depths displayed as a stacked histogram. Binned CSS read counts show that scaffold 2478 (v1 scaffolds) is a likely chimera between regions from chromosomes 3R and 5R, with the breakpoint near bin 50.

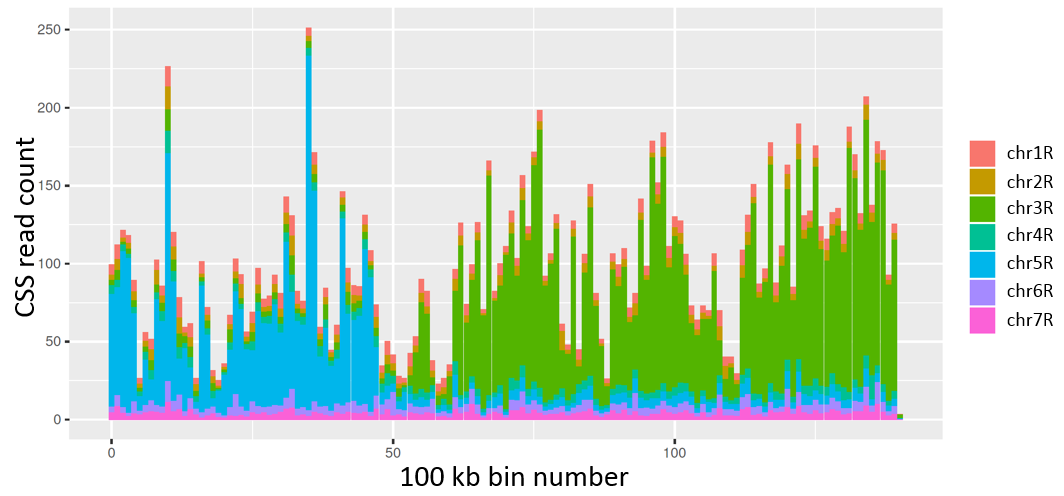

#### Supplementary Figure X.X. S-HICDEMO

An example of Hi-C data, represented with an asymmetry plot, used as an aid for manual assembly curation. This plot is over 1 Mbp bins on chromosome 6R, showing evolution between steps 5 (top panel) and the final assembly (bottom panel). Stark diagonals and discontinuities in the step 5 asymmetry plot reveal errors in scaffold order and orientation later corrected. The discernible break in continuity in the final asymmetry plot marks the centromere. Vertical bars represent scaffold boundaries.

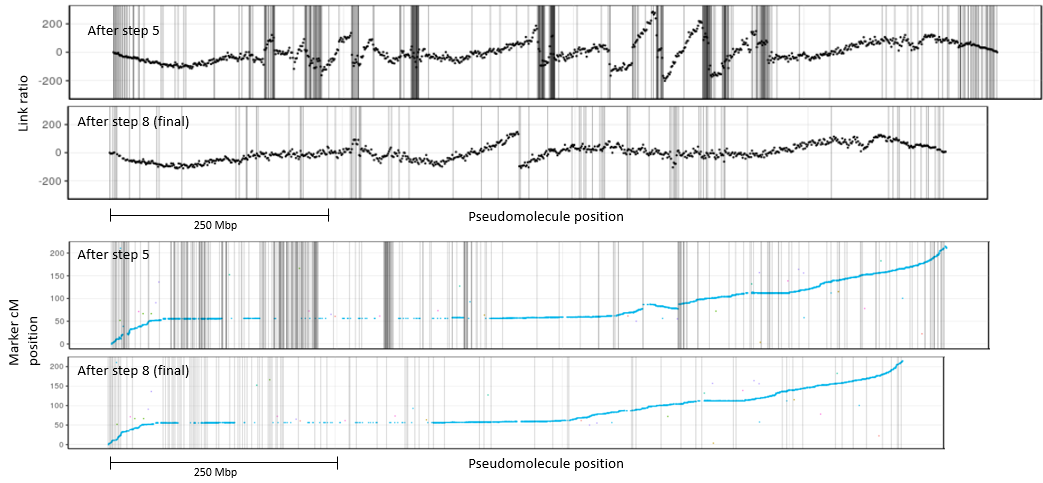

#### Supplementary Figure X.X S-MAPDEMO

An example of genetic map marker positions used as an aid for manual assembly curation. Genetic map positions of markers (dots) superimposed onto chromosome 5R, showing evolution between steps 5 (top panel) and the final assembly (bottom panel). An inverted scaffold can be seen in the step 5 plot around 600 Mbp from the 5’ end. Vertical bars represent scaffold boundaries.

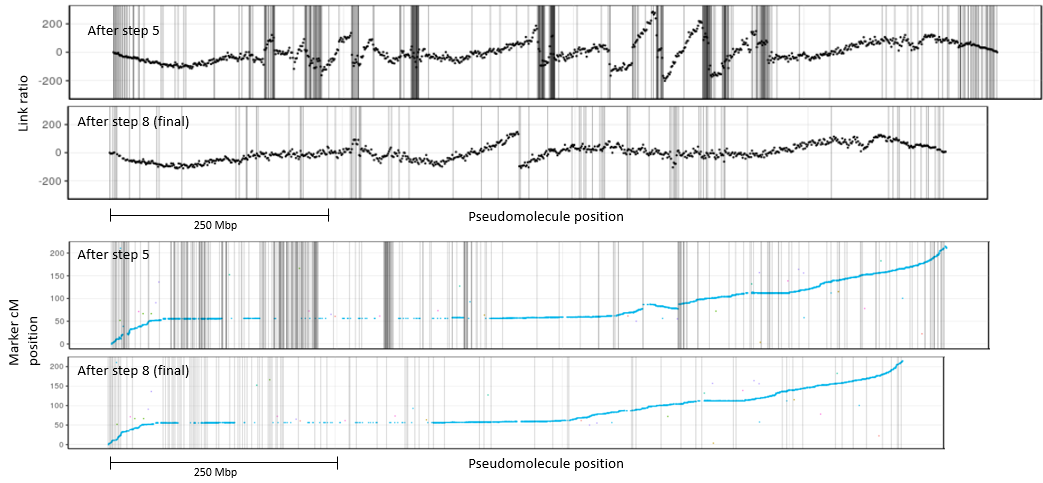

#### Supplementary Figure X.X S-OPTDEMO

Visualisation of optical map alignments aids manual AGP editing. An excerpted portion of the optical map alignment is shown from the short arm of chromosome 1R at step 5 in the assembly process. The horizontal red bar represents scaffolds (boundaries marked with black vertical bars), arranged into putative pseudomolecules. Black horizontal bars show aligned optigs, with coloured lines linking labels to their aligned positions on the scaffolds. The directions of the optigs are arbitrary. Colours are used to distinguish distinct alignment sections. Green boxes represent contiguous scaffold groups which, as the optical map alignments suggest, could be joined to form superscaffolds. Blue arrows mark changes in scaffold orientation that the optical map alignments suggest will improve the scaffold orientation.

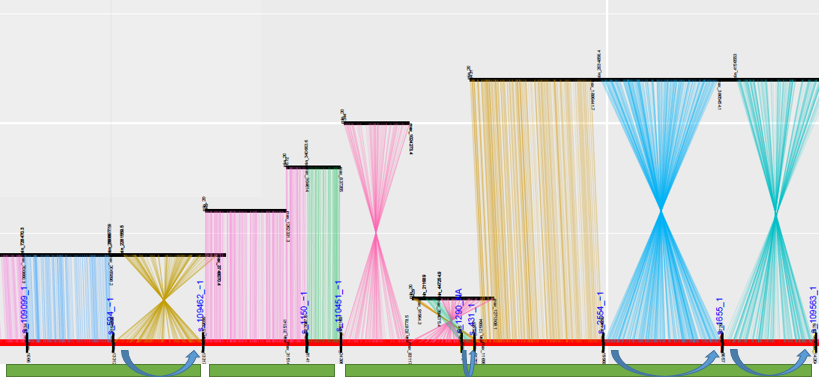

#### Supplementary Figure X.X. S-OPTRESDEMO

Resolution of the centromeric region of chromosome 7R (~250—300 Mbp) showing how adjustments to scaffold order improves agreement with the optical map. The optical map is displayed as described in figure S-OPTDEMO (with colours greyscale), and the changes in order displayed as in figure S-ASSEVO, with the subtending arrangement representing the result after step 5 (see figure S-ASSOVER), and the overlying arrangement representing the final assembly.

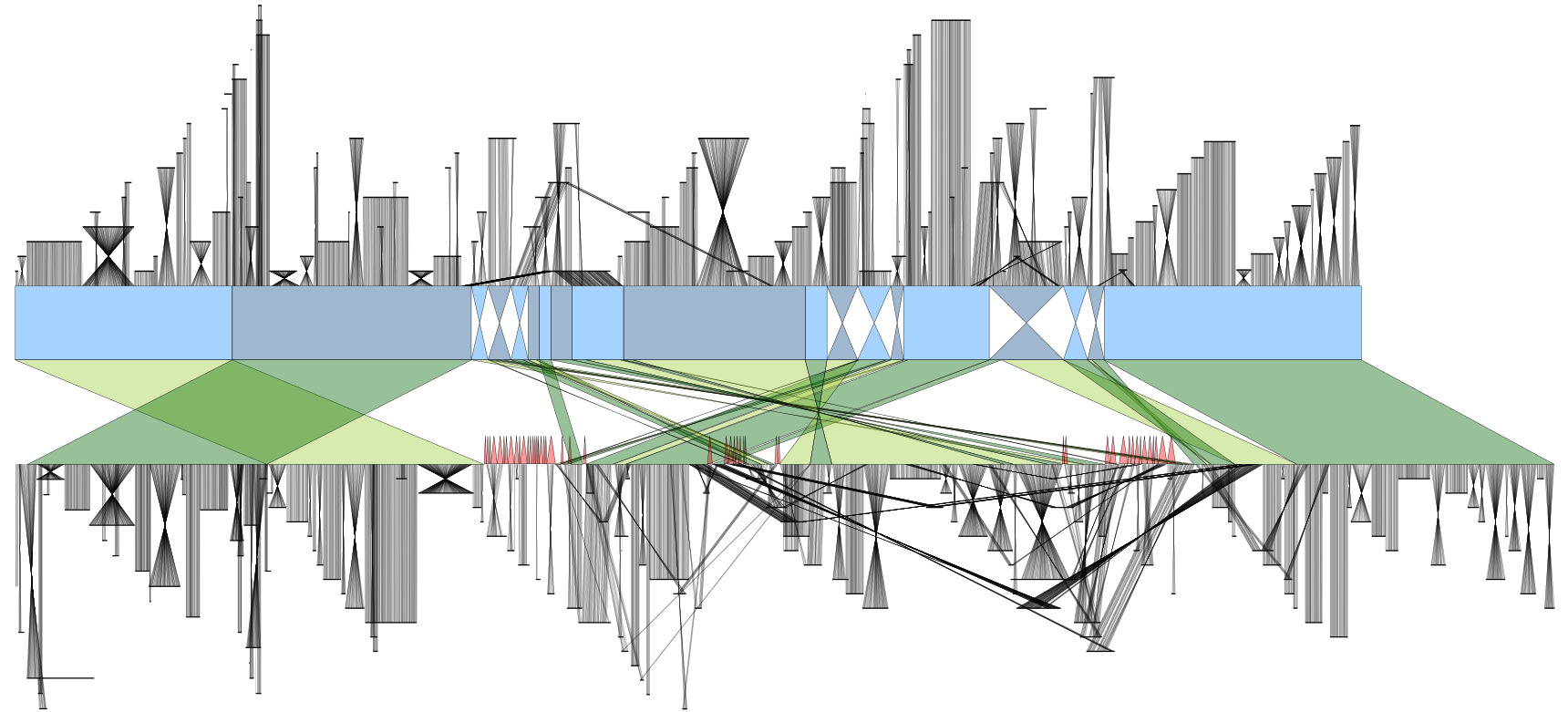

### Supplementary Note X. S-ASSDATA: Assembly data generation and processing

We describe here the generation and primary processing of datasets used to assemble the genome. Datasets generated for other analyses are described in the online methods. Tabulated descriptive statistics and quality control information are given in tables S-HTSLIBSTAT—S-OPTSTAT, and figures S-X.X—S-X.X. Access information for all data is given in table S-DATAACCESS.

#### X.X. Paired-end and mate paired read sequencing

DNA was isolated at IPK Gatersleben from fresh leaf tissue of Secale cereale inbred line ‘Lo7’ provided by KWS Lochow GMBH using the high molecular weight (HMW) phenol-chloroform protocol described by Dvorak et al. (1988)^16^. The DNA was provided to NRGene Israel for sequencing library construction using in-house implementations of standard Illumina protocols. The TruSeq protocol for PCR-free shotgun paired end reads (<https://www.illumina.com/documents/products/datasheets/datasheet_truseq_dna_pcr_free_sample_prep.pdf>) was used to produce four paired end libraries with two distinct insert size ranges (tbl. S-HTSLIBSTAT). The Nextera Gel-Plus protocol (<https://www.illumina.com/content/dam/illumina-marketing/documents/products/datasheets/datasheet_nextera_mate_pair.pdf>) was used to produce six mate paired libraries with three insert size ranges (tbl. S-HTSLIBSTAT). All libraries were sequenced on Illumina platforms to achieve between 36X and 88X approximate coverage of an estimated 7.9 Gbp genome (tbl. S-HTSLIBSTAT). Data primary processing a QC were performed prior to assembly as part of NRGene’s DeNovoMagic3.0 assembly pipeline.

#### X.X. 10X Chromium linked read sequencing

The HMW DNA was quantified by fluorometry using Qubit 2.0 Broad Range (Thermofisher) and size selection was performed to remove fragments smaller than 40 kb using pulsed field electrophoresis on a Blue Pippin (Sage Science) according to the manufacturer's specifications. Final DNA integrity and size were determined using a Tapestation 2200 (Agilent), and Qubit 2.0 Broad Range (Thermofisher), respectively. Library preparation was performed as per the 10X Genomics Genome Library protocol (10x Genomics) and uniquely barcoded libraries were prepared and multiplexed for Illumina sequencing. Three lanes of data were produced on the HiSeq2500 (paired end 150 bp, v2 chemistry), and an additional lane on the HiSeqX (paired end 150 bp). Reads were processed and molecule positions, lengths, coverage, and candidate chimera breakpoints were inferred using house R scripts based on those now implemented in the TRITEX pipeline^10^(figs. S-10XHIST1 & S-10XHIST2).

#### X.X. Hi-C sequencing

Leaf material (~1.5 g) was collected from one-week-old seedlings of lines ‘Lo7’ and ‘Lo225’ (accessions provided by KWS Lochow GMBH), and three wild rye gene bank accessions from the IPK Gatersleben, including *Secale vavilovii* (GBIS ID: R 1003), *Secale strictum* (R 2446), and *Secale sylvestre* (R 925). The ‘Lo7’ libraries used in pseudomolecule construction were prepared using the Tethered Conformation Capture (TCC) protocol detailed in Himmelbach et al. (2018)^15^, digesting with the restriction enzyme *Hind*III (recognition site AAGCTT). All samples were subsequently used to construct Hi-C libraries using the DpnII-restricted in situ Hi-C method described in Padmarasu et al. (2019)^17^. Hi-C libraries were size selected on SYBR Gold-stained gels to isolate fragments in the size range of 250—500 bp. The presence of the desired construct was assayed by digesting the library with *Cla*I and visually observing the shift in size profile to low molecular weight fragments compared to undigested control library. The libraries were sequenced on one lane of the HiSeq 2500 platform in high output mode (2 x 100 bp).

The libraries were trimmed at the appropriate Hi-C linker sequence (*Hind*III: AAGCTAGCTT; *Dpn*II: GATCGATC) using bbduk (<https://github.com/BioInfoTools/BBMap>; options ktrim=r, k=8, mink=2, qtrim=rl, trimq=20). The *Hind*III-digested TCC library used for pseudomolecule construction was mapped to the NRGene scaffolds, allowing an in silico digest of the scaffolds with the *Hind*III recognition sequence and subsequent identification of read pairs representing valid Hi-C links between restriction fragments, as described in Beier et al. (2017)^9^ based on Burton et al. (2015)^11^. Validly-mapped read pairs that do not show an arrangement suggesting they originated from the linking of restriction fragments are marked as “paired end” reads during this step (tbl. S-HICSTAT; fig. S-HICDIST), and are treated as contaminants and therefore removed from further analysis. Mapping and valid pair identification were performed identically for the *Dpn*II-digested in situ Hi-C libraries, except these were mapped directly to the pseudomolecule sequences, in silico digested with the *Dpn*II recognition sequence. Standard mapping and QC statistics were produced and inspected to confirm the libraries are of acceptable quality (tbl. S-HICSTAT; fig. S-HICDIST).

#### X.X. Chromosome Sorted Shotgun (CSS) sequence

DNA was isolated from preparations of individual rye chromosomes by chromosome flow-sorting as described earlier^18^, and prepared 500 bp insert Illumina sequencing libraries from each, using the manufacturer’s protocol, for sequencing on the GAIIx platform (paired end 2 x 150 bp; see tbl. S-CSSSTAT). Adapter contamination was removed using cutadapt^19^ (flags -a AGATCGGAAGAGC -A AGATCGGAAGAGC -m 30), and mapped to the ‘Lo7’ assembly scaffolds using minimap2^20^ (default parameters, short read algorithm -ax sr; tbl. S-CSSSTAT). Mapped read pairs (filtered using samtools^21^ view, flags -q30 -F260) were counted for 100 kbp non-overlapping bins across each scaffold, and normalised to reads-per-million (rpm) for each chromosome. Each bin was assigned to the chromosome with the highest rpm. A great deal of stochasticity occurs across scaffolds, probably owing in part to contamination of chromosome preparations with small amounts of unintended chromosomes, and also to similar repetitive sequences occurring in multiple chromosomes. To help smooth out some of this stochasticity, the bin calls were smoothed using a hidden markov model (HMM), treating a bin’s chromosome assignment as observed states, and the true chromosome as a hidden state. The model parameters were inferred using the Baum-Welch algorithm, as implemented by the R package `mhsmm`^22^, training on contigs longer than 10 Mbp, with the initial emission probabilities set up with a 100 times greater probability of bin assigned to chromosome X emitting chromosome X, in order force convergence towards the correct observation/emission correspondence. Similarly, transitions were initialised with 100 times greater probabilities for states remaining the same than changing. Initial state probabilities were made equal. The model converged over 33 iterations. The Viterbi algorithm was then used to infer the most likely bin states from the trained model.

#### X.X. Bionano optical map generation and alignment to assembly scaffolds

Long-range scaffolding of the genome sequence was supported by an optical map constructed from the ‘Lo7’ inbred line. HMW DNA was prepared from 10.5 million mitotic chromosomes (~22 µg DNA) purified by flow cytometry according to the methods of Kubaláková et al. (2003)^18^ and Šimková et al. (2003)^23^. The HMW DNA was labelled using NLRS DNA Labeling Kit (Bionano Genomics) at Nt.BspQI sites (GCTCTTC motif) and analyzed on the Irys platform (Bionano Genomics). A total of 2570.5 Gbp single molecule data > 150 kbp, corresponding to 347 ‘Lo7’ genome equivalents, was collected from nine Irys chips. The single molecule data were used to de novo assemble a map of the rye genome ‘Lo7’. A total of 2.57 Tbp of single molecule data with a length N50 of 217 kbp was used in a de novo assembly using Bionano Solve 3.1. Standard parameters for Irys data were used without “extend and split” and without haplotype refinement in order to create a single map for each allele with the parameter file “optArguments_nonhaplotype_noES_irys.xml.” In brief, the *de novo* assembly was accomplished by generating an assembly graph from alignments, satisfying a pValue threshold of 1e-10, finding during a pairwise comparison of all of the molecules, followed by refinement of label positions and extension of the ends of the maps based on molecules aligned to the maps with a pValue threshold of 1e-11. After five rounds of extension and refinement, a final refinement was conducted with a pValue threshold of 1e-15, a *de novo* assembly map was produced with a total length of 6,660 Gbp and a map N50 of 1.671 Mbp (tbl. S-OPTSTAT). The *de novo* assembly was used for scaffolding the sequence assembly (rye_Lo7_NRGene_scaffolds.fasta) using standard hybrid scaffold settings and selecting “resolve conflicts” for sequences and maps. Sequence contigs were *in silico* digested based on the recognition sequence for Nt.BspQI. The pattern of motif sites on the *in silico* map could be anchored to Bionano maps by comparing patterns that matched with a pValue threshold of 1e-10 using Bionano Solve 3.1. Chimera detection was accomplished by aligning contig maps to Bionano maps with pValue threshold of 1e-13 and finding divergence, which are often sequence chimeras, these disagreements were resolved by cutting either the contig or the map, depending on the quality of the genome map at the divergent position. In total, sequence contigs at 487 loci were predicted to be incorrectly assembled and we resolved by cutting the contigs. A total of 1100 error resolved sequence contigs were scaffolded into an assembly with an N50 of 28.76 Mbp. Software used for this assembly and hybrid scaffold is available at www.bnxinstall.com/solve/Solve3.1_08232017.tar.gz and additional details description can be found at <https://bionanogenomics.com/support-page/bionano-solve/>.

#### X.X. Lifting of genetic marker positions from Bauer et al. contigs

The draft ‘Lo7’ genome assembly of Bauer et al. (2017)^8^ was co-opted as a source of mapped genetic markers. Contigs of the Bauer et al. assembly (n=1,581,707) had themselves been anchored to 87,820-markers of a high-density map constructed using the Rye600k SNP genotyping array, by genotyping a near-homozygous recombinant inbred line (RIL) population with parents ‘Lo7’ (seed) and ‘Lo225’ (pollen)^8^. These contigs were mapped to the raw scaffolds of the new assembly using minimap2^20^ with preset parameters for genome-to-genome mapping. Unique, primary, non-supplementary alignments with mapping qualities exceeding 30 were retained (n=1,535,590). The genetic positions associated with 44,371 such contigs were lifted to the new assembly scaffolds, with each genetic position being linked to the leftmost mapping position of the contig.

#### Supplementary Figure X.X. S-HICDIST

Fragment length histograms for Hi-C libraries, shown for both valid links and paired-end read contamination. Hi-C libraries for all taxa were constructed using the *Dpn*II enzyme, except the ‘Lo7’ library used in assembly, which was constructed with *Hind*III (as indicated in the facet label).

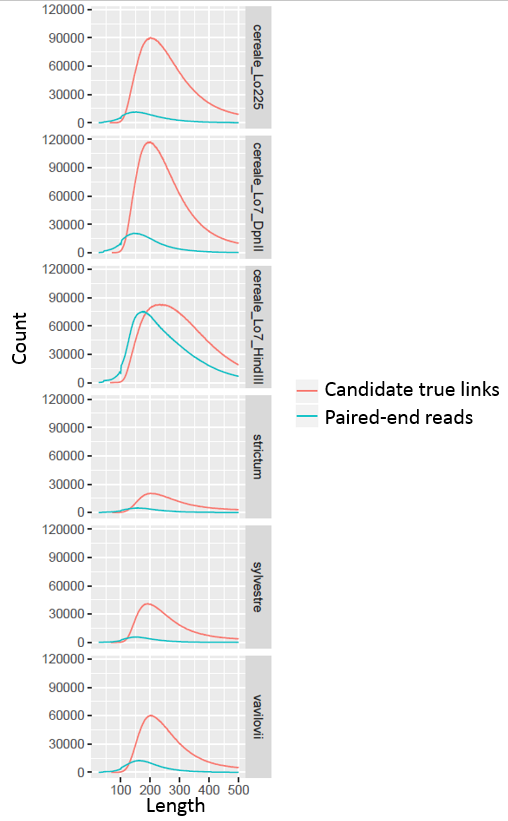

#### Supplementary Figure X.X. S-10XHIST1

Frequency histogram over the numbers of read pairs representing an inferred 10X molecule in 10X libraries mapped to the ‘Lo7’ assembly scaffolds. Read pair counts are binned into groups of 5.

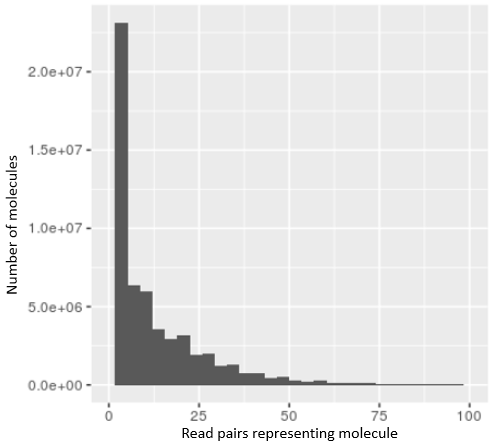

#### Supplementary Figure X.X. S-10XHIST2

Inferred frequencies of molecule lengths in 10X libraries mapped to the ‘Lo7’ assembly scaffolds. Molecule lengths binned into groups of 20,000.

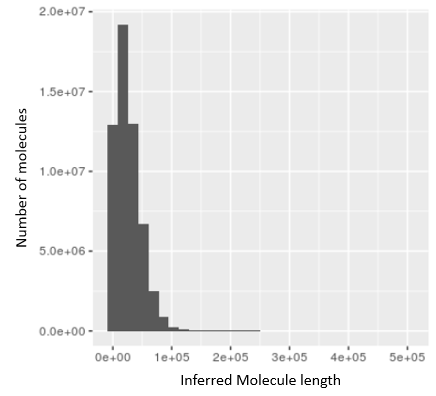

### Supplementary Note X. S-REP. Investigations into the repetitive genome

Extensive investigations were made of the repetitive space of the rye genome (using both element annotation and kmer-based approaches), with the goals of interrogating the completeness of the assembly (fig. S-RPT_ASSCMP), the effect of repetitive sequence upon this completeness as well as on the arrangement of scaffolds into pseudomolecules (fig. S-SATS), the efficacy of various tandem repeats as cytogenetic tools when used as probes for FISH, the evolution of the genome size and structure (fig. S-FISH), and the evolutionary history of transposable elements (figs. S-TEMAINPROG, S-TETERMPROF). This section presents the results of these investigations to support the arguments and comments made in the main text. The annotation of these elements is described in the online methods.

#### Supplementary Figure X. S-RPT_ASSCMP

*Poaceae* genome assemblies can be compared using repetitive DNA as a proxy for completeness**. a)** Mathematically defined overall repetitivity in the form of 20mer frequencies. 20mers occurring 10 times or higher covered only 16% of the draft rye genome assembly released in 2017^8^, whereas in the current assembly they account for 66% which is near the target value of 71% determined from 1X genome coverage of randomly sampled Illumina reads. The gap seen around a frequency of 100,000 (arrow) relates to the highly repetitive tandem repeats, which are far better represented in the present assembly. The arrow denotes the large general increase (mostly TEs), tandem repeats are around 10e5, and are still depleted even in the new assembly. **b)** Owing to their correlation with genome size, the number of retrievable full length LTR-retrotransposons can serve as a metric for the assembly quality of difficult repetitive regions^24^. The graph shows the number of retrieved fl-LTR candidates in different genome assemblies. Triangles represent earlier contig assemblies, which rarely correctly reconstructed the (almost) identical 1—2 kb long terminal repeats of fl-LTRs. The arrow shows the improvement in assembly improvement between the current rye assembly and the draft assembly published in 2017^8^. Circles denote more complete assemblies. Sb: *Sorghum bicolor*^25^; Zm: *Zea mays*^25^; Hv: *Hordeum vulgare*^26,27^; Sc: *Secale cereale*^8,28^; Td: wild emmer wheat^29^;
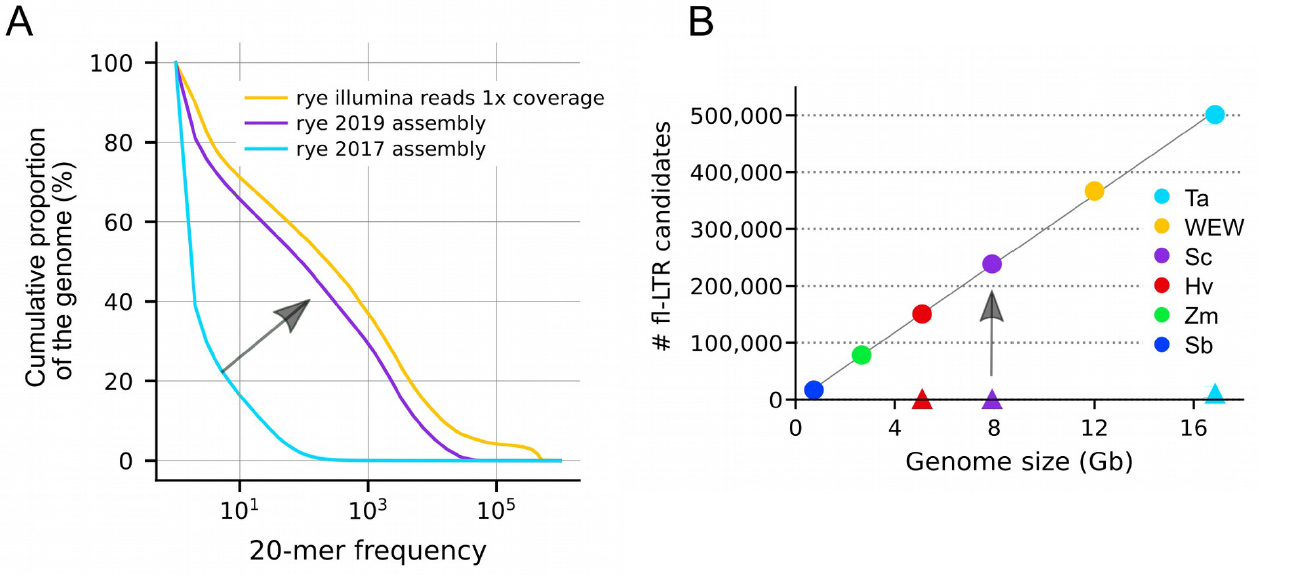

#### Supplementary Figure X. S-SATS

Localisation of tandem repeats in the rye assembly. **a)** Overall frequency of tandem repeats broken down into three main categories: (i) satellites with monomer units >= 100 bp (representing 44.6 % of all tandem repeats), (ii) minisatellites with 10—99 bp units (52%) and (iii) microsatellites with 2—9 bp units (3.4%). **b)** Chromosomal distribution of the three different tandem repeat types. Satellites are prominently located at several chromosome ends: short arms of chr 1R to 6R, most on 5RS and long arms of 2R and 3R. Most of the satellite sequences could not be assigned to a position on one of the seven chromosomes, and these where merged into chrUn. Minisatellites have two notable hot-spots on chr5L and are found in higher concentrations in the centromere. These can be used to identify the centromere in the assembly. Microsatellites are distributed more or less across all chromosomal regions.

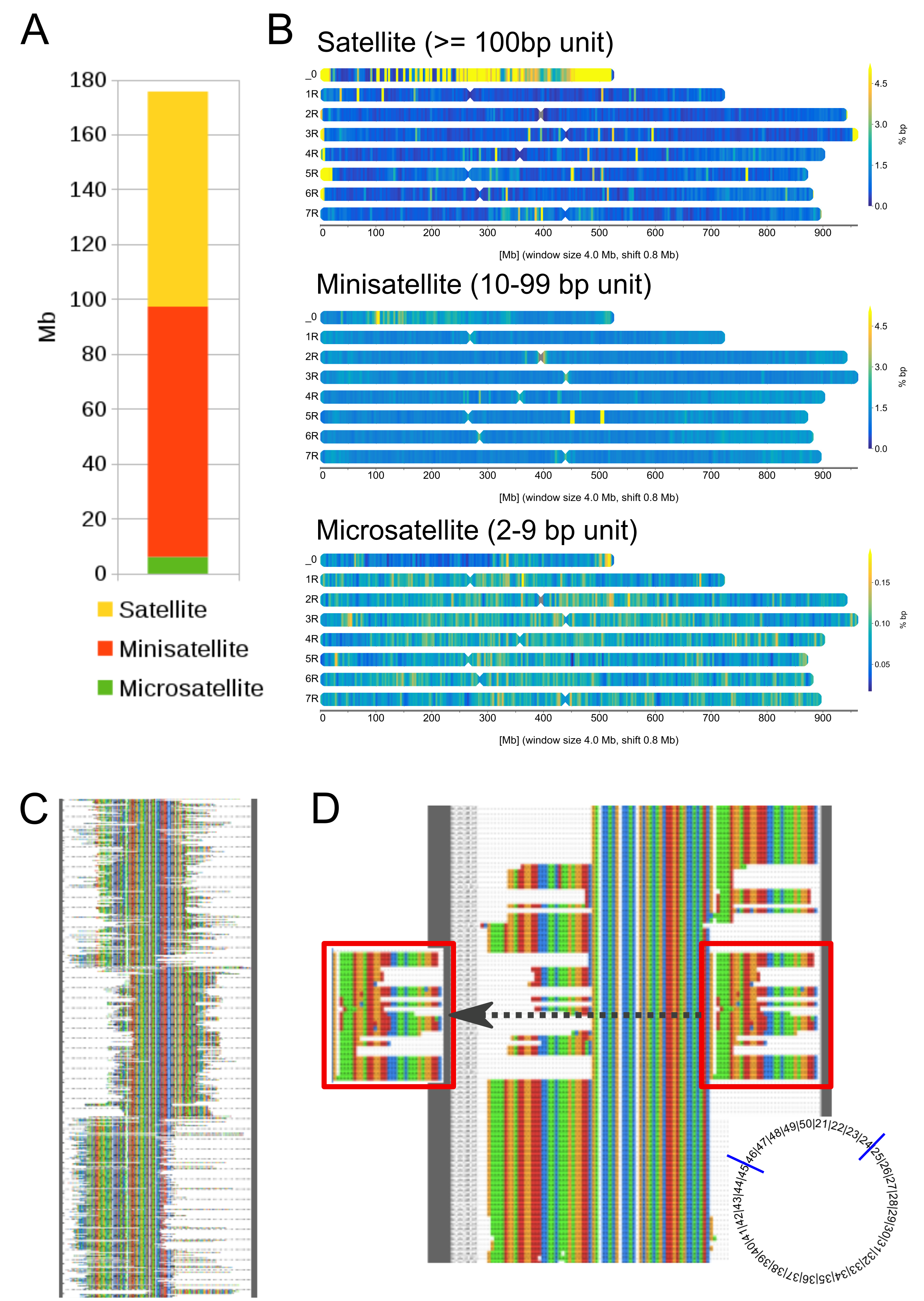

#### Supplementary Figure X. S-SATSFISH

*In silico* prediction and fluorescent in situ hybridisation (FISH) profiles for eight *de novo* identified tandem repeat families (tbl. S-FISH). Each subfigure shows the chromosomal distribution of tandem repeat sequence clusters in the assembly (left, bar charts) and their locations on the seven rye chromosomes as revealed by FISH (right, photograph and ideogram). The probes designed to Sat20/517 bound non-specifically under the tested hybridisation conditions (methods). The two satellite tandem repeat families from which the probes ScSat213 and ScSat380 appear to bind telomeric repeats which, notably, are displaced or not detectable from the subtelomeres wherever chromosomal rearrangements in the rye lineage have affected the structure of those regions (chromosomes 4R—7R). However, the predicted position of ScSat213 is in interstitial is in scattered interstitial clusters, while the only two significant clusters of ScSat380 are predicted at subtelomeres. Both these observations as evidence that subtelomeric clusters tend to be collapsed or unassembled, since ScSat380, while present on all fourteen subtelomeres only appears in the assembly at two, and ScSat213 at none. A possible reason that comparatively minor assembled interstitial clusters are overemphasised in the bar chart. The remaining five of eight FISH probes bound as predicted by the annotation (see also table S-FISH). Among them is ScSat44 which represents a new 5R specific probe. The bar in the upper right image represents approximately 10 µm and the boxed chromosomes have been digitally moved to fit into the frame.

(Continued over)

(Cont.) Supplementary Figure X. S-SATSFISH
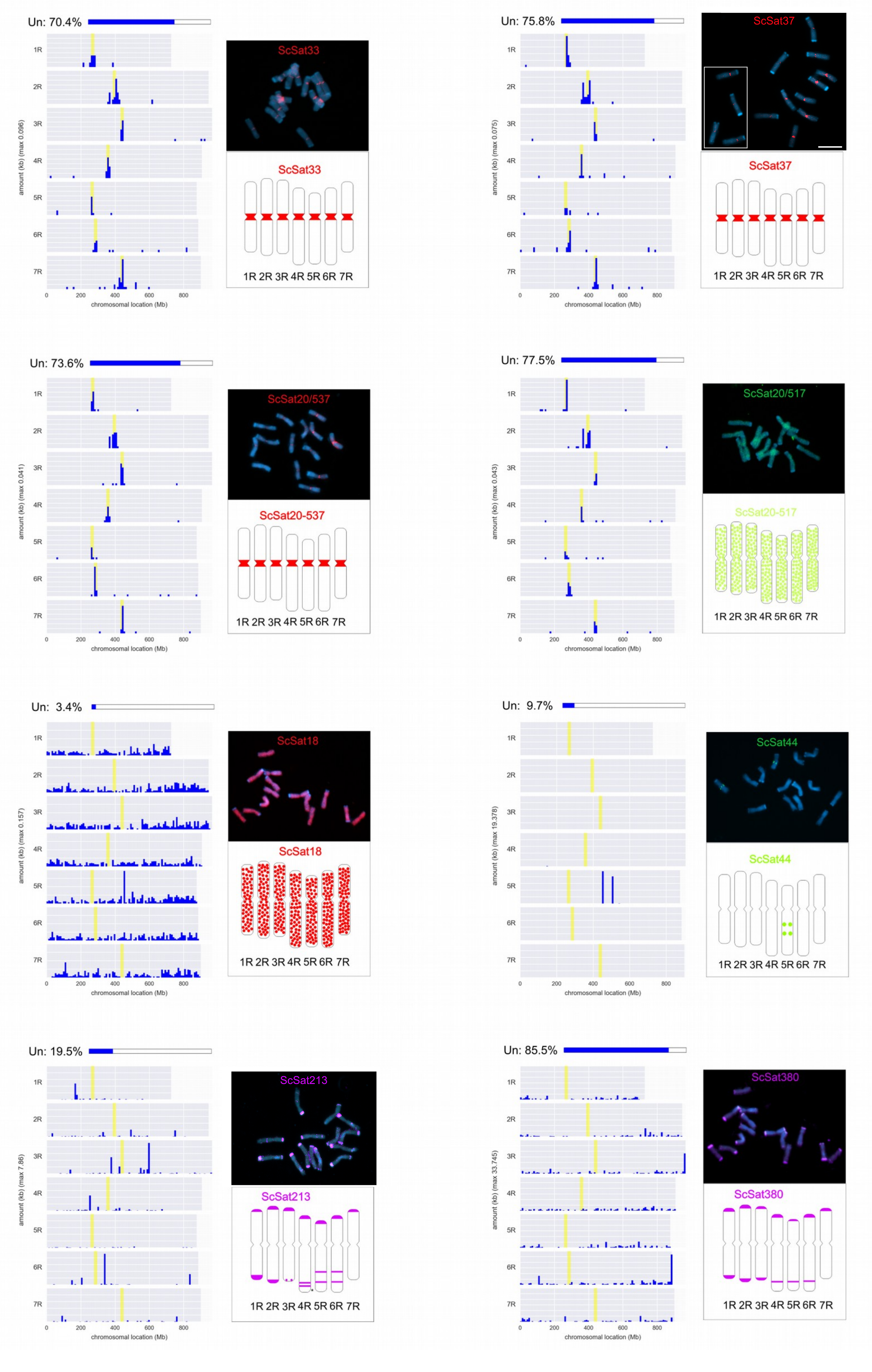

#### Supplementary Figure X.X. S-KMERREP

Kmer repetitiveness compared between rye and barley, chromosomes 1—7 shown left-to-right. The overall increase factor in assembly size from barley^27^ to rye is about 1.5 (4,507 vs 6,735 Mbp), which is suspected to be accounted for largely in repetitive sequence, and the bars below the chromosome numbers depict the size increase factor for the short and long chromosome arms respectively, which show (except for chromosome 6) no large deviations from the overall 1.5 increase. The two line charts track changes in repetitivity along the chromosomes (median 20mer frequencies over 4 Mbp sliding windows with a 0.8 Mb shift) for barley (A) and rye (B). Rye displays a similar division into chromosomal compartments of different repetitivity grades as previously described for barley^27^. The suspected introgressions that affected the transposon profiles at the terminal long arms chromosomes of 4R and 6R (main text; fig. M-TRACKSd—e) may also have caused local reductions in the 20mer repetitive content.

Tracks C and D compare rye-barley syntenic blocks (colour-coded to correspond to the colours assigned to barley chromosomes in main figure M-TRACKS) with ideograms for FISH probes ScSat213 and ScSat380, to emphasise the correspondence between terminal rearrangements and the displacement of telomeric FISH hybridisation signals (see fig S-SATSFISH).

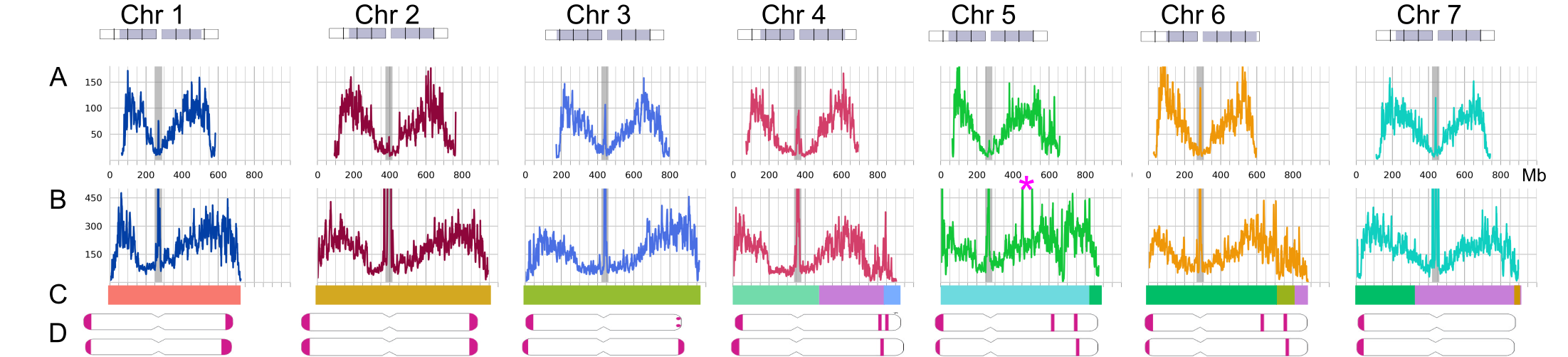

#### Supplementary Figure X.X. S-TEMAINPROF

Distributions of annotated TE families along the ‘Lo7’ pseudomolecules. **a)** Distribution of main TE families along rye chromosomes. Shown are the number of full length copies in bins of 40 Mbp. Note that the short (left on the image) ends of 4R and 6R have a different TE composition than the rest of the genome. **b)** Contribution (in %) of diagnostic TE families to the terminal 150 Mbp of chromosomes 1R, 2R and 4R. Some TE families are strongly enriched on 4R while others are depleted or virtually absent. **c)** Examples of distributions of TE families along chromosome 4R. Colours correspond to different TE subfamilies, which are dissected in greater depth in figs. S-ANGELA—S-CEREBA.

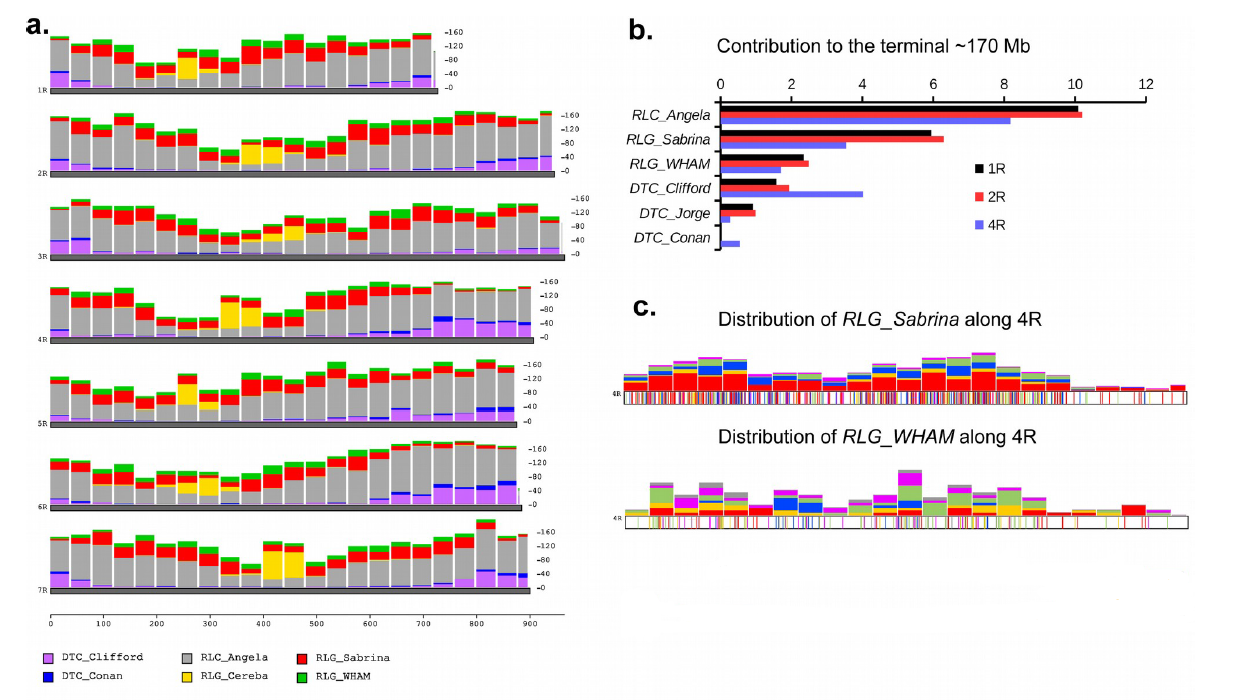

#### Supplementary Figure X.X. S-TETERMPROF

Relative abundance of four major TE families in the terminal 175 Mbp of all rye chromosomes. Here, we counted all full-length copies of the four TE families in the terminal 170 Mbp of the long chromosome arms. Note that chromosome 4R has a composition distinct from all others with much higher numbers of DTC_Clifford elements while RLG_Sabrina and RLG_WHAM are practically absent, which we interpret as evidence of possible ancient translocation events.

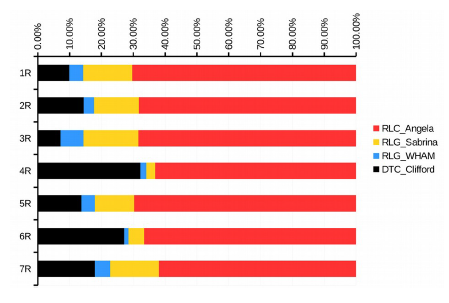

#### Supplementary Figure X.X. S-SUBRATE

We assert in the main text that substitution rate variation across the genome are adequate to explain findings previously interpreted as signals of reticulate evolution without requiring on intraspecific translocations^28^. Rates of synonymous substitutions per synonymous site in fourfold degenerate codon sites in coding regions of genes, between ‘Lo7’ and the D genome of bread wheat^30^. Substitution rates were determined by alignment of bi-directional closest homologs from rye and the wheat D genome (Methods). The x-axis gives the chromosomal position in Mbp, while the y-axis shows the number of synonymous substitutions per synonymous site calculated in a running average over 100 genes. The blue line shows substitution rates in all genes, while the red line shows substitution rates in single-copy genes only. The observed rates grade gradually, rather than changing blockwise, across the genome, and the rates between which they grade (between ~ .05 and .14) are consistent with those previously estimated^28^.

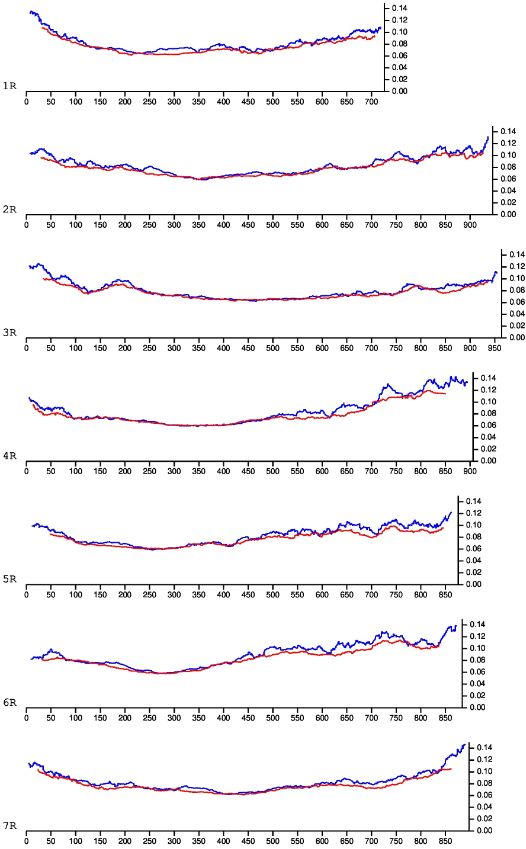

#### Supplementary Figure X.X. S-RECOMB

Recombination rate can be a driver of substitution rates. We therefore estimate here the recombination rate variation across the genome, inferred from the genetic map coordinates lifted from the ‘Lo7’ x ‘Lo225’ genetic map reported in Bauer et al. (2017). Positions on the x-axis give the mean physical positions of groups of 100 consecutively-placed markers (taken in steps of 50 markers). The y-axis (cM per bp) was calculated using the R function ‘lm’ as the slope of a least-squares regression line ( cM ~ bp ) over each 100 marker group.

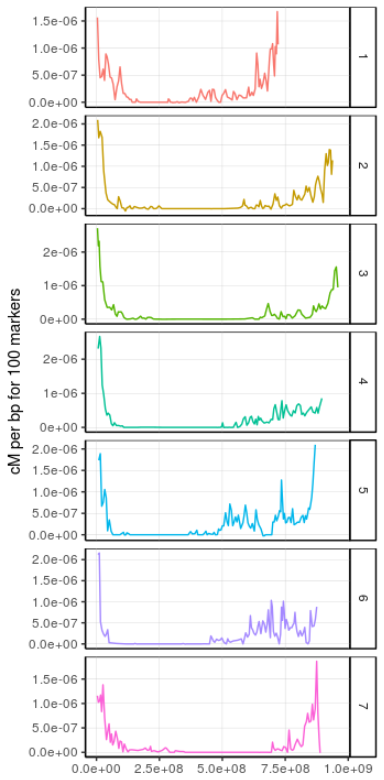

#### Supplementary Figure X.X. S-TETRANSLOC

We diagram here a model for the evolution of chromosome 4R. Chromosome 4R is a composite chromosome that consists of segments of ancestral Triticeae chromosomes (as inferred by rye-wheat-barley collinearity^28^) 4, 6 and 7, that underwent rearrangement in the rye lineage since its split from wheat and barley. The segments are indicated and the orientation of the ancestral chromosome segments is indicated with arrows. The rye ancestor diverged into different lineages. In the ‘Lo7’ lineage, RLG_Sabrina and RLG_WHAM retrotransposons were very active, while in a second, hypothetical, lineage, the CACTA family DTC_Clifford was highly active. At least 1.8 million years ago, the two lineages recombined, leading to the introgression of a terminal ~170 Mbp segment into the ‘Lo7’ lineage. Subsequently, the genome was invaded by RLC_Angela retrotransposons.

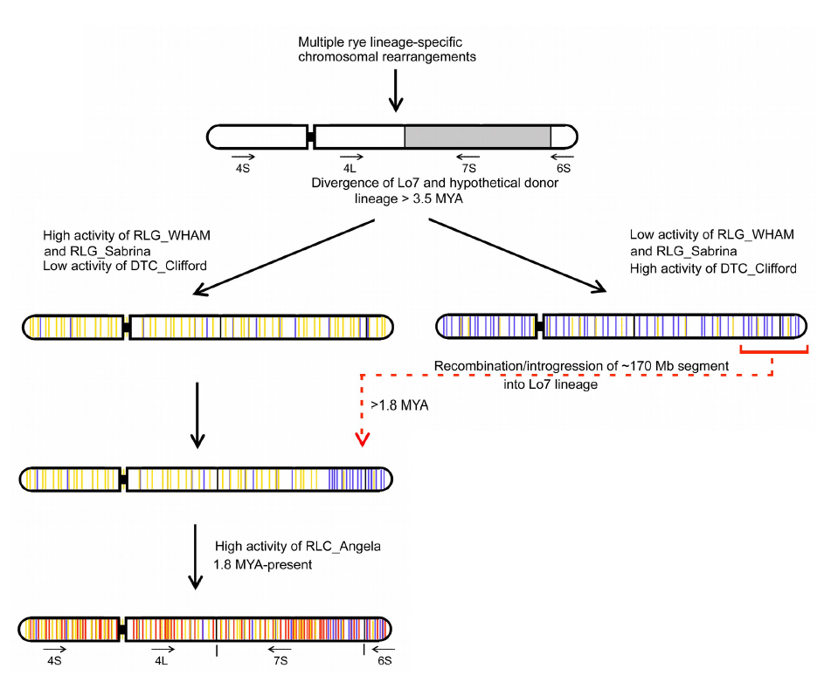

### Supplementary Note X. S-TEEXP: Arguments on the timing of TE superfamily expansions in various Triticeae

As mentioned in the main text analysis of retrotransposon, evolution using recently assembled reference quality Triticeae genomes show that in rye, barley and each individual wheat subgenome, the TE superfamilies RLG and RLC appear to have experienced their major periods of dominance in the same order^31,32^. We attribute this to parallel evolution and expand upon our reasoning here.

This common order could constitute evidence for A) parallel evolution, whereby some common (e.g. environmental) trigger promoted expansion of different families at different times; B) parallel evolution driven primarily by expansion properties inherent to the TE families themselves, but moderated by the genomic context; or C) frequent horizontal exchange of TE-containing germplasm followed by expansion of the newly acquired element. The common trigger hypothesis (A) conflicts with the observation that the expansions of each family happened at different times, even differing between wheat subgenomes^31^. Horizontal exchange (B) does not explain the common expansion order per se; the question merely regresses to asking why the order in which these horizontal exchanges donated TEs of a particular family was the same for each species (and indeed, why a particular germplasm exchange should transfer only one family of TE). We also show below that there is no strong evidence for intraspecific germplasm transfer from non-rye Triticeae into the rye lineage. Thus, we argue that parallel evolution directed by properties of the common ancestral suite of TEs (C) is most likely. Under this model, it would make sense that the Gypsy superfamily was a more aggressive expander in the context of the A-B-D-H-R ancestral genome, but was in every instance outcompeted by members of the Copia superfamily later. This would predictably have occurred once RLG-suppressive elements evolved in each genome, independently, since there will always come a point at which unchecked expansion of a TE family is detrimental. Dormant elements of the Copia superfamily, inherited from the common ancestor, happened to be best suited to a fertile genomic niche populated with suppressed Gypsy elements. Ergo, this Gypsy-to-Copia progression was set about by events in the shared ancestral genome, but the rates of expansion and suppression would have depended upon functional and selective peculiarities of each (sub)genome, thus explaining the common order but different expansion times.

### Supplementary Note X. S-SV: *Secale* diversity and segregating structural variations

We expand here upon the discovery of structural variants (SVs) using Hi-C as presented in the main text, the statistical test that was run to affirm association between SVs and the peri-centromeric low-collinearity modules.

#### Supplementary Figure X.X. S-HICSV

Hi-C asymmetry plots revealing Structural Variations (SVs)—Inversions in particular—between the genomes of ‘Lo7’ and representatives of the genus *Secale*. The four relatives studied are related to the reference line ‘Lo7’ as follows: [ [ *S. cereale* line ‘Lo7’, *S. cereale* line ‘Lo225’ ], *S. vavilovii* ], [ *S. strictum*, *S. sylvestre* ]. SVs result in result in discontinuities in the ratio of Hi-C links mapping left:right (r) relative to ‘Lo7’ (r_sc_). Inversions often produce clean, diagonal lines. Indels, translocations, and deletions all cause interruptions, but identification of all SV types can be confounded by sequence errors, contiguous/overlapping SVs, low mapping rates when aligning reads from divergent genomes, and other factors. Visually-identified candidate SVs are shaded, with common non-grey colours indicating possible synapomorphic (shared, derived) SVs. Shading is omitted from some anomalies at centromeres where missing sequence causes artefacts. Blue lines denote the boundaries of the low collinearity regions (LCMs) for each chromosome. The rightmost inversion marked on 5R corresponds to the region of recombination suppression marked on figure 1A. The observed SVs are consistent with the inferred phylogenetic relationships and limited horizontal genetic exchange since all putatively shared SVs appear to be synapomorphic. This is expected because large SVs not only reflect, but also likely reinforce, reproductive barriers between groups.

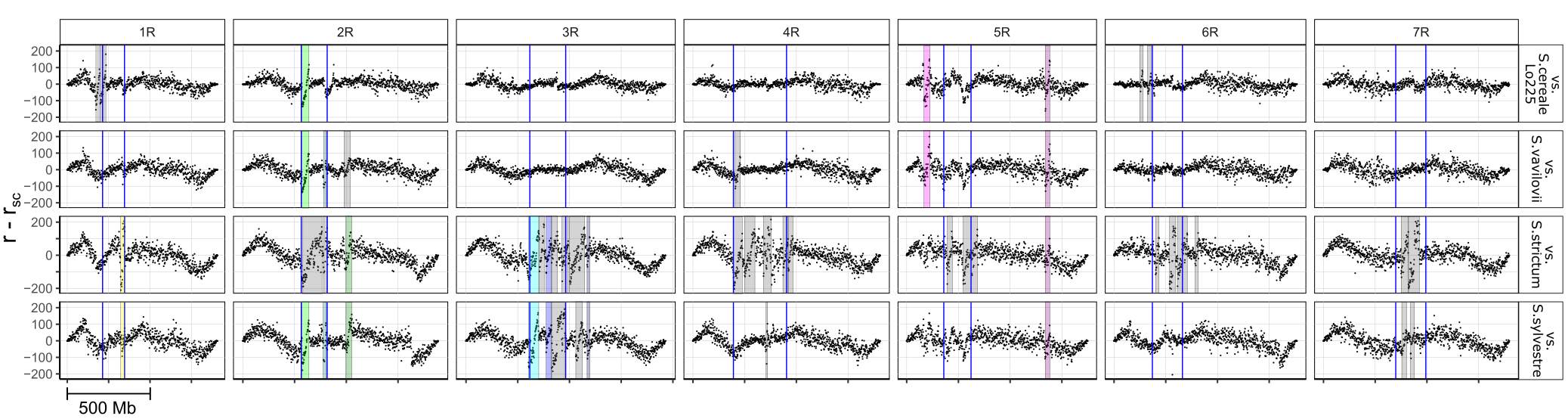

#### Supplementary Figure X.X. S-HICPERM

We used a permutation method to confirm that these large SVs do show an affiliation for LCMs, by randomly reshuffling the positions of observed SVs on the chromosomes. Overlaps were not allowed, and synapomorphic SVs were placed at the same positions on multiple genomes. The count axis (figure below) records the number of times SVs overlapping the LCM under random placement (10000 iterations). The black vertical line shows the number of LCMs overlapping in the dataset (1). Testing the hypothesis that the data are produced by randomly-placed SVs results in a p-value of < 1/10000, demonstrating that SVs almost certainly tend to occur in association with these areas of disrupted collinearity.

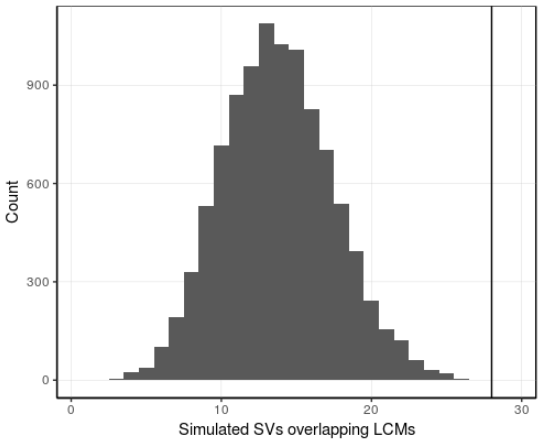

### Supplementary Note X. S-COLIN

We used a best-reciprocal-matches approach with transcriptome data to create high resolution collinearity maps between rye and its Triticeae relatives barley^27^ (*Hordeum vulgare* cv. Morex), and the three subgenomes of bread wheat^30^ (*Triticum aestivum* cv. Chinese Spring; Methods). Figures S-COLIN_lg1—S-COLIN_lg7 break down the results to show collinearity between every chromosome of every genome. As noted in the main text, the curvature of collinear regions indicates changes in the rate of genome expansion/contraction since genetic isolation between two species. The colour key, which applies across the whole note, relates to the homologous group of the chromosome on the horizontal faceted axis.

#### Supplementary Figure X.X. S-COLIN_lg1

Chromosome-wise best-reciprocal-matches showing Triticeae collinearity for chromosome homologous group 1.

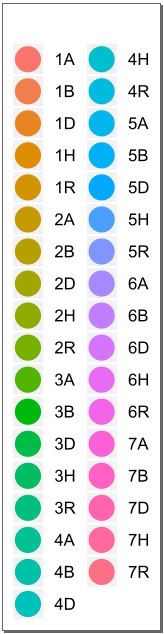

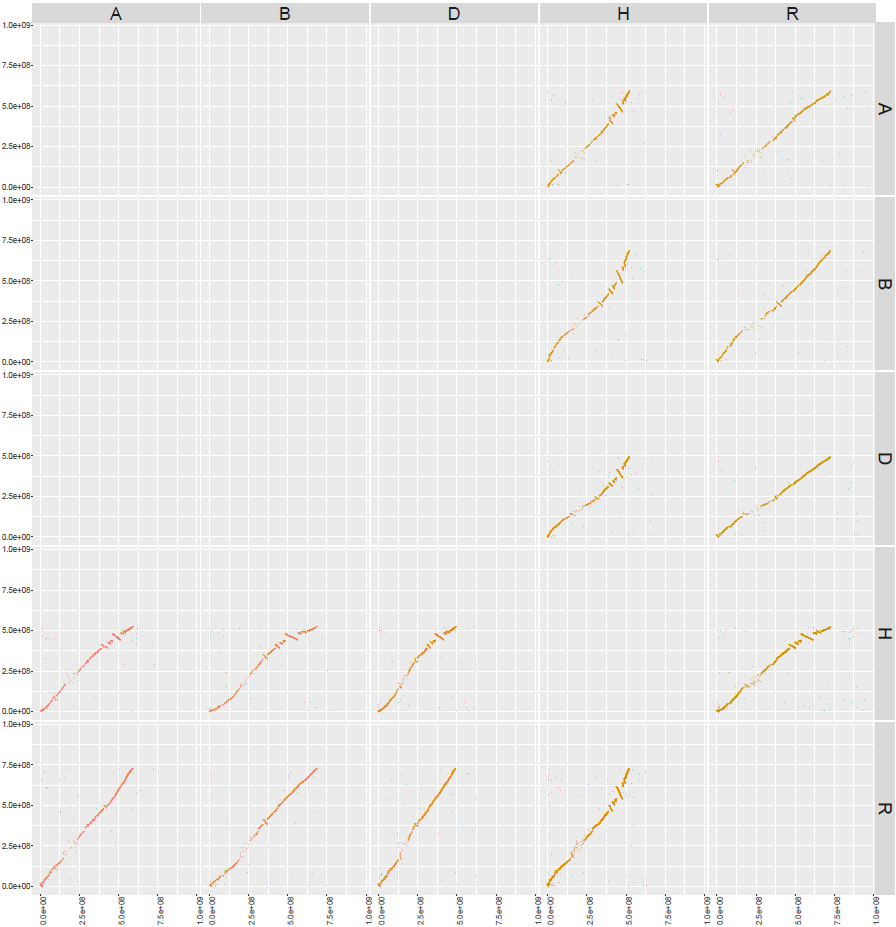

#### Supplementary Figure X.X. S-COLIN_lg2

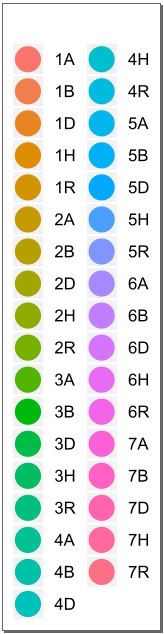
Chromosome-wise best-reciprocal-matches showing Triticeae collinearity for chromosome homologous group 2.
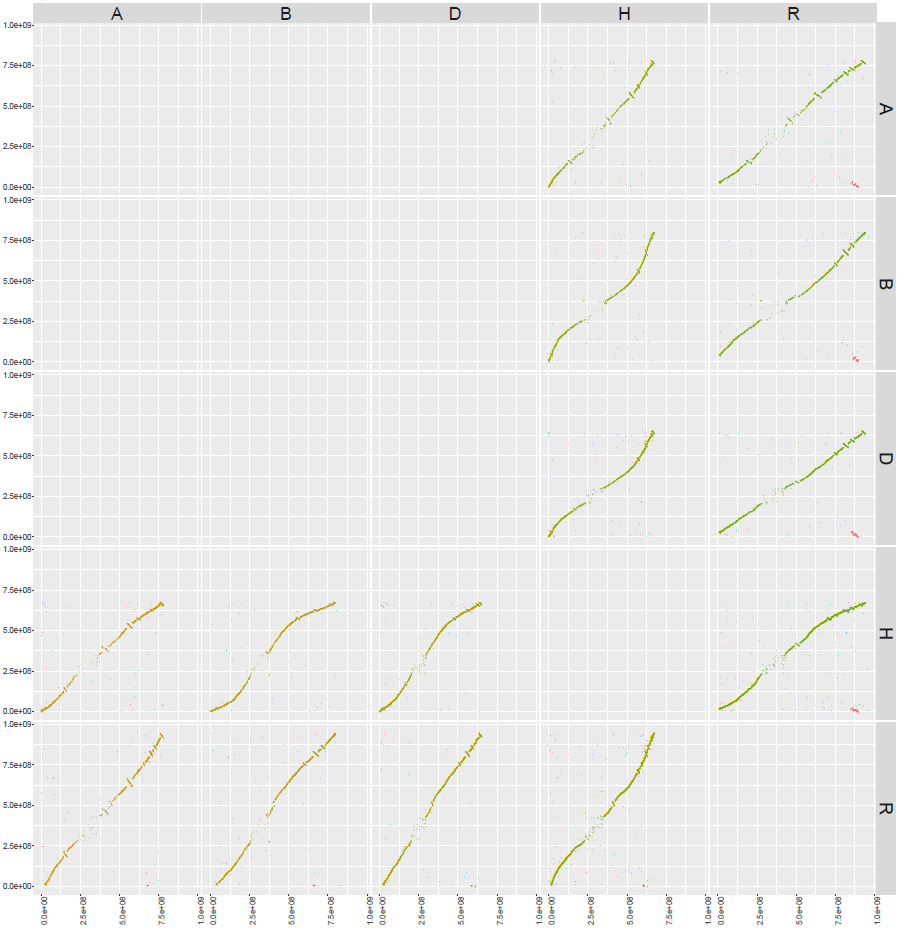

#### Supplementary Figure X.X. S-COLIN_lg3

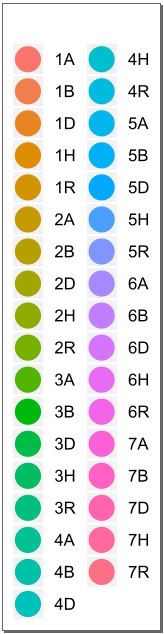
Chromosome-wise best-reciprocal-matches showing Triticeae collinearity for chromosome homologous group 3.
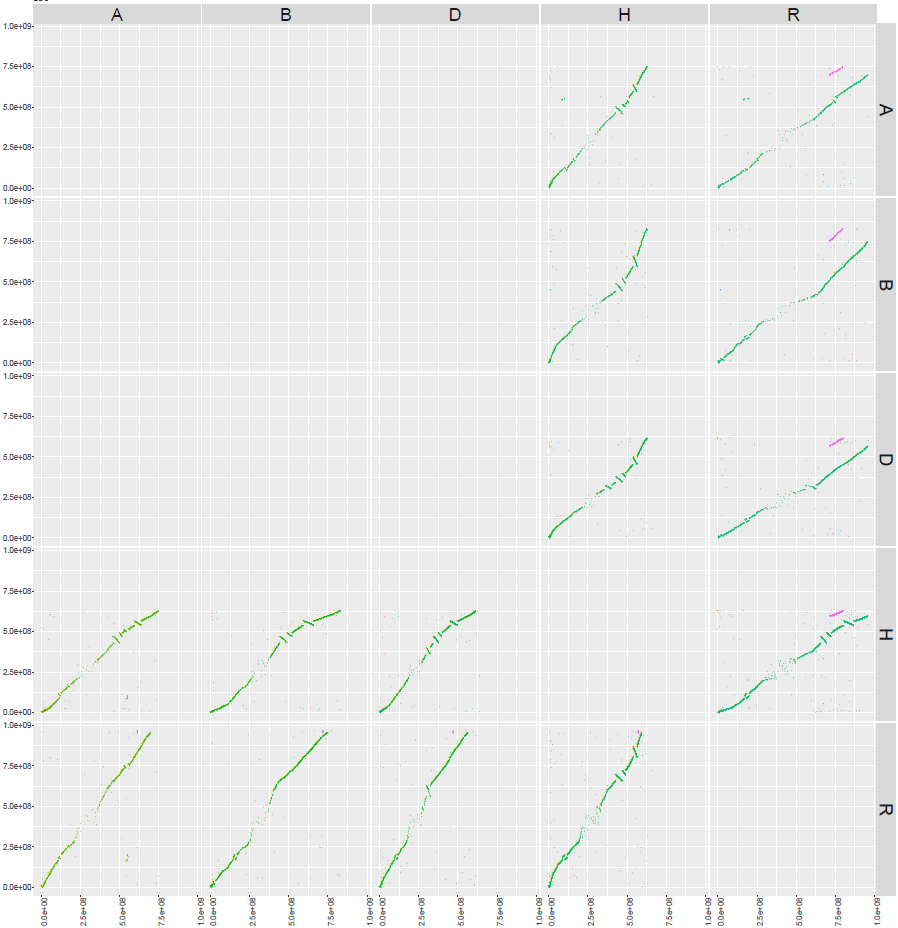

#### Supplementary Figure X.X. S-COLIN_lg4

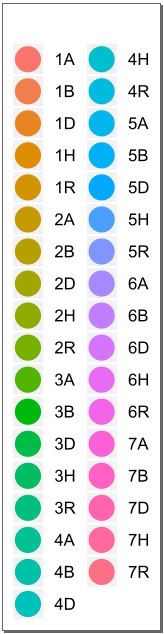
Chromosome-wise best-reciprocal-matches showing Triticeae collinearity for chromosome homologous group 4.
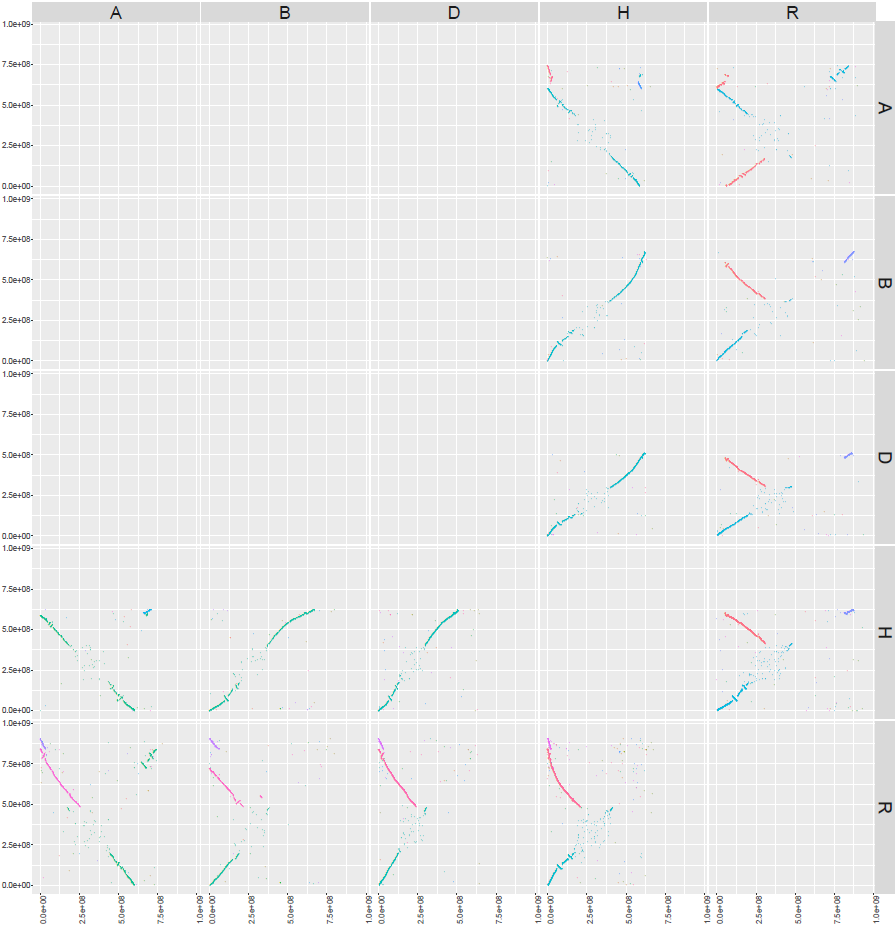

#### Supplementary Figure X.X. S-COLIN_lg5

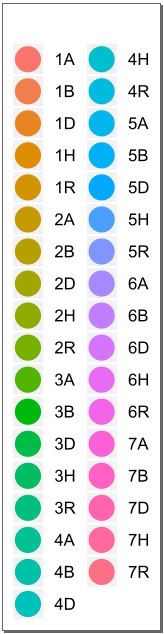
Chromosome-wise best-reciprocal-matches showing Triticeae collinearity for chromosome homologous group 5.
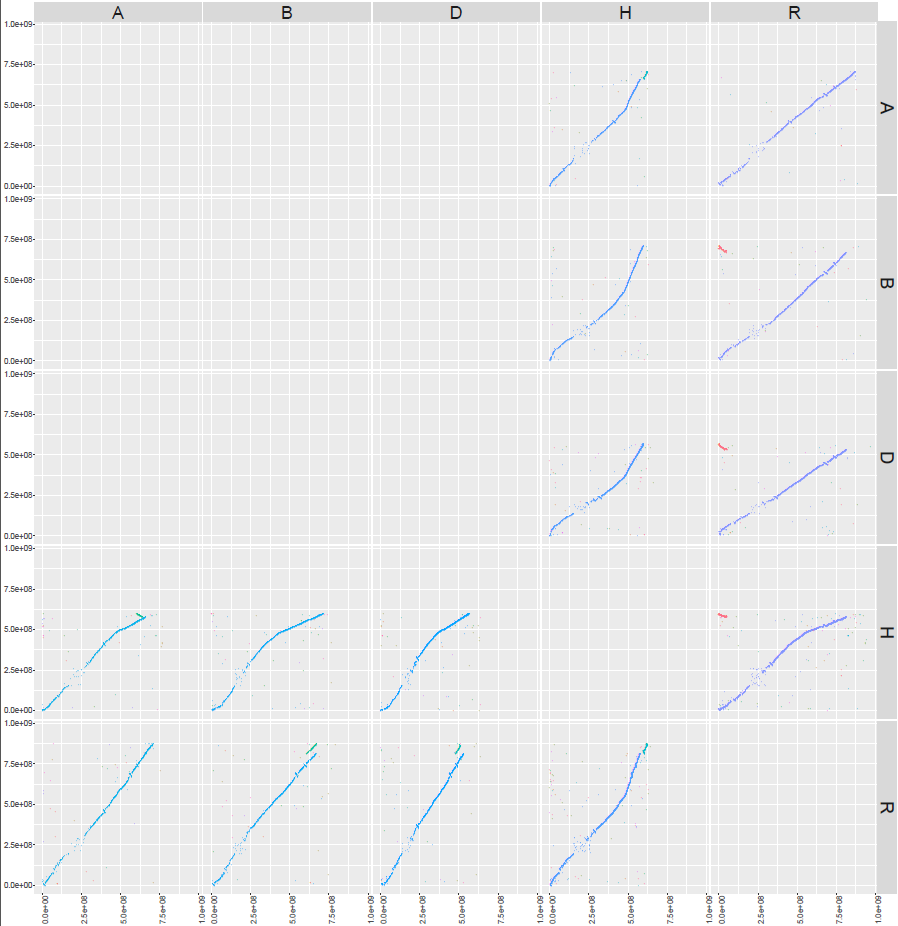

#### Supplementary Figure X.X. S-COLIN_lg6

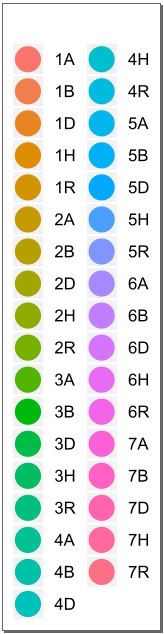
Chromosome-wise best-reciprocal-matches showing Triticeae collinearity for chromosome homologous group 6.
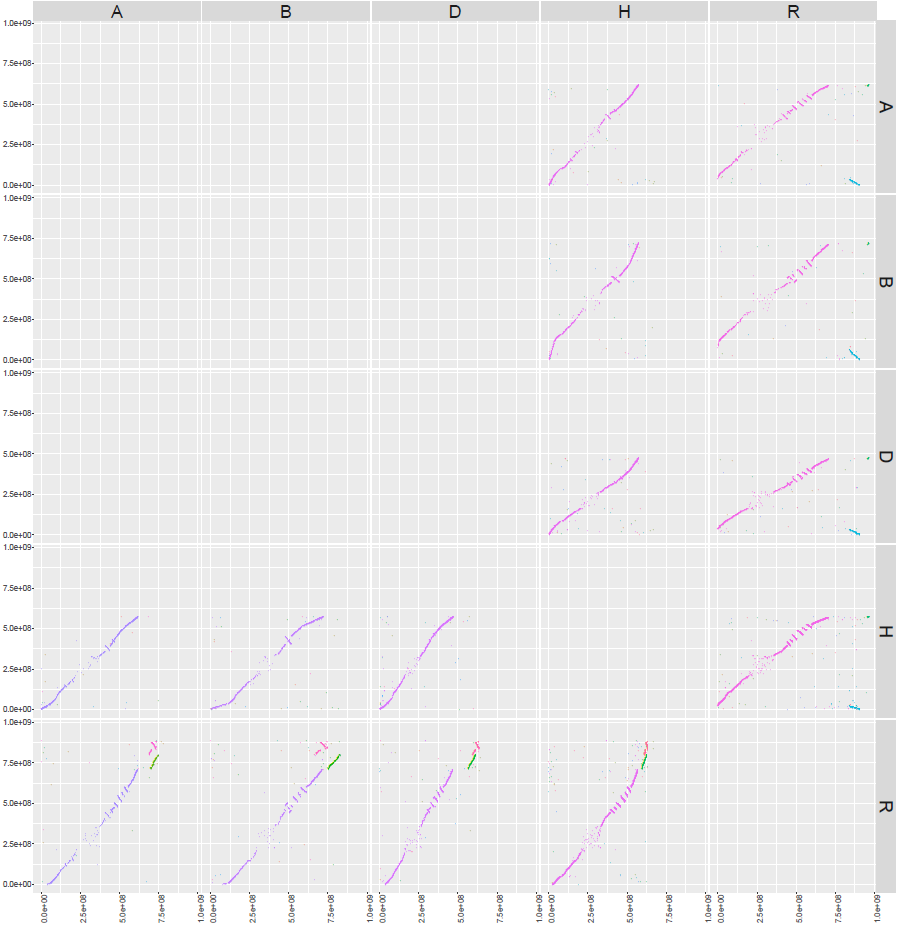

#### Supplementary Figure X.X. S-COLIN_lg7

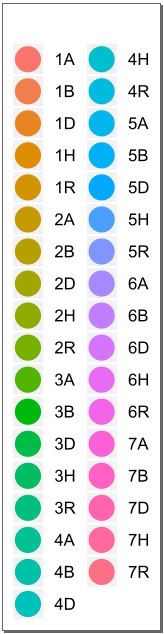
Chromosome-wise best-reciprocal-matches showing Triticeae collinearity for chromosome homologous group 7.
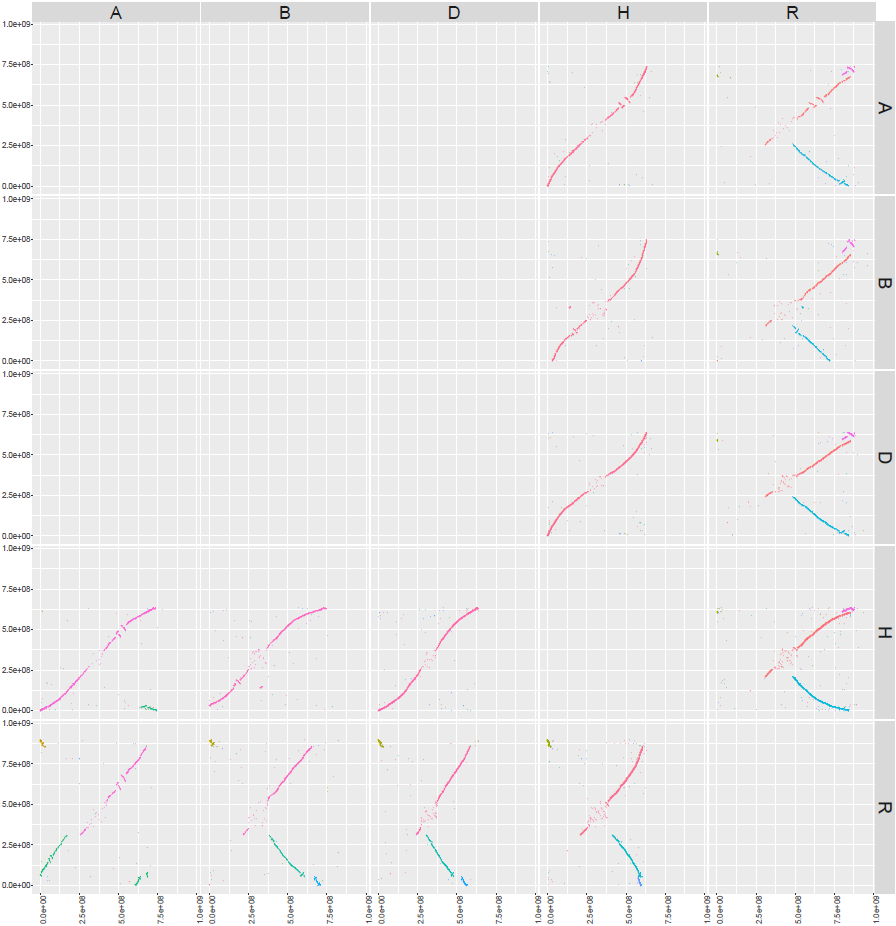

### Supplementary Note X. S-OUTIN

The mTERF and RFL gene families are expected to increase in size in outcrossing taxa, as predicted by evolutionary theory^33^. We conducted a preliminary analysis on this question here to exploit the availability of new Triticeae genomes that can assist in testing this hypothesis.

#### Supplementary Figure X.X.

Comparative counts of annotated mTERF (y-axis) and RFL (x-axis) genes in inbreeding and outcrossing species with high quality annotated genomes. The information sources are given in table S-PPR_BREEDINGSYS. In the case of sub-genomes in polyploid species, the reproductive strategy of their direct ancestors was used in their classification as inbreeding or outcrossing.

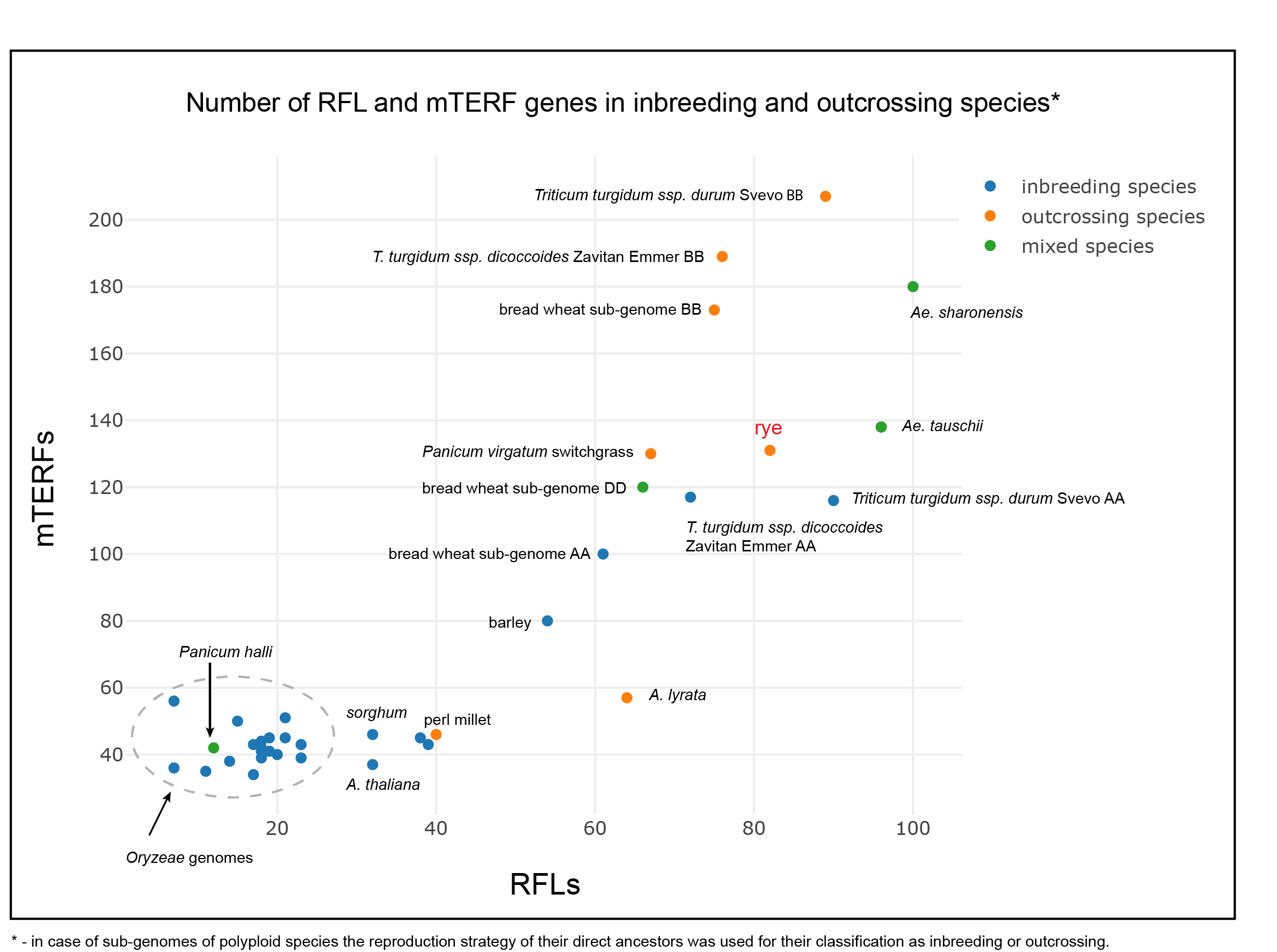

### Supplementary Note X: S-RFMULTI

Main text figure M-GENES compares the structures of the Rf^multi^ locus in rye and wheat, and we expand it here to include more detail including transcript identifiers.

#### Supplementary Figure X.X. S-RFMULTI

Organisation of RFL genes on 1R (A) at the ‘Lo7’ Rf^multi^ locus (B) compared to its wheat (Chinese Spring) counterpart on 1B (C). Flanking markers are shown on either end of the rye sequence. Two full-length wheat RFLs and a putative rye ortholog are highlighted yellow. PPR genes are coloured red.

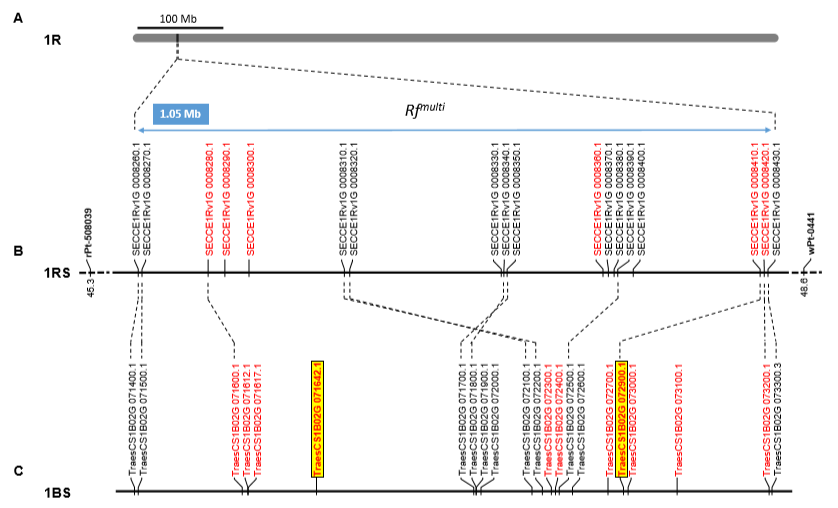

### Supplementary Note X. S-NLR: Comparative cluster arrangement and phylogenetics of resistance genes

Phylogenetic and homology search-based analyses (Methods) were applied to families of resistance genes (R genes), members of which are known to confer resistance to fungal pathogens in particular (figs. S-NLRPHYLO—S-LR10CMP). A similar approach was taken to identify possible low temperature tolerance (LTT) genes of interest in rye, based upon their homology to wheat LTT genes (fig. S-CBFPHYLO). Results underlying the main arguments and conclusions of the main text are expounded in detail in the following note.

#### X.X Phylogenetic and homology-based investigations of resistance gene orthologs

Nucleotide-binding leucine-rich repeats (NLR) are commonly associated with pest and pathogen resistance. We conducted a survey of NLR genes known to confer resistance to pathogens in various model systems and identified promising targets for future investigation based on their evolution within the Triticeae. For each NLR family of interest, we examined the phylogenetic relationships among members of the family within and between taxa, and used sequence homology to reconstruct structural changes in any notable loci containing these genes.

We identified homologs of *Pm2* and *Pm3*, two wheat powdery mildew resistance genes^34,35^, of *Mla*, an allelic series of more than 30 barley resistance genes^36^ and of wheat resistance genes *RGA2* and *Lr10^37^* in the three reference genomes.

The *Pm2* and *Pm3* genes are implicated in powdery mildew resistance in wheat^34,35^, and the recognition by wheat *Pm2* of the rye powdery mildew protein *AvrPm2* suggests an active *Pm2* allele would be beneficial to rye^38^. Phylogenetic analysis revealed two clades of *Pm2*, one of which includes a resistance-associated allele known from wheat (fig. S-PM2PHYLO). Resistant-clade *Pm2s* appear in a single copy on each group 5 wheat subgenome, but have been tandemly duplicated on rye 5R, and quadruplicated in barley 6H (main fig. M-GENESg—k; fig. S-PM2CMP). The rye duplication was followed by a series of TE insertions, one of which is probably implicated in the fragmentation of one of these *Pm2* copies (fig. S-PM2EVO).

*Pm3* cluster on the translocation-friendly 1RS and fall into three clades (fig. S-PM3PHYLO), the loss of one clade in rye and another in barley suggesting some inter-clade redundancy. Rye *Pm3* derives from the same clade as its orthologs on 1B, 1D, and 1H. Two further rye orthologs were identified as *Pm8* and *Pm17*, both known rye-to-wheat introgressions^39,40^, the latter evidently a chimera derived from unequal crossing over between *Pm8* and an unknown *Pm*-family gene not present in ‘Lo7’ (fig. S-PM3CMP).

*Mla* genes share close homology with wheat stem rust resistance genes *Sr33* and *Sr50*, both originally introgressions from rye 1RS, suggesting a gene family adapted to a potentially wide range of fungal interactions^41^. Three subclades can be identified (fig. S-MLAPHYLO), and like wheat chromosome 1A, ‘Lo7’ contains no members of subclade 2 which contains the known resistance allele for barley; Rather, the 1R cluster is likely composed of an expanded number of subclade 1 and 3 *Mla* genes (fig. S-MLACMP).

In wheat, the leaf rust resistance locus *Lr10* occurs in two distinct haplotypes, H1 and H2, distinguished by the presence and arrangements of *Rga2* family genes^37,42^. Analysis of TE insertions suggests that haplotype H2 derives from TE-driven rearrangements of an H1-like ancestral haplotype, and the conservation of H1 and H2 among diverse wheats suggest long-term stabilising selection affecting both alleles. We identified in rye an ancient derivative of the H2 allele (fig. S-LR10CMP), demonstrating the extreme length of time over which stabilisation has been occurring, and inviting further research into the reason for *Rga2-a*’s persistence across three species.

#### Supplementary Figure X.X. S-NLRPHYLO

Phylogenetics and genome-wide distribution of genes in rye, broken down by subclade. A) Multidimensional scaling of the 997 NLR genes annotated in rye based on pairwise distances calculated from phylogenetic tree shown in panel B. The colours separate three main groups of NLRs. The black dots indicate homologs of *Pm2*, *Pm3*, *Mla* and *RGA2*. B) Phylogenetic relationships between 792 manually-annotated NLR genes (tbl. S-NLR). Colours correspond to the subclades in A and homologs of *Pm2*, *Pm3*, *Mla* and *RGA2* are marked.

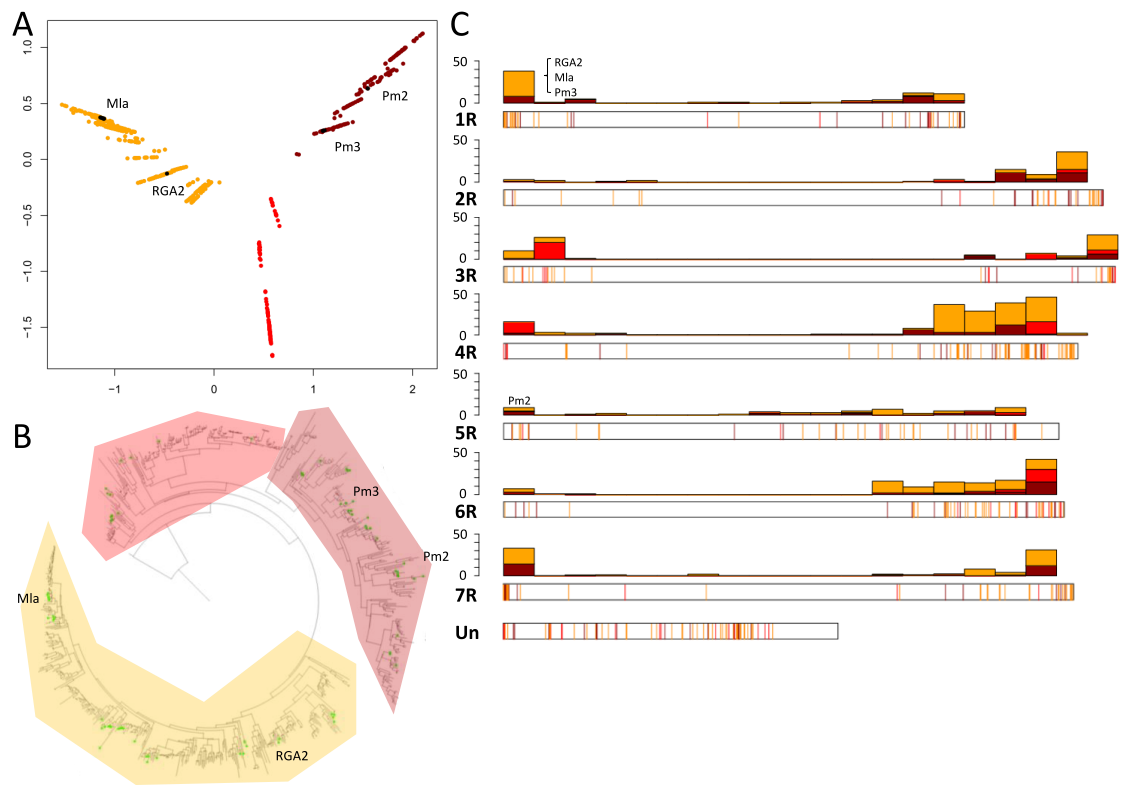

#### Supplementary Figure X.X. S-PM2PHYLO

Phylogenetic relationships among full-length *Pm2* homologs in rye, wheat and barley. The two main clades are shown in red and blue. Bootstrap support values are given at nodes.

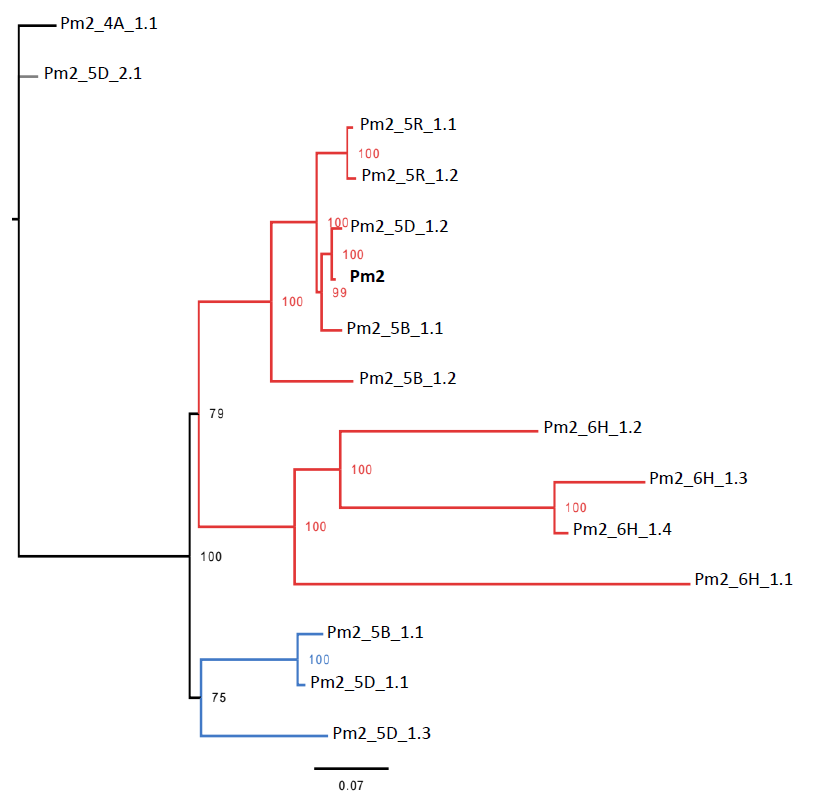

#### Supplementary Figure X.X. S-PM2COMP

Physical organisation of *Pm2* cluster haplotypes in the (sub)genomes of cultivated Triticeae. Identifiers refer to S-PM2PHYLO with colours separating phylogenetic clades. Pseudogenes (i.e. those which contain in-frame stop codons and/or frameshifts) are marked with a psi symbol (Ѱ).

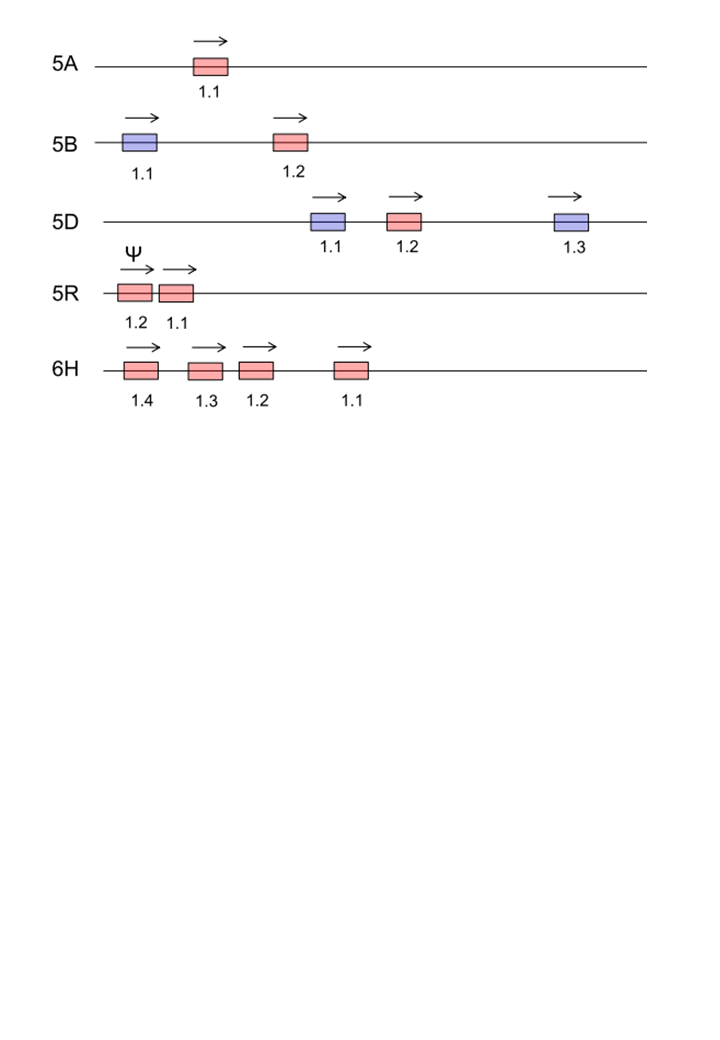

#### Supplementary Figure X.X. S-PM2EVO

Suggested mechanism leading to the duplication and pseudogenisation of the *Pm2* locus in ‘Lo7’. The two *Pm2* homologs are shown in light red, transposable elements in orange, yellow, green and purple. The two horizontal arrows indicate the duplication.

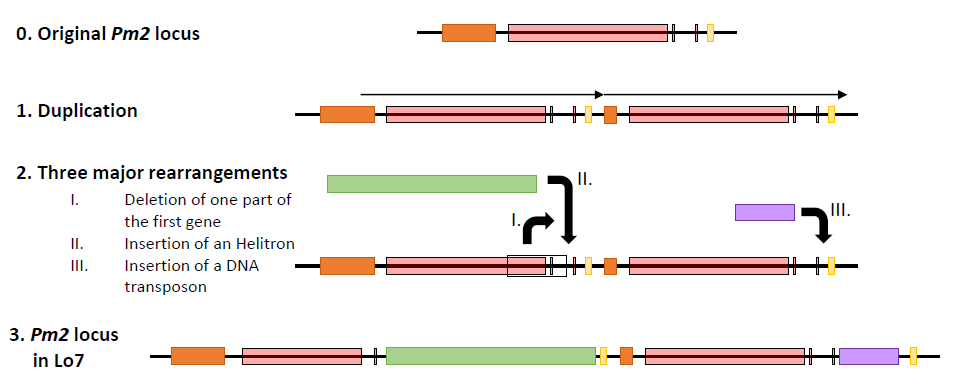

#### Supplementary Figure X.X. S-PM3PHYLO

Phylogenetic relationships among full-length *Pm3* homologs in rye, wheat and barley. The three main clades are shown in red, yellow and blue. The homologs corresponding to Pm3 and Pm8 are indicated. Bootstrap support values are given at nodes. Scale bar gives substitutions per site.

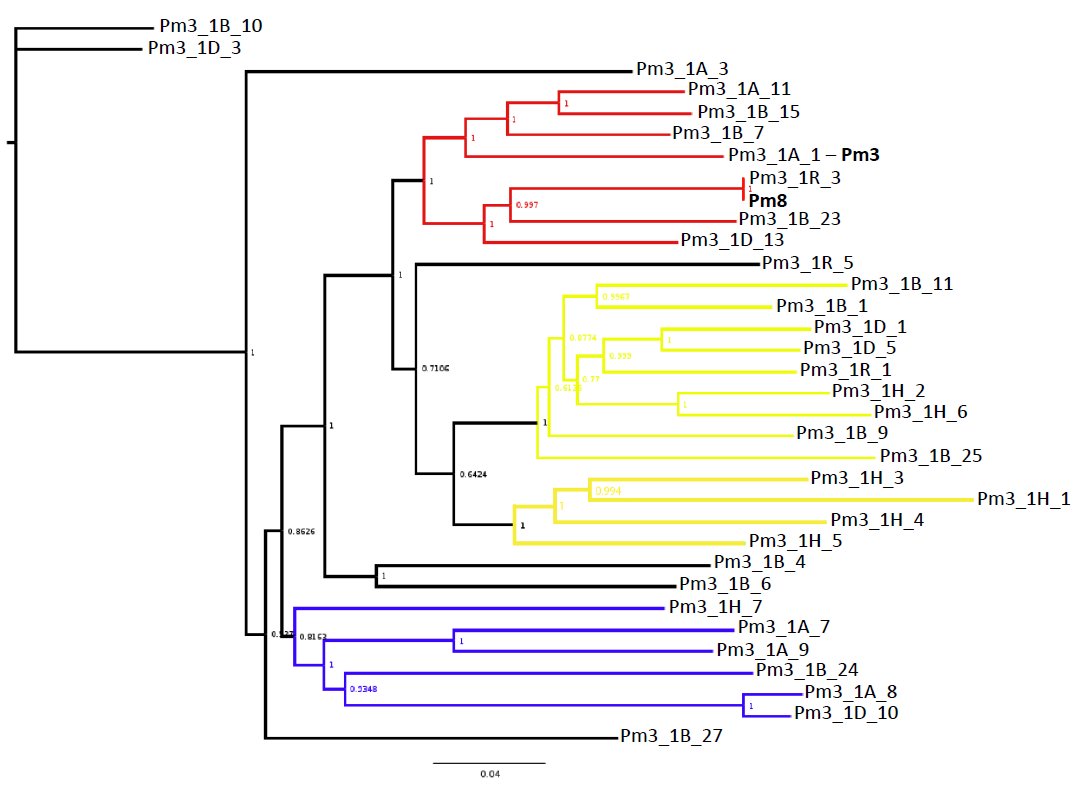

#### Supplementary Figure X.X. S-PM3COMP

Physical organisation of the *Pm3* gene cluster haplotypes in the (sub)genomes of cultivated Triticeae. Identifiers refer to S-PM3PHYLO with colours separating phylogenetic clades. Pseudogenes (i.e. those which contain in-frame stop codons and/or frameshifts) are marked with a psi symbol (Ѱ). A novel recombinant allele, *Pm17*, which falls outside the cluster and shares homology in one portion with rye *Pm8* is also diagrammed (tbl. S-NLR).

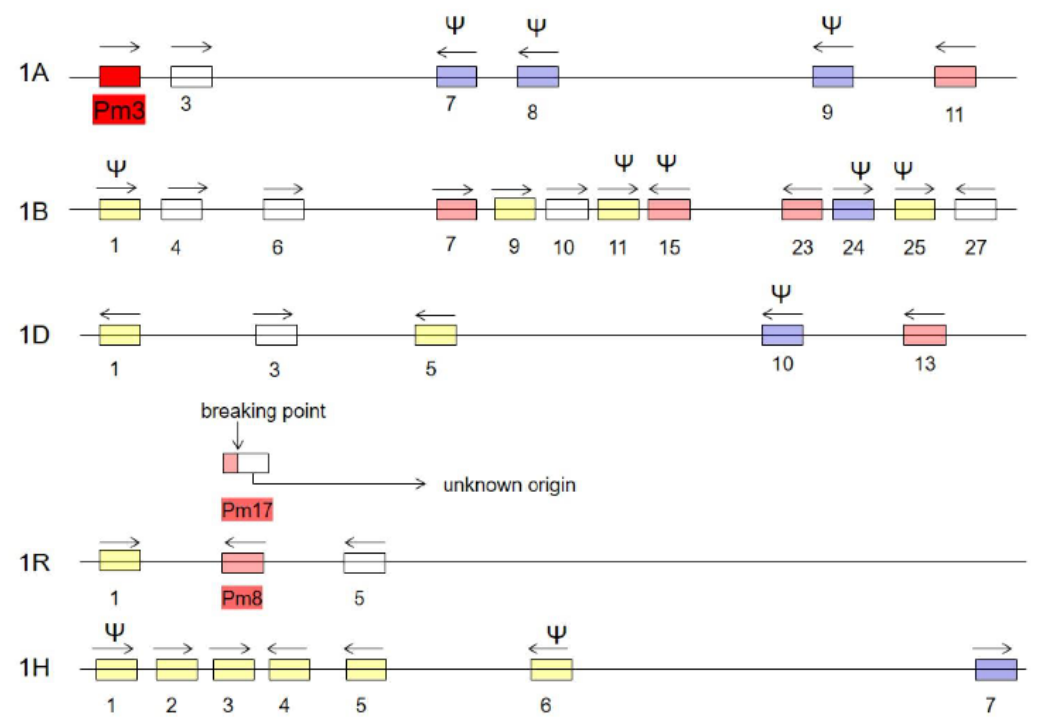

#### Supplementary Figure X.X. S-MLAPHYLO

Phylogenetic relationships among full-length *Mla* homologs identified in rye, wheat and barley (including 32 known *Mla* alleles, *TmMla1*, *Sr30* and *Sr50*). The two main clades are shown in red and blue. Homologs are indicated with black names and known alleles/genes with blue or red names. Bootstrap support values are given at nodes. Scale bar gives substitutions per site.

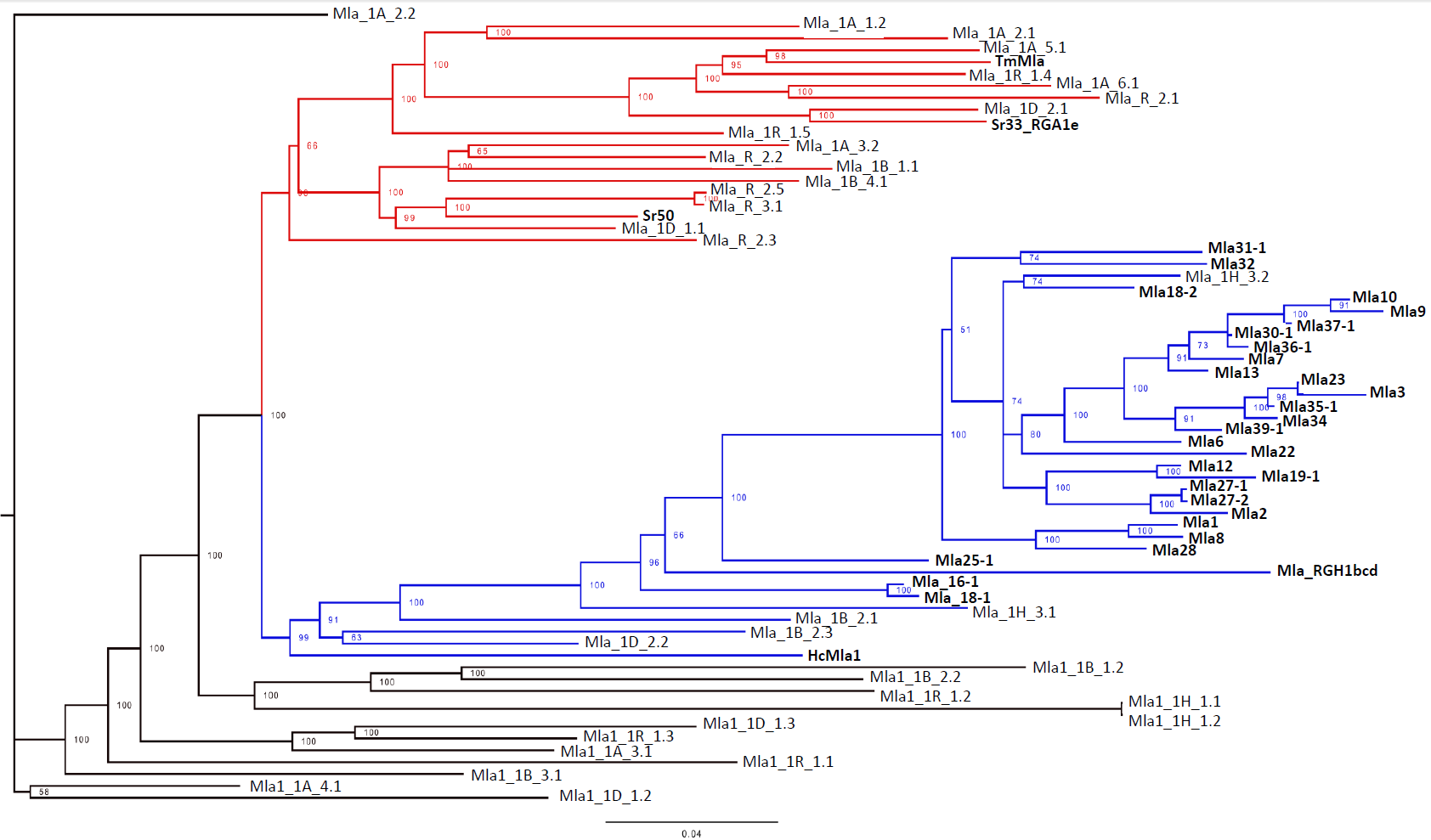

#### Supplementary Figure X.X. S-MLACOMP

Physical organisation of *Mla* cluster haplotypes across the (sub)genomes of cultivated Triticeae. Identifiers refer to S-MLAPHYLO and with colours separating phylogenetic clades.

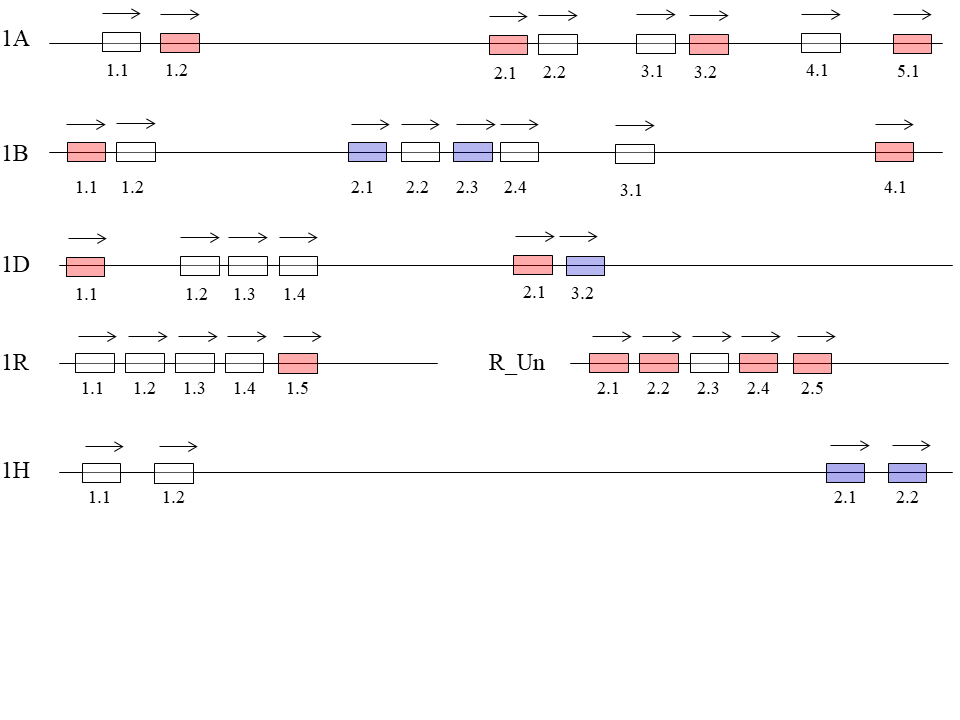

#### Supplementary Figure X.X. S-LR10CMP

Physical organization of *Lr10* haplotypes in wheat and rye. The first two haplotypes correspond to the described *Lr10* haplotypes H1 and H2. The third represents the situation in Chinese Spring, where the H2 haplotype is present. The fourth represents the situation in ‘Lo7’, where some rearrangements are observed compared to the H2 haplotype.

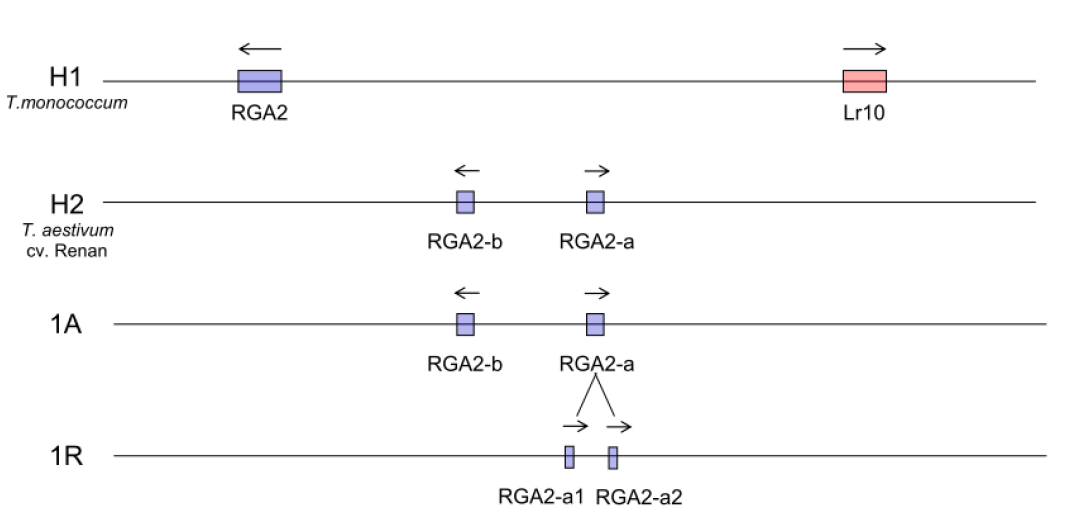

#### X.X Phylogenetic and homology-based investigations of low temperature tolerance gene homologs

To investigate the possible causes of low temperature tolerance (LTT) in rye, we conducted homology searches for the LTT-associated *Cbf* family genes, a cluster of which characterise the LTT locus *Fr2* in wheat^43-46^.

#### Supplementary Figure X.X. S-CBFPHYLO

Homology analysis of rye protein data shows that the *Fr2* region of ‘Lo7’ contains a cluster of 21 *Cbf* genes. Protein sequences for genes located between 614.3—616.5 Mbp on chromosome 5 were aligned by MUSCLE^47^ and a maximum-likelihood tree was constructed by MEGAX^48^. Each *Cbf* gene is named according to its best BLASTn^49^ match to previously named *Cbf* genes in ‘Lo7’^50^. A complete list of genes within the interval, including their positions and descriptors are provided (Supplemental Table S-FR2). The position of Group IV member *Cbf-14* (similar to wheat *TaCbf-A14*), whose transcript levels predict cold tolerance, is marked^51^. Bold arrows mark *Cbf* genes for which copy number variation between ‘Lo7’ and ‘Puma’ was detected (red=strong evidence; blue=moderate evidence). Genes from the interval that are not *Cbf* genes form an outgroup. The scale bar indicates substitution rate.

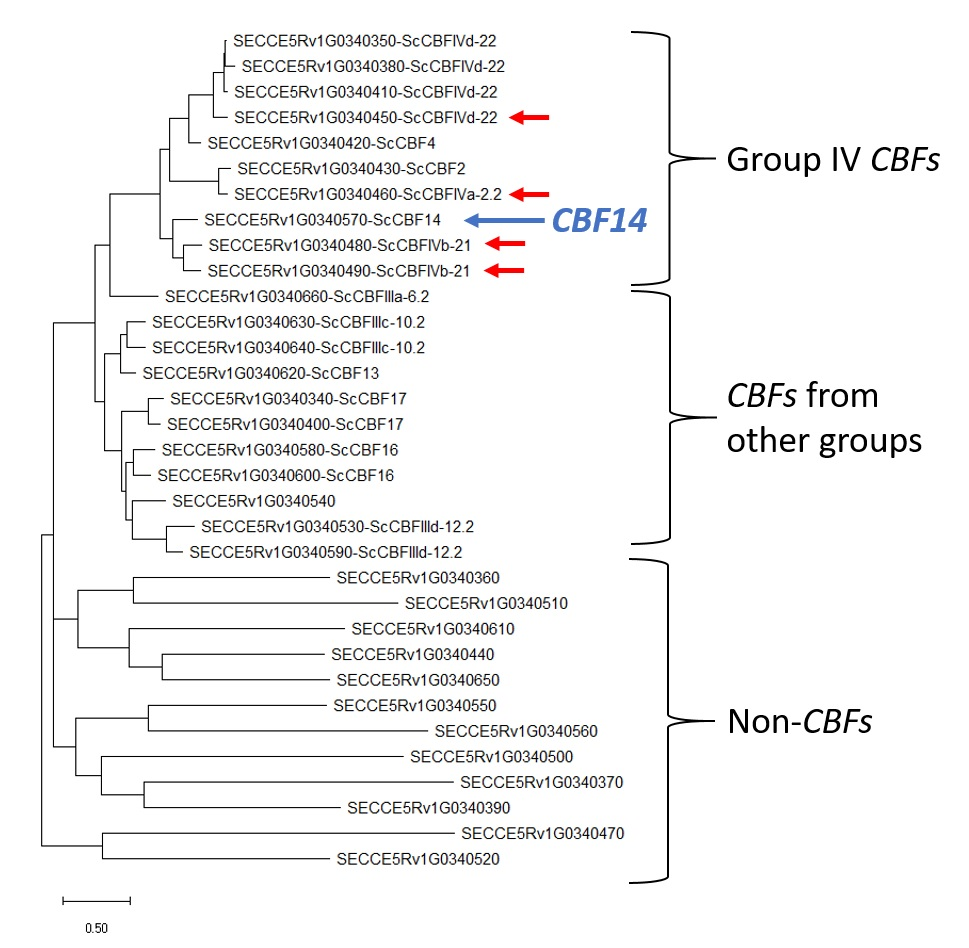

### Supplementary Note X. S-COLD: Experimental investigations of low temperature tolerance.

Investigations in LTT suggested a possible deletions in the *Vrn1* gene promoter, which was characterised by 10X Genomics sequencing (Methods; fig. S-VRN1DEL). Subsequent expression profiling experiments reinforced the hypothesis that attenuation of *Vrn1* expression may reduce LTT, but that the observed deletion alone is insufficient to guarantee *Vrn1* expression during cold acclimation (Methods; figs. S-LT50EXPT—S-VRN1EXPN).

Supplementary Figure X.X. S-VRN1DEL

Sequencing read depth of 10x Genomics Chromium reads in the gene *Vrn1* for ‘Lo7’, ‘Puma’ and the ‘Norstar5A:5R’ translocation line. The gene structure of *SECCE5Rv1G0353290*, which is on the ‘-’ strand, is indicated by vertical grey bars (top). A 597 bp deletion in the promoter of the gene in ‘Puma’ and ‘Norstar5A:5R’ was identified based on a reduction in read depth (red lines).

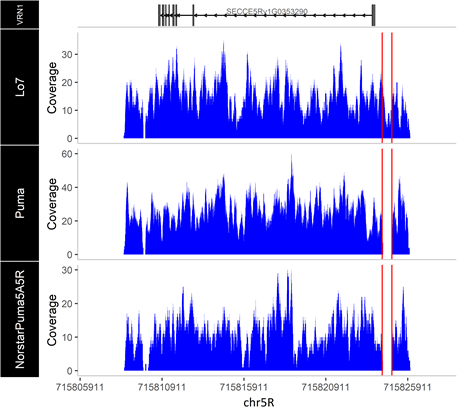

#### Supplementary Figure X.X. S-LT50EXPT

Experimental design and LT50 temperatures for ‘Norstar’ (dark blue), ‘NorstarPuma5A:5R’ (orange), ‘Puma’ (light blue), and the temperature sensitive control Winter Manitou (dark green). Crown temperature (black – left vertical axis) and day length (light green – right vertical axis) are indicated over the 70 day treatment (horizontal axis). Tissues isolated at each time point were also used for RNA sequencing analyses (Methods).

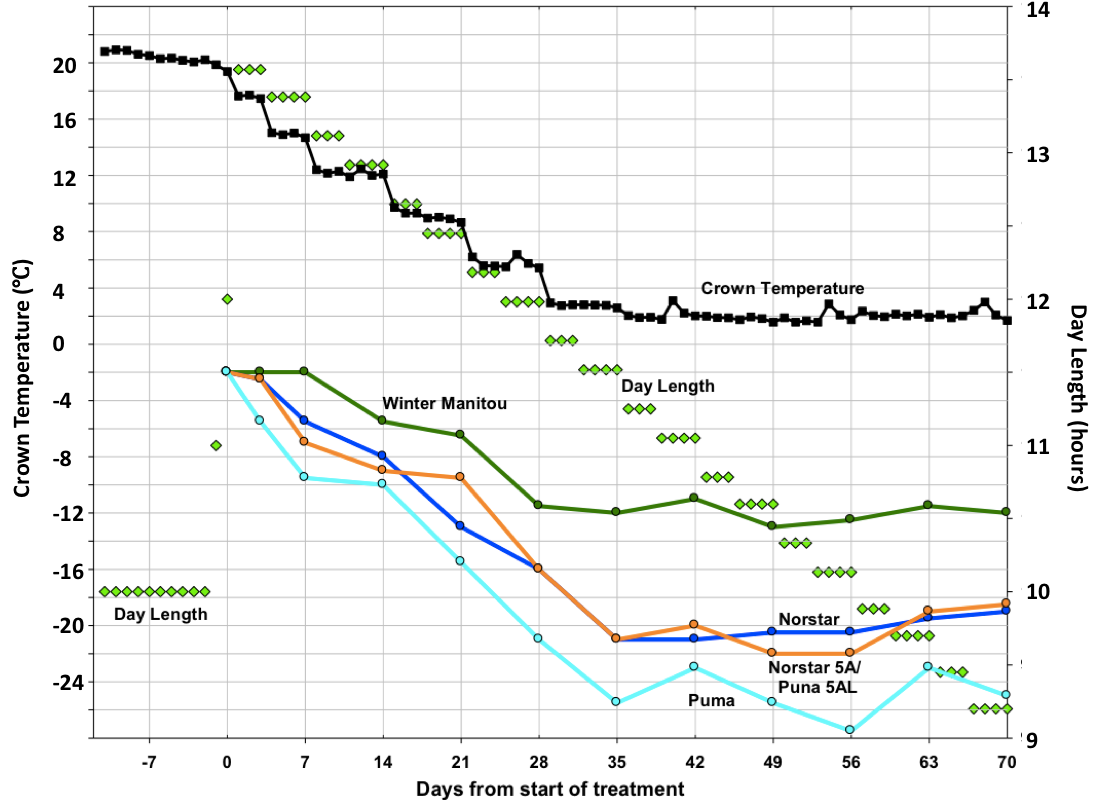

#### Supplementary Figure X.X. S-VRN1EXPN

Expression of the gene *Vrn1* in ‘Puma-SK’ (blue) compared to ‘NorstarPuma5A:5R’. Count reflects the normalised expression of the gene in response to reduced temperature and photoperiod over a 70 day time period (Methods).

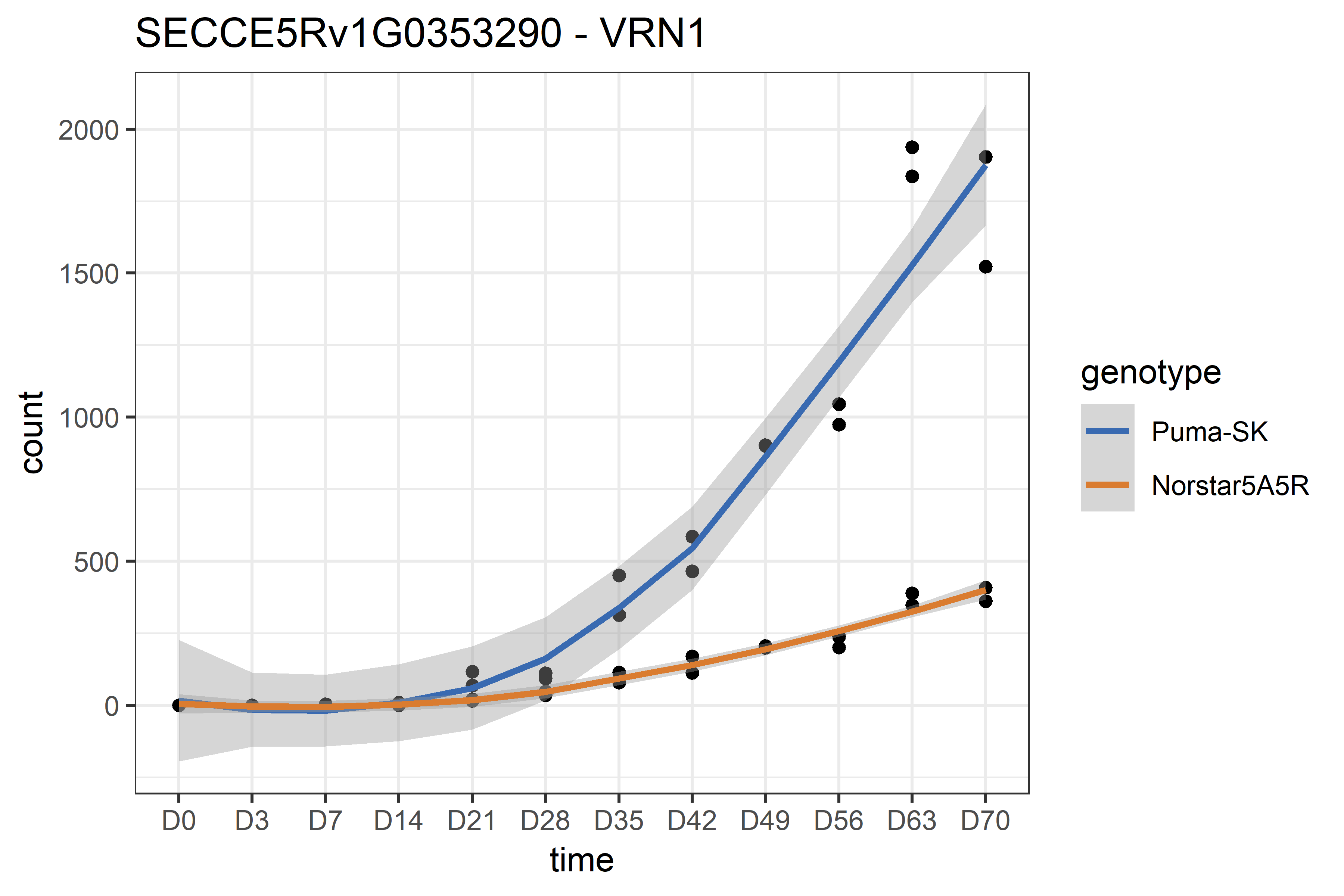

### Supplementary Note X. S-INTROG.

High throughput sequence data can be effectively utilised to characterise rye introgressions into wheat based on comparative mapping depths (Methods). This is easiest to appreciate by appraising a large number of different introgression haplotypes visually, which we provide here in figure S-CLASSCASES, to supplement the smaller number shown in main text figure M-INTROGb. By calling variants in the data, we were able to extract further information, for instance by characterising the chromatin origins in a novel wheat line in which a recombination has occurred between 1AL.1RS and 1BL.1RS introgressions (Methods; S-KSRECOMB). A genetic similarity matrix between 1RS introgressions also presented the opportunity to test the supposed common origins of the respective 1AL.1RS and 1BL.1RS introgressions (Methods; S-1RCOMMON).

#### Supplementary Figure X.X. S-CLASSCASES

A sample of genotypes from each of four bread wheat diversity panels displayed in groups according to the automated classification of rye chromatin introgressions. Row labels have the format [Classification]_[Panel]_[Sample ID]. Samples classified as “ambiguous” tend to have a moderate overabundance of 1RS reads, and underabundance of 1AL or 1BL, though far less pronounced than in the more confidently assigned samples. The causes of this ambiguity are currently unresolved. Sequencing depth does not appear to be a factor. Sample cross-contamination or barcode misassignment are possible but unlikely at this scale.

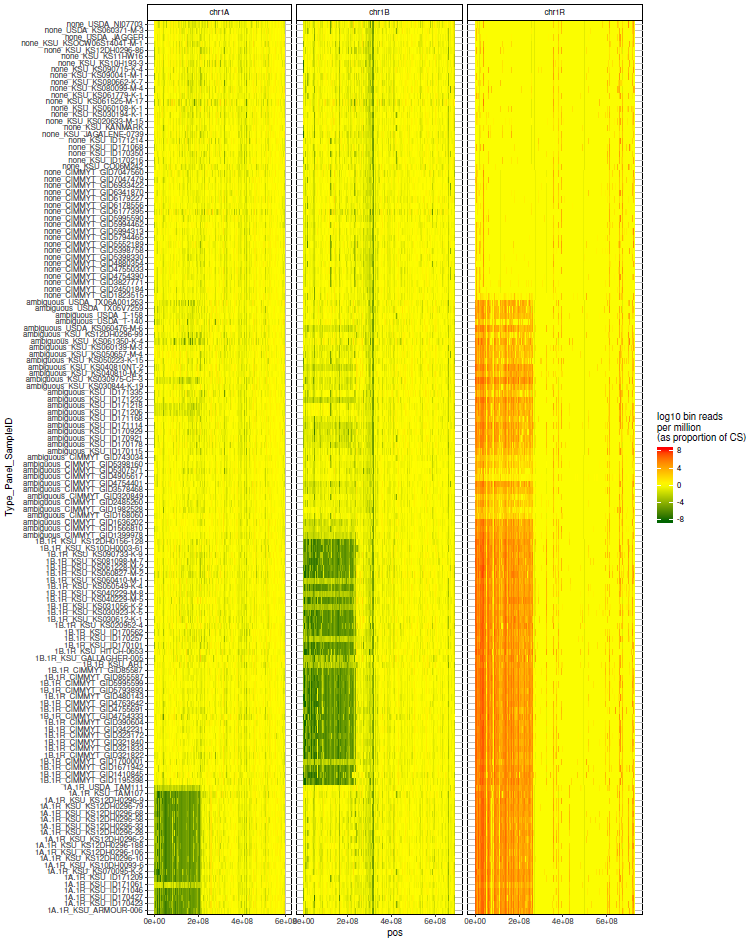

#### Supplementary Figure X.X. S-KSRECOMB

Characterisation of the recombination event between 1AL.1RS and 1BL.1RS introgressions. Chromosome 1R of the recombinant individual KS090616K was assigned to parental haplotypes Larry (carrier of 1BL.1RS introgression) and TAM112 (carrier of 1AL:1RS introgression).

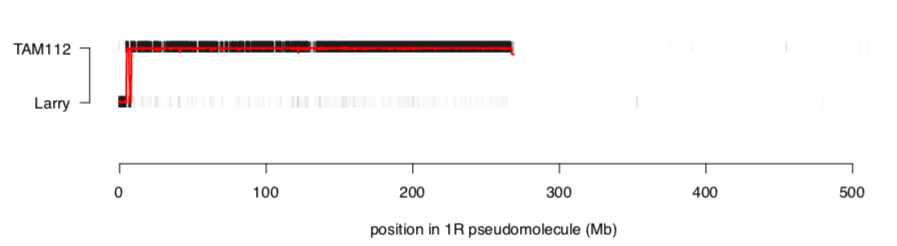

#### Supplementary Figure X.X. S-1RSCOMMON

Predicted 1RS carriers were selected to form a combined 1RS panel, in which SNPs were identified and pair-wise identity by state (IBS) percentages calculated. The square root values of percent different calls were used to derive a heatmap for all pair-wise comparisons. The heatmap dendrogram unambiguously shows two major clusters, consistent with common origins of each 1RS donor. The larger cluster corresponds to 1BL.1RS class, and the smaller cluster corresponds to 1AL.1RS class. Within 1BL.1RS class, sequencing technology appears to play a role, as exome sequenced lines show a subcluster slightly different from GBS lines.

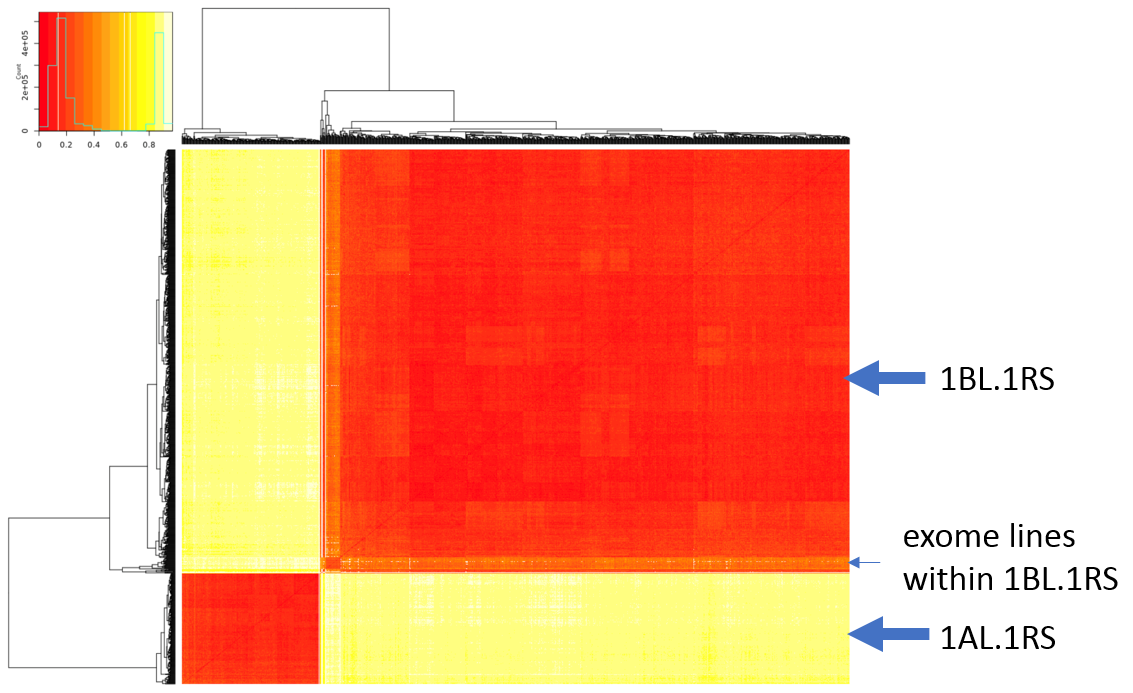

### Supplementary Note X. S-PUBLIC

A German federal agricultural trial with results relevant to the variable benefits of wheat-rye translocation lines under different climatic conditions is publicly reported in German (<https://www.bundessortenversuch.de/bsv-winterweizen>), but not published in the peer-reviewed literature; relevant findings are therefore compiled and summarised here in English.

#### Supplementary Figure X.X S-PUBLICTRIAL

The trial was conducted at 26 sites across Germany (upper left), during the exceptionally dry year 2018 (water balance for Summer and Spring shown upper middle and upper right; [www.dwd.de](http://www.dwd.de)), and compared the performance of released wheat varieties in yield and quality metrics among wheat lines and, included two with known 1RS.1AL (‘Asory’) and 1RS.1BL (‘Kamerad’) translocations. Trials were carried out in two intensity levels, with (blue columns) and without (green columns) growth regulator and fungicide treatment. Results from trials without growth regulator and fungicide treatment (intensity level 1) are basis for the description of ripening date, plant height, stem characteristics and susceptibility to diseases. Results from the intensity level 2 with growth regulator and fungicide treatment form the basis for the description of the 4 quality classes, A – bread wheat, B – milling, C – biscuit & feed wheat, E – premium quality bread wheat. Grain yield of individual genotypes refers the mean grain yield of three reference varieties (‘RGT Reform’, ‘Nordkap’ and ‘Elixer’), that is calculated for both intensities levels and set to 100%, indicated as a horizontal red line.

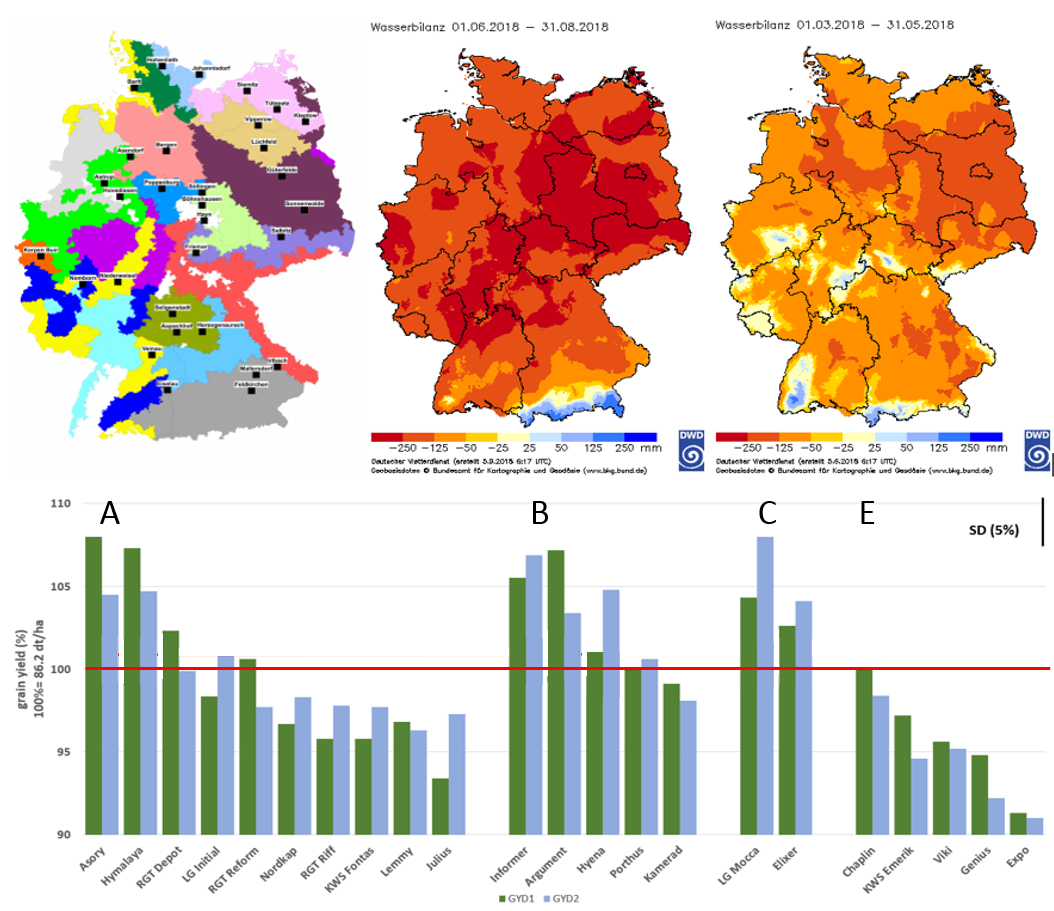
